# Design, synthesis, and characterization of [^18^F]mG2P026 as a high contrast PET imaging ligand for metabotropic glutamate receptor 2

**DOI:** 10.1101/2021.06.29.450249

**Authors:** Gengyang Yuan, Maeva Dhaynaut, Nicolas J. Guehl, Sepideh Afshar, Dalena Huynh, Sung-Hyun Moon, Suhasini Iyengar, Hye Jin Kang, Mary Jo Ondrechen, Georges El Fakhri, Marc D. Normandin, Anna-Liisa Brownell

## Abstract

An array of triazolopyridines based on JNJ-46356479 (**6**) were synthesized as potential PET imaging ligands for metabotropic glutamate receptor 2 (mGluR2) in the brain. The selected candidates **8**-**11** featured an enhanced positive allosteric modulator (PAM) activity (37-fold max.) and an apparent mGluR2 agonist activity (25-fold max.) compared to compound **6**. Radiolabeling of compounds **8** and **9** (also named mG2P026) was achieved via the Cu(I)-mediated radiofluorination in the automated TRACERLab^TM^ FXF-N platform. Both [^18^F]**8** and [^18^F]**9** were obtained with satisfactory radiochemical yields (> 5%, non-decay corrected), high molar activity (> 180 GBq/μmol), and excellent chemical and radiochemical purities (> 98%). Preliminary characterization of [^18^F]**8** and [^18^F]**9** in rats confirmed their excellent brain permeability with [^18^F]**9** showing better brain heterogeneity and favorable binding kinetics. Pretreatment with different classes of PAMs enhanced the radioactivity uptake for both [^18^F]**8** and [^18^F]**9** at the regions of interest by 20.3-40.9% and 16.7-81.6%, respectively, due to their pharmacological effects. Further evaluation of [^18^F]**9** in a nonhuman primate confirmed its superior brain heterogeneity in mapping mGlu2 receptors and its higher specific binding than [^18^F]**6**. Pretreatment with 0.5 mg/kg BINA led (2) to an enhanced brain uptake of [^18^F]**9** by 3% in high tracer uptake regions that was consistent with the rat studies. Therefore, [^18^F]**9** has the potential to be translated for human studies.

## INTRODUCTION

As the major excitatory neurotransmitter in the vertebrate central nervous system (CNS), glutamate is involved in numerous physiological and behavioral processes.^1, 2^ It modulates synaptic responses by activating two different classes of receptors, the ionotropic glutamate receptors (iGluRs) and the G-protein coupled metabotropic glutamate receptors (mGluRs).^3^ Among the mGluRs, the metabotropic glutamate receptor 2 (mGluR2) is highly enriched in the forebrain and predominantly localized on the presynaptic nerve terminals.^4, 5^ Activation of mGluR2 reduces the glutamate level and GABA release, thereby it is hypothesized as a therapeutic intervention toward the neurological diseases that involve increased glutamate transmission.^6, 7^ Activation of mGluR2 has shown efficacy in multiple preclinical animal models of schizophrenia,^8–12^ anxiety,^7^ and depression^13^. LY2140023 (**1**), the prodrug of mGluR2/3 agonist LY404039, achieved clinical validation in a phase II clinical trials in schizophrenic human subjects (Figure 1).^11^ However, it was discontinued due to the lack of efficacy in the following three phase 2 or phase 3 clinical trials.^14, 15^ Subsequent studies suggested mGluR2 but not mGluR3 mediated the antipsychotic effect of mGluR2/3 agonist.^16, 17^

To improve selectivity of mGluR2 binding over mGluR3, recent mGluR2 based drug discovery is leaning toward the development of agents that bind to the allosteric sites within the hydrophobic seven-transmembrane region (7-TM) instead of the highly conserved orthosteric glutamate Venus flytrap domain (VFTD).^18^ Besides the enhanced binding selectivity, these ligands feature enhanced brain permeability and reduced liability of receptor desensitization than that of the amino acid-based orthosteric ligands.^19–21^ Numerous structurally diverse mGluR2 positive allosteric modulators (PAMs) have been reported, for example, BINA (**2**),^22^ AZD8529 (**3**)^23^ and JNJ-40411813^24^ (**4**, Figure 1). Compound **3** entered phase II clinical trials for schizophrenic patients in 2009 but was halted in 2011 because of lack of efficacy.^25^ Compound **4** advanced to a phase IIa study in schizophrenia, but its efficacy was not ideal.^26, 27^ Scaffold-hopping of the pyridone core of compound **4** led to a series of 1,2,4-triazolopyridines as potent and selective mGluR2 PAMs with improved drug properties.^28–30^ In this class of PAMs, [^11^C]JNJ42491293 ([^11^C]**5**, Figure 1) was the first disclosed positron emission tomography (PET) radioligand that was characterized in human.^31^ Despite the satisfactory in vitro, ex vivo and in vivo imaging results in rats, [^11^C]**5** showed apparent off target binding in human myocardium, precluding it as an useful imaging tool for mGluR2.^31^ As a noninvasive imaging technique, PET would enable quantification of the biodistribution, expression, and modulation of a protein target under normal and disease conditions. We have therefore developed [^18^F]JNJ-46356479 ([^18^F]**6**)^32, 33^ and [^11^C]mG2P001 ([^11^C]**7**)^34–36^ as alternative PET imaging tools (Figure 1). Compound **6** belongs to the 1,2,4-triazolopyridines^28^ and has been employed as an mGluR2 selective PAM to study the binding selectivity of [^11^C]**5**.^31^ Our studies have confirmed that both [^18^F]**6** and [^11^C]**7** are suitable ligands for imaging mGluR2 in the rodent and non-human primate brain.^32, 34, 36^ Meanwhile, we also found the apparent white matter binding of [^18^F]**6** at the later stage of the dynamic scan (60-120 min) in the primate studies.^32^ Although mGluR2 is known to bind to white matter,^37^ we concluded part of the binding was off target.^32^

To enhance the *in vivo* binding specificity of [^18^F]**6**, we intend to improve its binding affinity and optimize its physicochemical properties. Although a vast chemical space of the 1,2,4-triazolopyridines has been covered by patent applications and research articles,^28–30, 38^ their use as PET imaging ligands has not been explored. Herein, we focused on four close analogues of compound **6** to fine-tune its binding and physiochemical properties in compounds **8**-**11**. Moreover, we explored the potential of modifying the core scaffold of **6** in producing new classes of mGluR2 PAMs as described for compounds **12** and **13**. In addition to the mGluR2 functional affinity, the synthesized ligands were also characterized for their binding toward other mGluRs, especially their agonist activity against mGluR2/3. Furthermore, molecular docking was used to probe the binding interactions at the allosteric site. Once labeled with fluorine-18, radioligands [^18^F]**8** and [^18^F]**9** were evaluated by PET imaging studies in rats and/or a non-human primate to examine their feasibility as an *in vivo* imaging tool for mGluR2.

**Figure 1.**
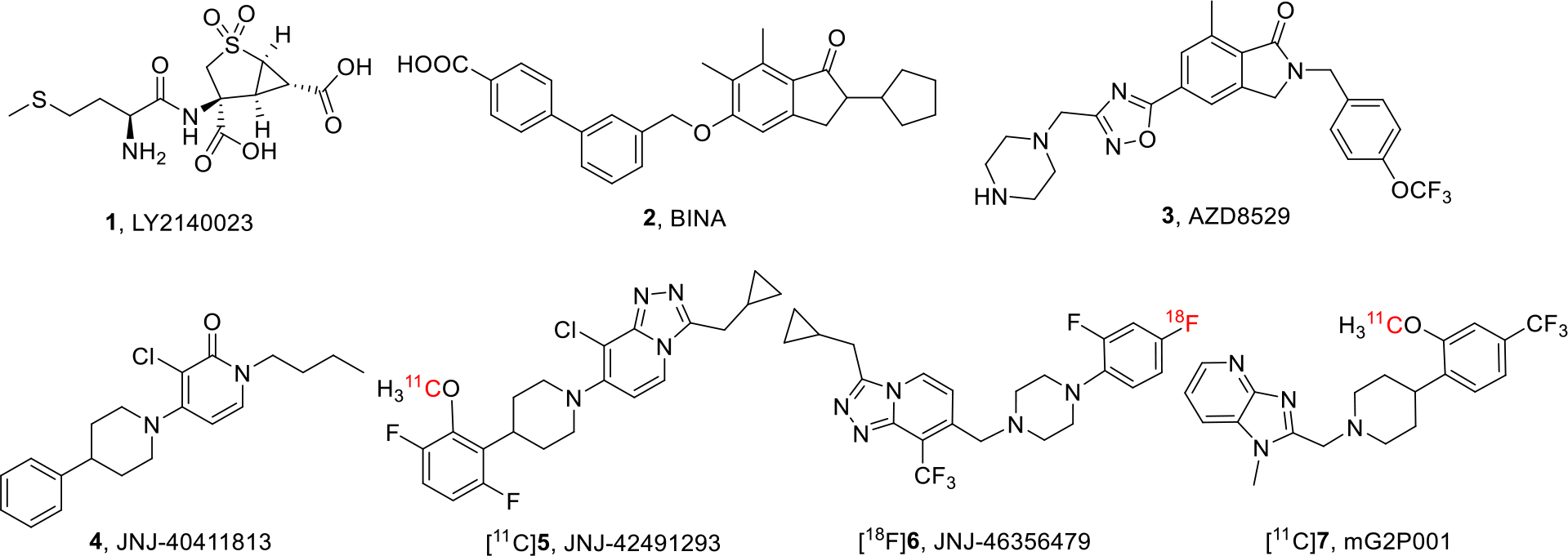
Structures of typical mGluR2 PAMs and PET radioligands

## RESULTS AND DISCUSSION

### Chemistry

We aimed to synthesize PET radioligands with favorable physiochemical properties, enhanced binding profiles and ease to incorporate fluorine-18. As scheme 1 shows, the structural variations based on **6** were focused on 1) replacing the piperazine linker with the less basic piperidine in compound **8**, 2) fine-tuning the fluorine substituents at the tail arene of compounds **9-11**, and 3) modifying the core scaffold to give compounds **12-13**. Compound **12** is a Regio-analogue of **6**, where the position of aryl-CF_3_ and cyclopropyl methylene groups on the 1,2,4-triazolopyridine were changed. Compound **13** changes to a pyrazolopyrimidine analogue by altering the nitrogen atoms and substituents on the 1,2,4-triazolopyridine core. The amide group of **13** was not further reduced to maintain a lower lipophilicity (cLogP = 3.89, Table 1).

Syntheses of compounds **8**-**11** were achieved by the same reductive amination conditions as compound **6**.^28, 32^ The coupling reactions occurred between aldehyde **14** and the corresponding amines **15**-**18** as shown in Scheme 1. Compound **14** was prepared via the literature method.^28^ Compound **12** was obtained in a similar manner as **6** but using 2,3-dichloro-5-(trifluoromethyl)pyridine (**19**) as a starting material. The 8-chlorotriazolopyridine **22** was obtained from **19** following a sequential reactions of chlorine displacement, acylation and cyclodehydration. The aldehyde intermediate **24** was prepared from **22** via a Suzuki coupling to incorporate the vinyl group and a subsequent oxidative reaction with osmium tetroxide. A final reductive amination between **24** and 1-(2,4-difluorophenyl)piperazine led to compound **12**. Synthesis of compound **13** began with the condensation reaction between compounds **25** and **26** to get compound **27**.^39^ Compound **27** was then subjected to oxidative cyclization, halogen exchange with CF_3_ source, and hydrolysis to give intermediate **30**. The final coupling reaction between **30** and the 1-(2,4-difluorophenyl)piperazine led to compound **13**.

**Scheme 1.**
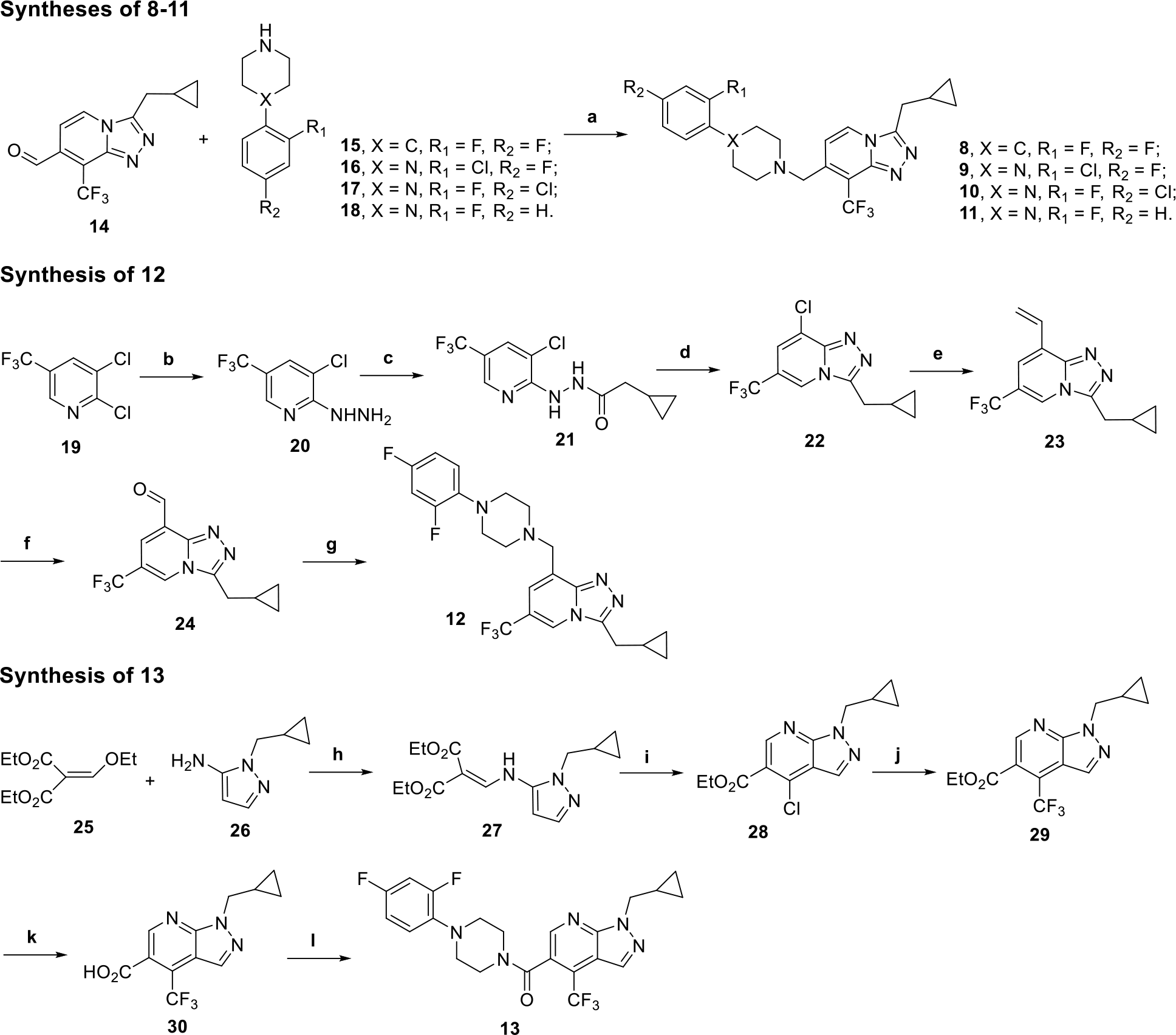
Syntheses of compounds **8**-**13**. Reagents and conditions: (a) TEA, MgSO_4_, NaBH(OAc)_3_, rt, 12 h; (b) NH_2_NH_2_, 1,4-dioxane, 80 °C, 16 h; (c) CyPrCOCl, Et_3_N, CH_2_Cl_2_, rt, 16 h; (d) POCl_3_, 1,2-dichloroethane, 150 °C, 10 min, microwave; (e) 4,4,5,5-tetramethyl-2-vinyl-1,3,2-dioxoborolane, Pd(PPh3)3, 1,4-dioxane, sodium bicarbonate, 150 °C, 15 min, microwave; (f) NaIO_4_, OsO_4_, 1,4-dioxane/H_2_O, rt, 2 h; (g) 1-(2,4-difluorophenyl)piperazine, TEA, MgSO_4_, NaBH(OAc)_3_, rt, 12 h; (h) EtOH, reflux, 16 h; (i) POCl_3_, reflux, 16 h; (j) i. NaI, acetyl chloride, CH_3_CN, 50 °C, 4 h; ii. CuI, methyl 2,2-difluoro-2-(fluorosulfonyl)acetate, DMF, 100 °C, 6 h; (k) LiOH, THF/H_2_O, rt, 12 h; (l) 1-(2,4-difluorophenyl)piperazine, HATU, DIPEA, DMF, rt, 16 h.

### In vitro properties

Compounds **8-13** were characterized for their functional affinity as mGluR2 PAMs and their binding selectivity against mGluR1-6 and 8 following our previous methods.^32, 34^ As shown in Figures 2a-c and Table 1, compound **8** showed an enhanced PAM activity than that of compound **6** in our cAMP biosensor assay (EC_50_, 20 nM versus 166 nM). By screening its mGluR2/3 agonist activity, compound **8** was also found to have a more potent mGluR2 agonist activity (IC_50_ = 136 nM) but a similar mGluR3 agonist activity (IC_50_ = 3 µM) than compound **6** (mGluR2, IC_50_ = 2 µM; mGluR3, IC_50_ = 9 µM). Modifications at the tail aryl group in compounds **9-11** led to further enhanced mGluR2 PAM activities with compound **11** being the most potent PAM (EC_50_ = 4.4 nM). Compounds 9-11 also had apparent mGluR2 agonist activity, ranging from 82-371 nM, but weak mGluR3 agonist activity. Therefore, compounds **8-11** can be categorized as mGluR2 ago-PAMs. A further analysis of the most potent PAM **11** revealed that it had an allosteric binding affinity of pKB 5.83, and it enhanced glutamate binding affinity by 8.5-fold and increased glutamate efficacy by 12.28-fold with an agonist efficacy parameter τB of 17.05 (Figure 2d). On the other hand, the Regio-analogue **12** had a significantly decreased mGluR2 PAM activity (EC_50_ = 913 nM), while the pyrazolopyrimidine **13** failed to retain this functional affinity (EC_50_ = 7.3 µM). Besides, compounds **8-13** showed no obvious binding toward other mGluRs (Supporting Information Table S1). Additionally, although compounds **6** and **8-13** have relatively high cLogP values, their experimentally determined LogD_7.4_ values were in the range of 1.0-4.0 for CNS penetrants (Table 1).^40, 41^

**Figure 2.**
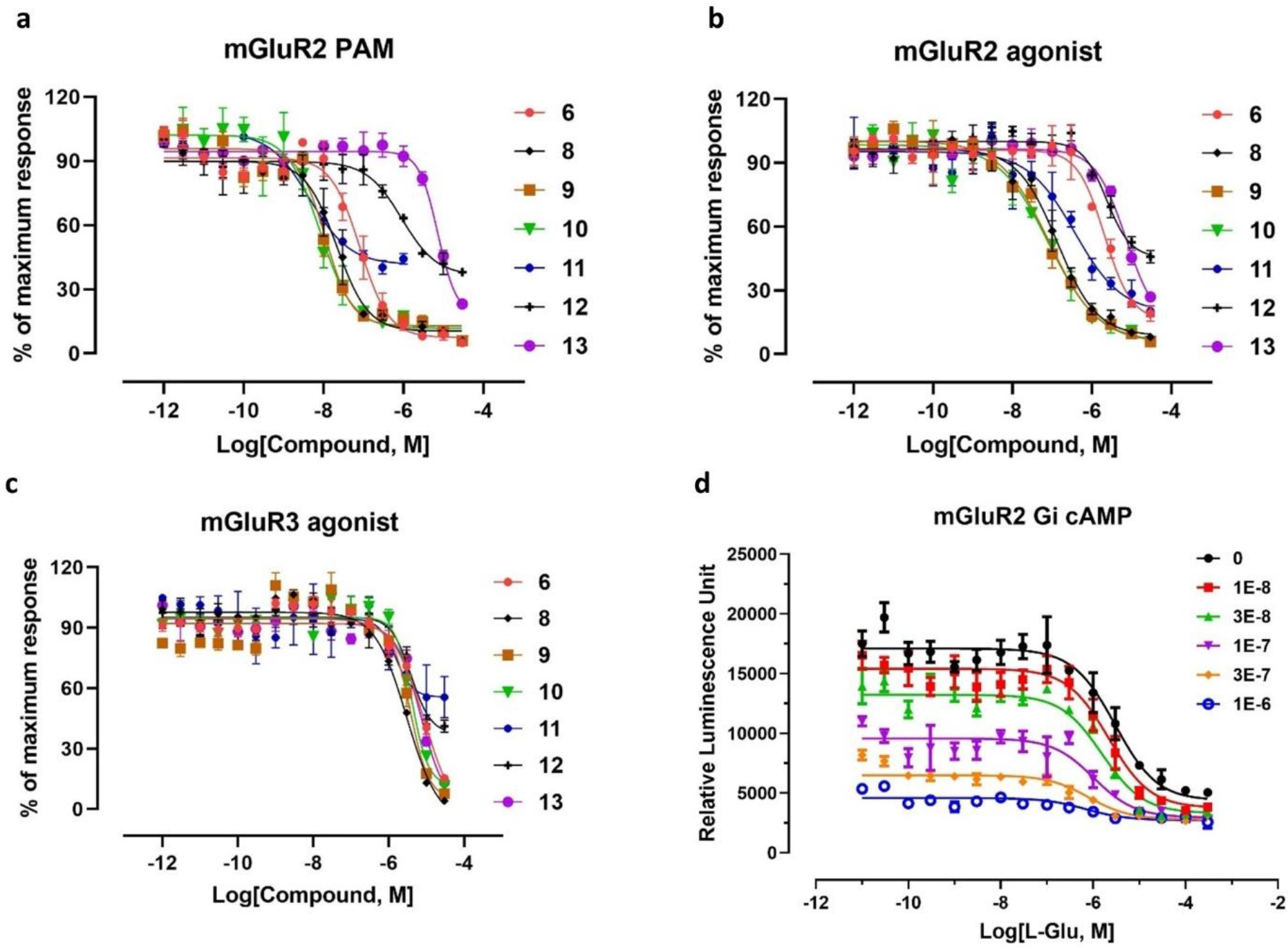
The binding properties of compounds **8**-**13**. (**a**) Binding curves of compounds **8**-**13** as mGluR2 PAMs; (**b**) compounds **8**-**13** binding at the orthosteric sites of mGluR2 and (**c**) mGluR3 receptors. The relative luminescence unit in the Y axis in **a**, **b** and **c** were normalized to their maximum response values; (**d**) characterization of compound **11** as an mGluR2 ago-PAM.

As reported in literature, there are probe-dependent effects for different classes of mGluR2 PAMs regarding to their mode of actions.^42–44^ Moreover, it was found that mGluR2 agonist could affect the binding event of a tritiated PAM radioligand by increasing the radioligand binding affinity toward mGluR2^44, 45^ and/or increasing the mGluR2 receptor density^46^ *in vitro*. As described in our previously work, compound 7 could dose-dependently enhance [^11^C]**7** radioactivity uptake in both rodents and monkeys, though it showed no apparent mGluR2/3 agonist activity (IC_50_ > 10 µM) in the same assays.^34, 36^ Therefore, the added feature of potent mGluR2 agonist activity in compounds **8-11** might result in unique features in characterizing the PET radioligands *in vivo*.^42, 43^

**Table 1.**
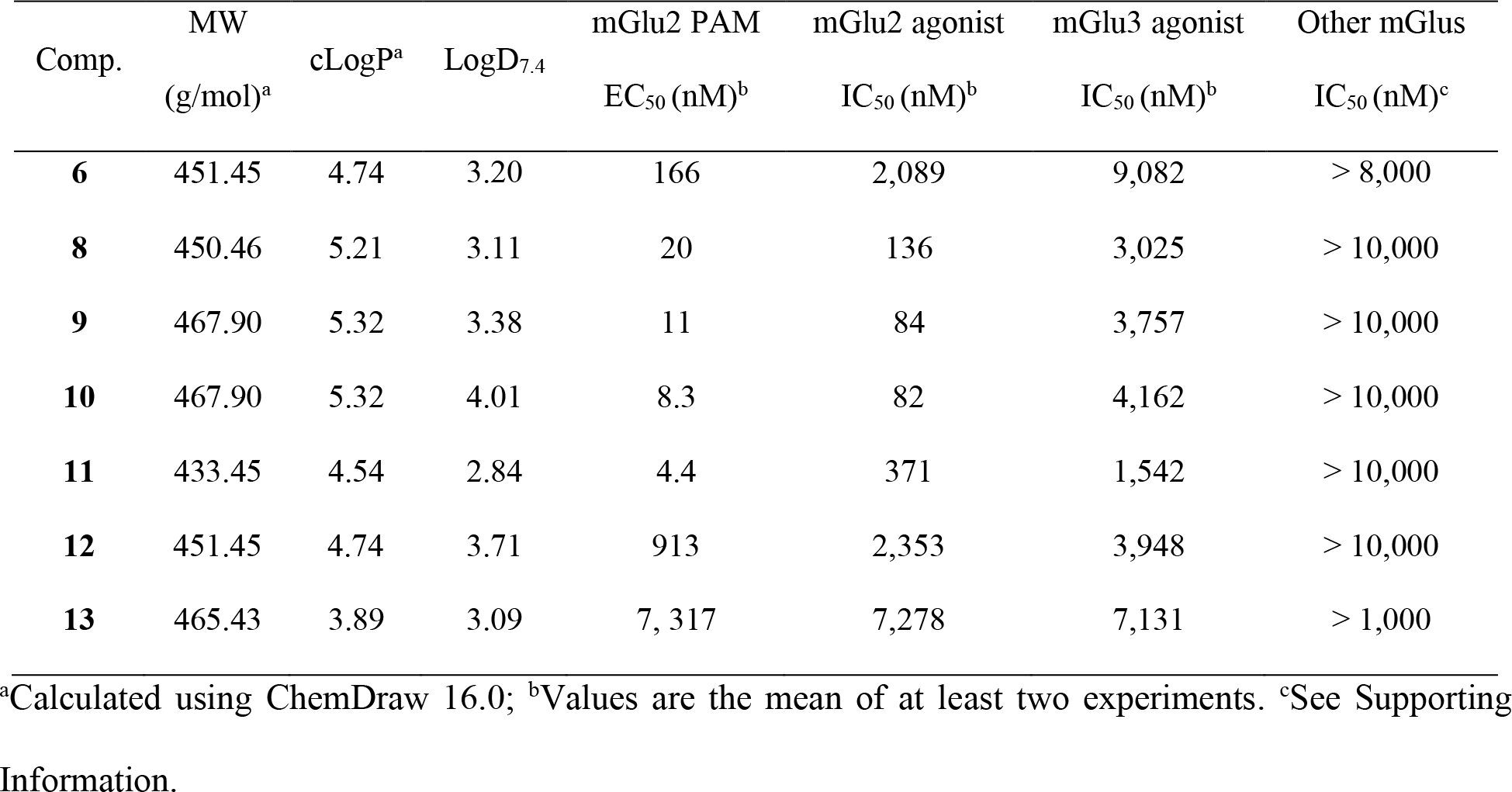
*In vitro* properties of compound **6** and its analogues **8**-**13**

### Structural Insights of the compounds 8-11

Compounds **8-11** were docked into our previously described mGluR2 homology model^34^ and their ligand-protein binding interactions were compared with that of compound **6**. As shown in Figure 3a, compounds **6** has its heterocyclic core buried at the bottom of the allosteric binding site, which is surrounded by interacting residues of F623, F643, Y647, L732, R735, V736, L739, W773 and V798. The 1,2,4-triazolopyridine core of **6** forms π-π stacking interactions with residues of F623, F643, and W773. Compounds **8-11** occupy the same binding site and adopt similar binding poses as that of compound **6**. An overlaid binding image of **6** and **8-11** is shown in Figure 3b. The docking scores for compounds **6** and **8-11** were more than 10.0 kcal/mol, indicating their potent binding toward mGluR2. Although functional affinities are not necessarily consistent with binding affinities,^47, 48^ the predicted binding modes provide insights to the ligand-protein binding events. Detailed binding mode for each ligand and the corresponding docking score are included in the supporting information (Figure S1 and Table S2).

**Figure 3.**
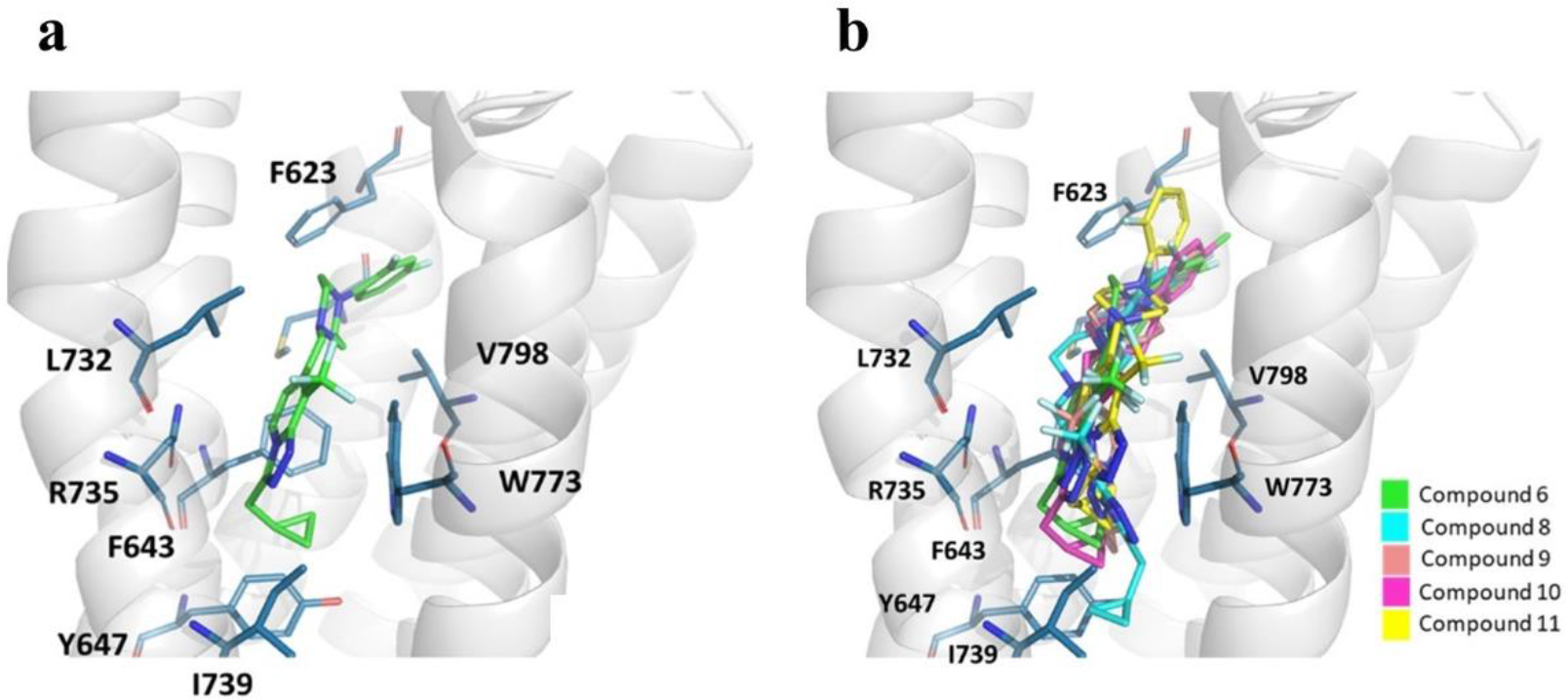
Molecular docking results of **6**, and **8**-**11**. (**a**) Binding poses of compound **6**; (**b**) Binding poses of **8**-**11** overlaid with **6** at the allosteric site. Images are rendered in Pymol 2.3.3.

### Radiochemistry

Compounds **8** and **9** were selected for radiolabeling with fluorine-18 due to their improved PAM affinities than that of compound **6**. Moreover, the *para*-fluoride in compounds **8** and **9** provides less steric hinderance when introducing bulky leaving group and/or during S_N_Ar replacement with [^18^F]F^-^ ion than the *ortho*-fluoride at compounds **10** and **11**. Both [^18^F]**8** and [^18^F]**9** were synthesized by the previously described method using our modified alcohol-enhanced Cu-mediated radiofluorination of organoboranes as [^18^F]**6** (Scheme **2**).^32, 49^ The boronic acid pinacol esters (Bpin) **37** and **38** were prepared via the reductive amination between **14** and the corresponding fragments **35** and **36**, respectively. Fragment **35** was prepared over **3** steps. The intermediate **34** was obtained via the Suzuki coupling between compounds **31** and **32** and the subsequent Miyaura borylation of **33**.^50^ Compound **34** was then hydrogenated and deprotected to give compound **35**. The order of the synthetic steps was important. Otherwise, if compound **33** was hydrogenated first using Pd/C (10 wt. %), the bromide group would be cleaved under the hydrogenation conditions. Alternatively, switching to PtO_2_ as a catalyst seems to have avoided this issue according to a recent publication.^51^ The automated synthetic procedures used for [^18^F]6^49^ were applied for the syntheses of both [^18^F]**8** and [^18^F]**9** in the TRACERLab^TM^ FXF-N platform. In the reaction, tetraethyl ammonium bicarbonate (TEAB) was used as a base and phase transfer agent, *n*-BuOH/dimethylacetamide (DMA) were used as solvents and [Cu(OTf)2py4] was used as a catalyst. The same stoichiometry for each reagent was applied. [^18^F]**8** was isolated with a radiochemical yield (RCY) of **8** ± 2% (non-decay-corrected, n = 5) and a molar activity (A_m_) of 273 ± 124 GBq/μmol (n = 2) at end of synthesis (EOS, t = 55 min). [^18^F]**9** was prepared with a RCY of 5 ± 3% (non-decay-corrected, n = 6) and a A_m_ of 348 ± 144 GBq/μmol (n = 3) at end of synthesis (t = 55 min). Both radiotracers were formulated into 10% ethanolic saline solution with excellent chemical and radiochemical purities (> 98%) at the time of injection.

**Scheme 2.**
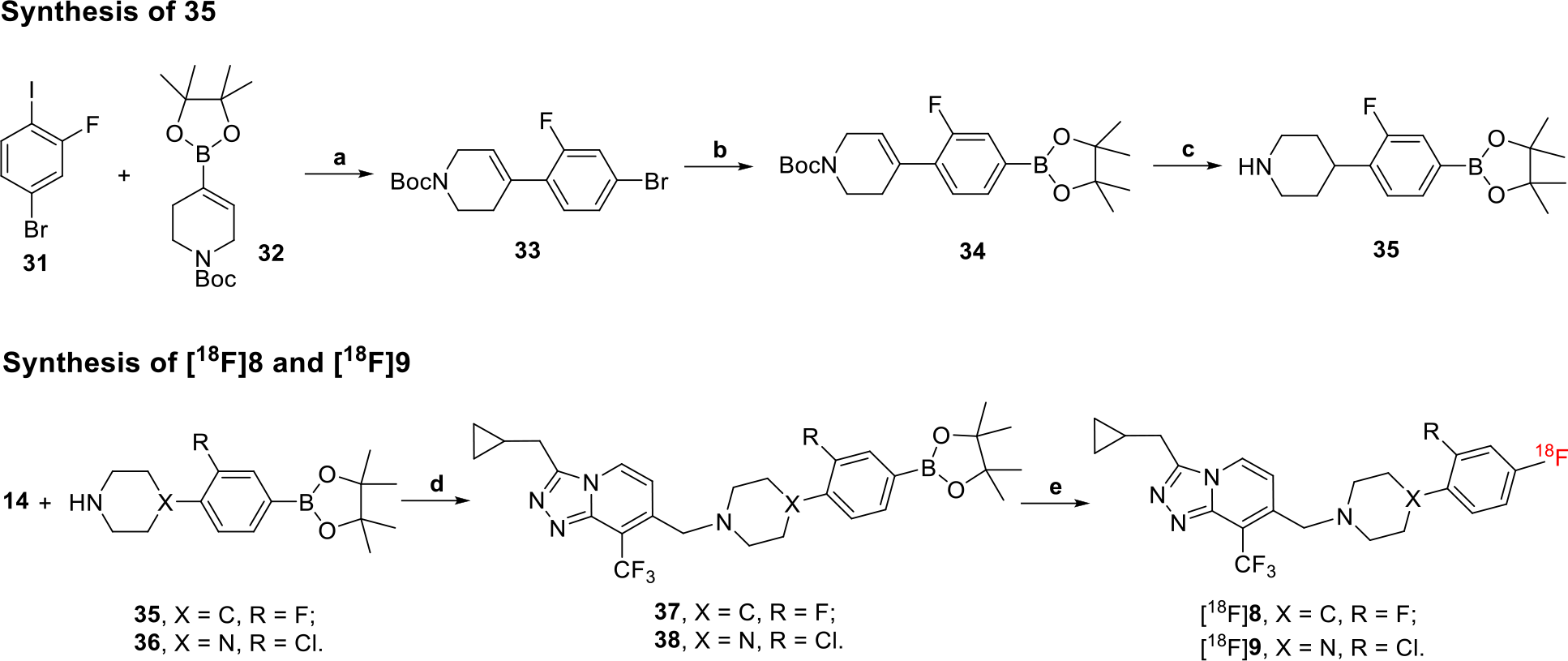
Syntheses of [^18^F]**8** and [^18^F]**9**. Reagents and conditions: (**a**) Na_2_CO_3_, Pd(PPh_3_)_2_Cl_2_, dimethyl ether/water (5:1, v/v), 85°C, 2 h; (**b**) KOAc, Pd(PPh3)2Cl2, 1,4-dioxane, 80°C, overnight; (**c**) i. H_2_, Pd/C (10 wt. %), 43 psi, rt, overnight; ii. 4N HCl in dioxane, rt, 1 h; (**d**) TEA, MgSO_4_, NaBH(OAc)_3_, rt, 12 h; (**e**) **37** or **38** (6.2 µmol), TEAB (14.1 µmol), [Cu(OTf)_2_py_4_] (13.3 µmol), DMA (0.8 mL), *n*-Butanol (0.4 mL), 130 °C, 10 min.

### PET imaging in Rats

The radioligands [^18^F]**8** and [^18^F]**9** were at first characterized in rat models (male, Sprague-Dawley). In each study, rats were anesthetized and 60-min dynamic PET scans were performed following the tail vein injection of radiotracer. Blocking agents were administered 10 min before radioactivity. Representative PET images of cumulative volumetric distribution of [^18^F]**8** and [^18^F]**9** are shown in Figure 4a. The summed images at time interval of 2-20 min are displayed on coronal, axial and sagittal levels. Both tracers were brain penetrant and clearly outlined the biodistribution of mGlu2 receptors in regions of striatum, thalamus, cortex, hippocampus and cerebellum. Time-activity curves (TACs) of [^18^F]**8** and [^18^F]**9** from their representative baseline studies showed rapid brain uptake and time-dependent radioactivity accumulation in different brain regions (Figure 4b). [^18^F]**8** peaked at 5 min with a SUV_max_ of 1.9 and the accumulations were similar across the region of interests. On the other hand, [^18^F]**9** had a SUV_max_ value of 1.7 at 5 min and the accumulations were similar in striatum, cerebellum, cortex and hippocampus, but was relatively lower in thalamus. The binding kinetics of [^18^F]**9** is more favorable for diagnostic purposes than [^18^F]**8** since it shows faster washout from all investigated brain areas than [^18^F]**8** resulting to lower radiation dose response. The estimated remaining radioactivity 60 min after the injection was 37% of the maximum binding using [^18^F]**9** while it was 58% using [^18^F]8.

To test the *in vivo* tracer binding selectivity and the pharmacological effects of mGluR2 PAMs toward radioligand binding, blocking experiments were carried out for [^18^F]**8** via self-blocking and for [^18^F]**9** via different classes of mGluR2 PAMs, including compounds **2, 7** and **9**, at different doses. As shown in Figure 4c, all the blocking experiments resulted in an enhanced radioactivity uptake in the brain regions of interest at the time window of 2-20 min. Self-blocking of [^18^F]**8** at a dose of 4 mg/kg increased its uptake by 20.3-40.9% and significant enhancement was achieved in all cortices, hippocampus, and cerebellum. On the other hand, [^18^F]**9** had the most elevated radioactivity uptake from the self-blocking studies (51.1-81.6%). No dose dependency was observed for self-blocking at the current doses of 2.0 mg/kg versus 4.0 mg/kg. Compound **7** elicited similar potentiation effects at a dose of 4.0 mg/kg compared to the self-blocking, ranging from 45.1-63.1% and significant enhancement was observed in all the investigated brain areas. This is consistent with our previous finding that compound 7 was able to significantly enhance the brain uptake of [^11^C]**7** in both rats and non-human primates.^36^ Moreover, although compound 7 reduced the brain uptake of [^18^F]**6** in rat brain by 11.6-18.8%, it enhanced the radioactivity uptake of [^18^F]**6** in a non-human primate by an averaged 19.4% when quantified by the regional total volume of distribution (*V_T_*) estimates. Pre-administration of BINA (**2**) resulted in a significantly enhanced binding of [^18^F]**9** in the thalamus, cingulate and prefrontal cortices, hippocampus, and cerebellum at a dose of 2 mg/kg, ranging from 16.7% to 29.3%. Whereas pretreatment of BINA at a lower dose of 1 mg/kg gave a negligible radioactivity enhancement at the brain areas investigated.

These results confirm that both [^18^F]**8** and [^18^F]**9** are suitable radioligands for imaging mGluR2 in the rat brain and their brain uptake can be enhanced by the pretreatments of mGluR2 PAMs. For diagnostic purpose, [^18^F]**9** has a better brain heterogeneity and more favorable binding kinetics than [^18^F]8.

**Figure 4.**
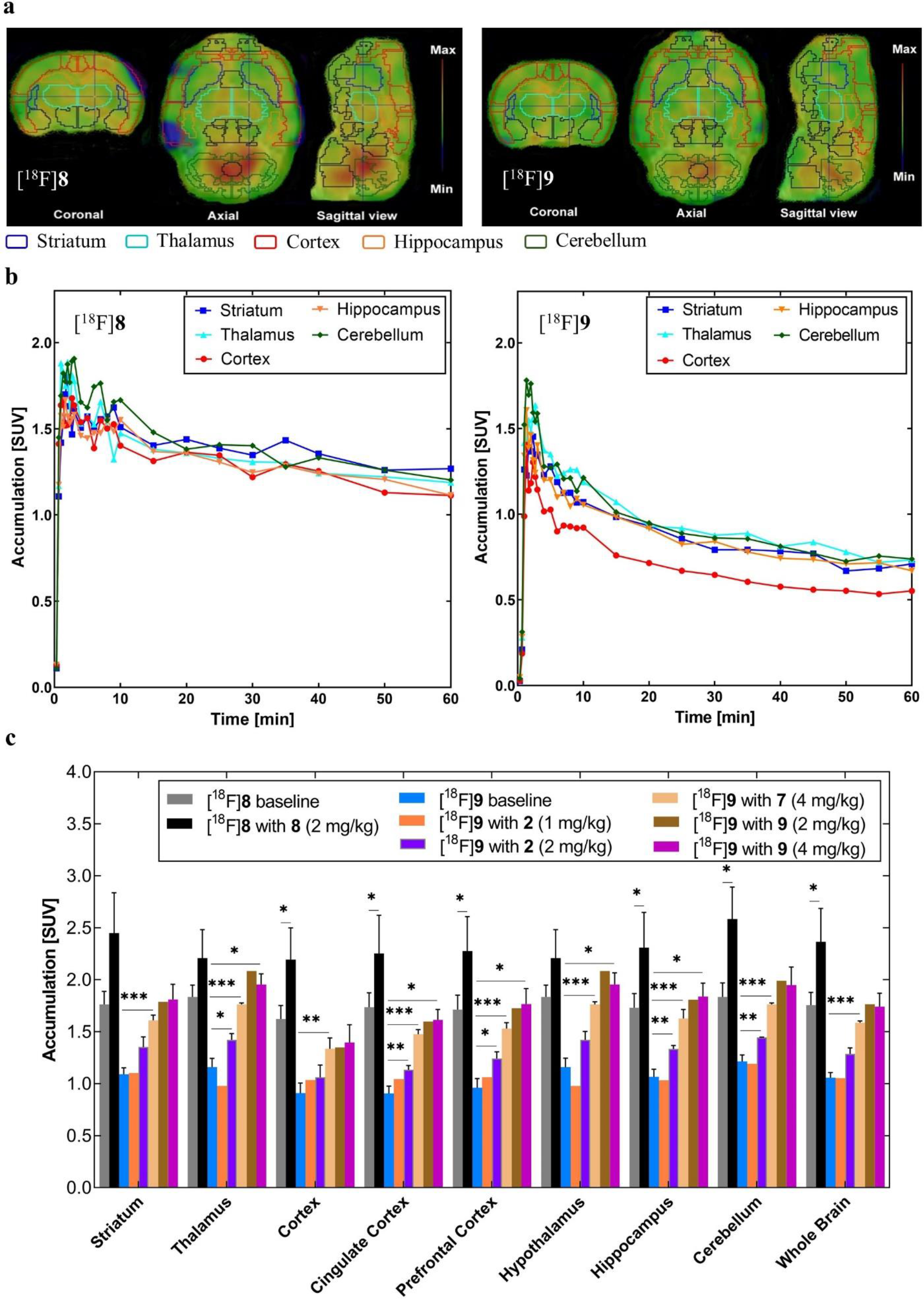
*In vivo* characterization results of [^18^F]**8** and [^18^F]**9** in rat brain. (**a**) Regional distribution of [^18^F]**8** and [^18^F]**9** at time interval of 2-20 min overlaid by color-coded regions of interest; (**b**) Representative TACs from the baseline studies in different brain areas; (**c**) Radioactivity uptake after pretreatment with different classes of mGluR2 PAMs using different doses at the time window of 2-20 min after administration of radioligand. \**p* ≤ 0.05, \*\**p* ≤ 0.01, and \*\*\**p* ≤ 0.001 (vs baseline). Pictures were rendered from Prism 9.0.

### PET imaging in a non-human primate

The feasibility of [^18^F]**9** as an mGluR2 PET imaging ligand was further evaluated in a cynomolgus monkey. In each study, a 120-min dynamic PET acquisition was performed following the injection of [^18^F]9. Compound 2 was used as a blocking agent in the monkey study, and it was injected 10 min before tracer injection. Parallel to this process, arterial blood was sampled to measure the tracer metabolism, plasma free fraction (*f_p_*), and the time courses of [^18^F]**9** concentration in the whole-blood (WB) and plasma (PL). As shown in Figure 5a, the WB/PL ratio of [^18^F]**9** was consistent across the studies and reached a plateau after 1 min post tracer injection with a mean value of 1.12 ± 0.09. RadioHPLC measurements indicated the presence of primarily polar and moderately polar metabolites as shown by the radiochromatograms (Figure 5b). Percent parent in plasma measurements revealed a rather slow rate of metabolism with 49.3 ± 1.4% of plasma activity attributable to unmetabolized [^18^F]**9** at 90 min (Figure 5c). Figure 5d shows the corresponding individual metabolite-corrected [^18^F]**9** SUV time courses in plasma. Plasma free fraction was 0.12 ± 0.06 across studies (range 0.16-0.076).

**Figure 5.**
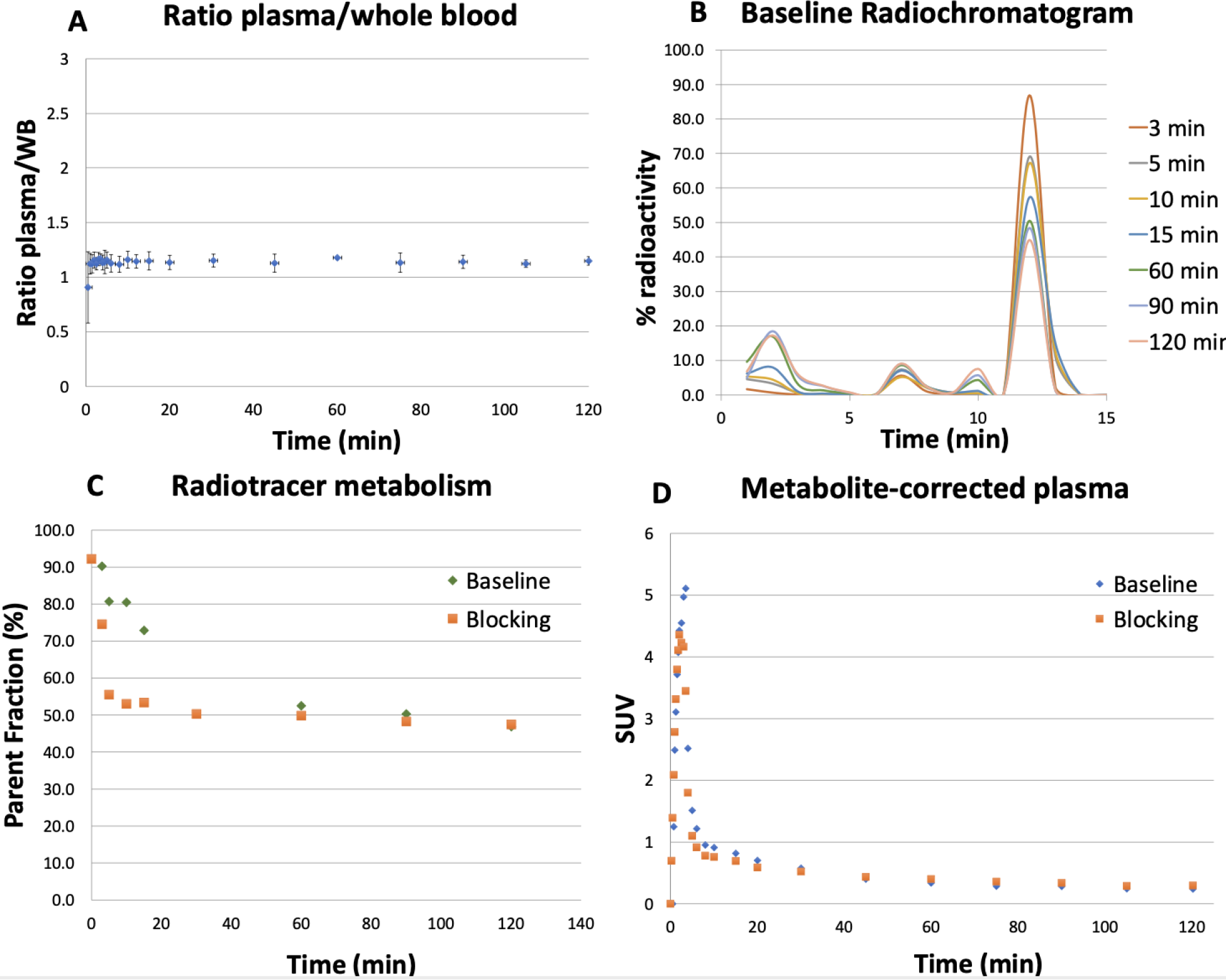
[^18^F]**9** analysis in arterial blood. (**a**) WB/PL ratio; (**b**) Representative radioHPLC chromatogram of plasma samples; (**c**) Individual time course of [^18^F]**9** percent parent in plasma (%PP); (**d**) Individual metabolite-corrected [^18^F]**9** SUV time courses in plasma.

[^18^F]**9** entered the monkey brain readily and peaked at 6 min post tracer injection (SUV > 3.5). Brain uptake and kinetics revealed heterogenous distribution of [^18^F]**9** across brain regions. Figures 6a-b show the selected brain TACs along with compartmental model fits for the cerebellum non vermis, striatum, thalamus, frontal cortex, hippocampus and whole brain from the baseline condition and after pretreatment with BINA (**2**, 0.5 mg/kg, iv.) 10 min prior to PET data acquisition. According to the Akaike information criteria (AIC),^52^ the preferred model was a reversible 2T model with a fixed vascular contribution *v* included as a model parameter (2T4k1v). This model provided stable regional total volume of distribution *V_T_* estimates under both baseline and blocking conditions. The modeling parameters and regional *V_T_* values derived from 2T4k1v and Logan graphical analysis were provided in supporting information for different brain regions in a baseline study (Tables S3 and S4). Logan plots linearized very well by a t* of 30 min (Supporting Information, Figure S6). *K_1_* values, reflecting tracer delivery, was ∼0.25 mL/min/cc in the whole brain based on the 2T model, indicating high brain penetration (Supporting Information, Table S3). The Logan *V_T_* estimates from this representative study demonstrated enhanced [^18^F]**9** uptake in brain areas with high tracer uptake as the thalamus (∼3.3%) after pretreatment with 0.5 mg/kg compound **2** (Figure 6c). Figure 6d shows the corresponding Logan *V_T_* images.

**Figure 6.**
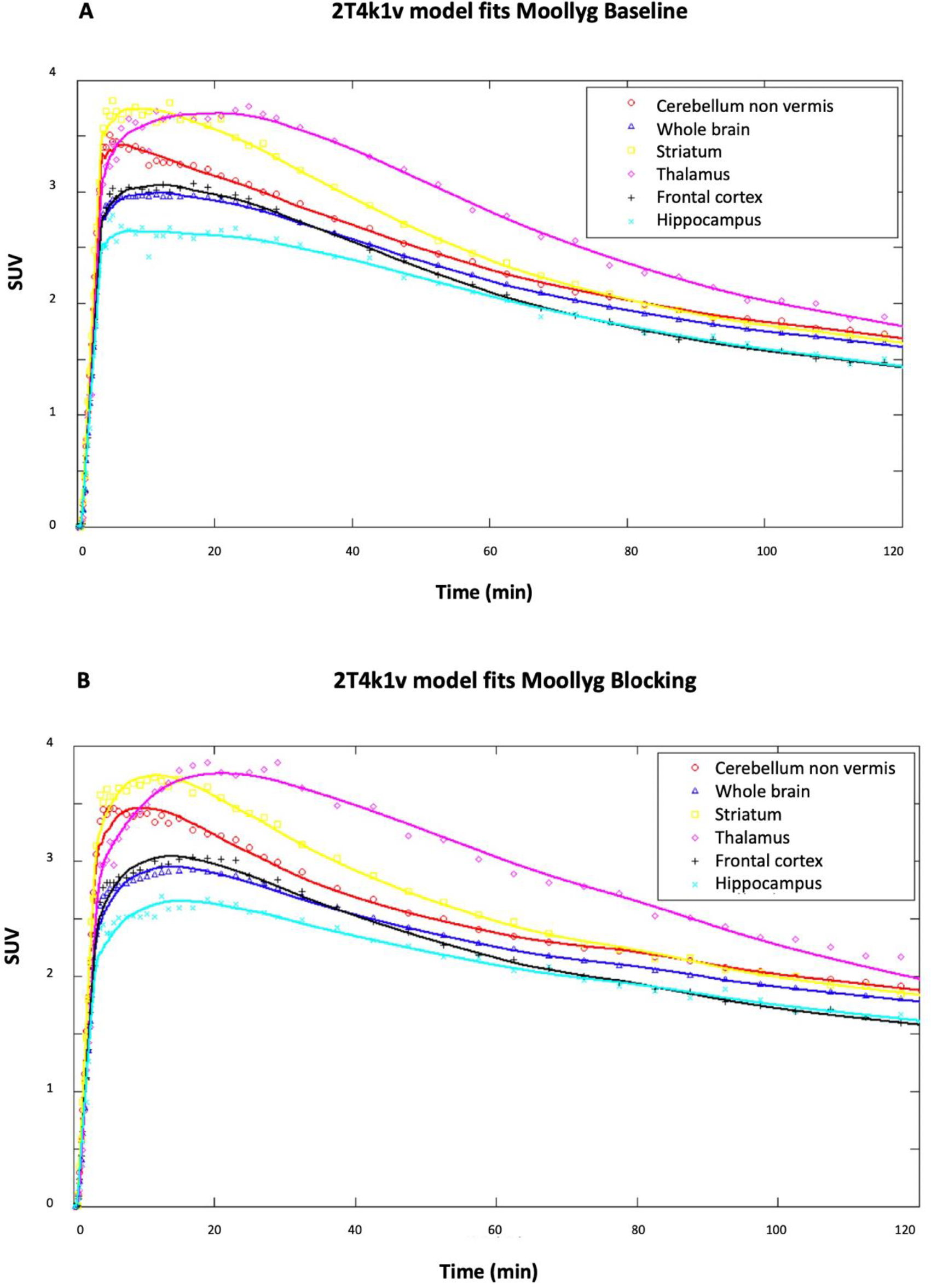
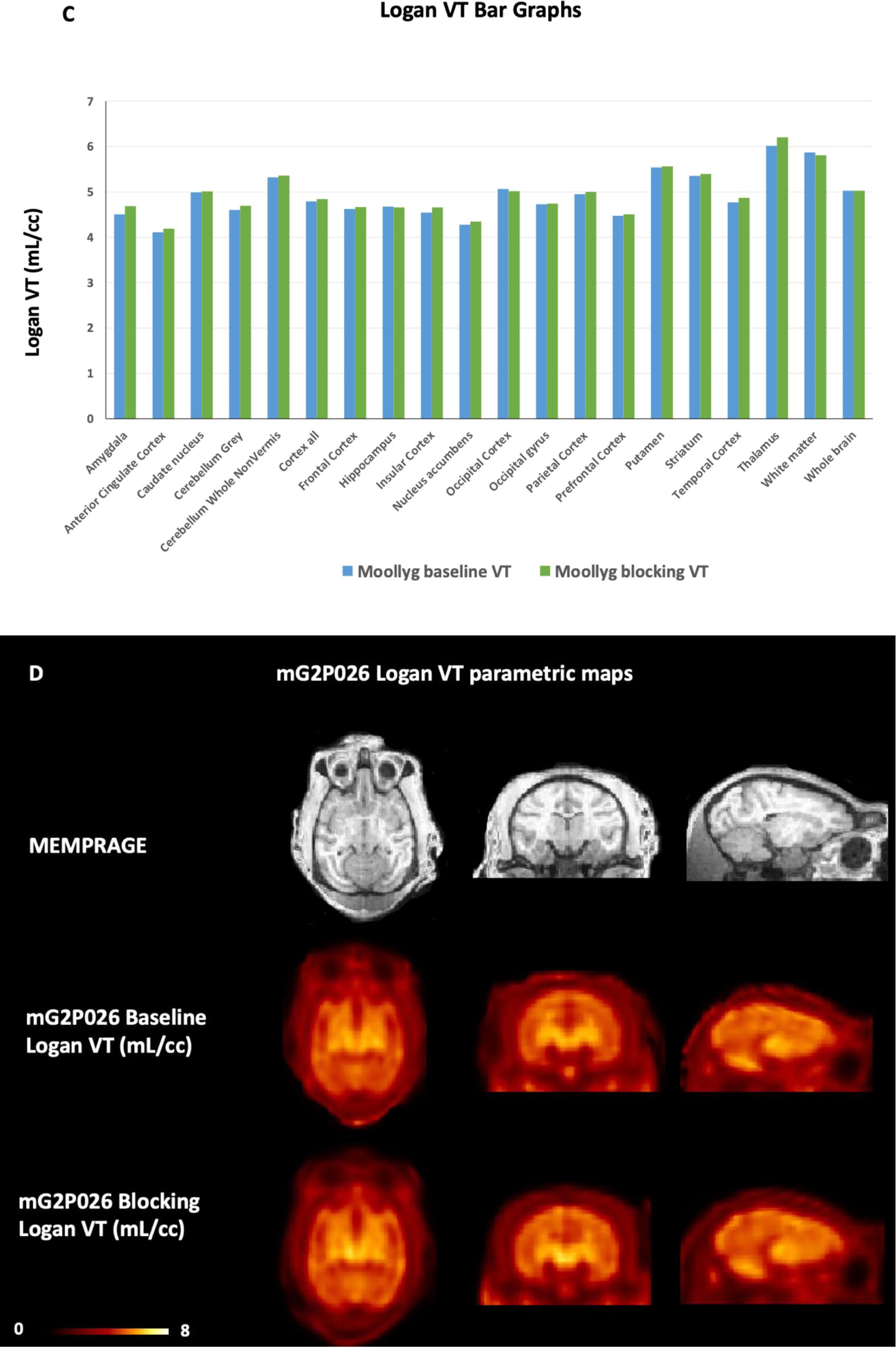
[^18^F]mG2P026 (**9**) in the primate brain. [^18^F]**9** kinetics and 2T4k1v model fits in six brain areas, obtained for a baseline study (**a**) and after pretreatment with 0.5 mg/kg compound **2** (**b**); (**c**) Logan *V_T_* bar graph of [^18^F]**9** at different brain regions; (**d**) Structural MRI (MEMPRAGE) and [^18^F]**9** Logan *V_T_* images at t*30min for the baseline (middle panel) and blocking conditions (bottom panel). Images are presented in the NIMH Macaque Template (NMT)^59^ space.

### Direct comparison between [^18^F]**9** and [^18^F]**6** using a graphical method

[^18^F]**9** displayed an enhanced heterogeneity across brain regions compared to [^18^F]**6** based on its TACs and summed SUV images at 0-30 min (Supporting Information, Figures S7 and S8). We further compared the performance of [^18^F]**9** as an mGluR2 PET imaging ligand with [^18^F]**6** using the graphical method reported by Guo et al.^53^ As Figure 7 shows, a comparison of the regional Logan *V_T_* values of [^18^F]**9** and [^18^F]**6** indicates a strong linear relationship. The negative y-intercept of this linear regression suggests that [^18^F]**9** has a higher specific binding than [^18^F]6.

**Figure 7.**
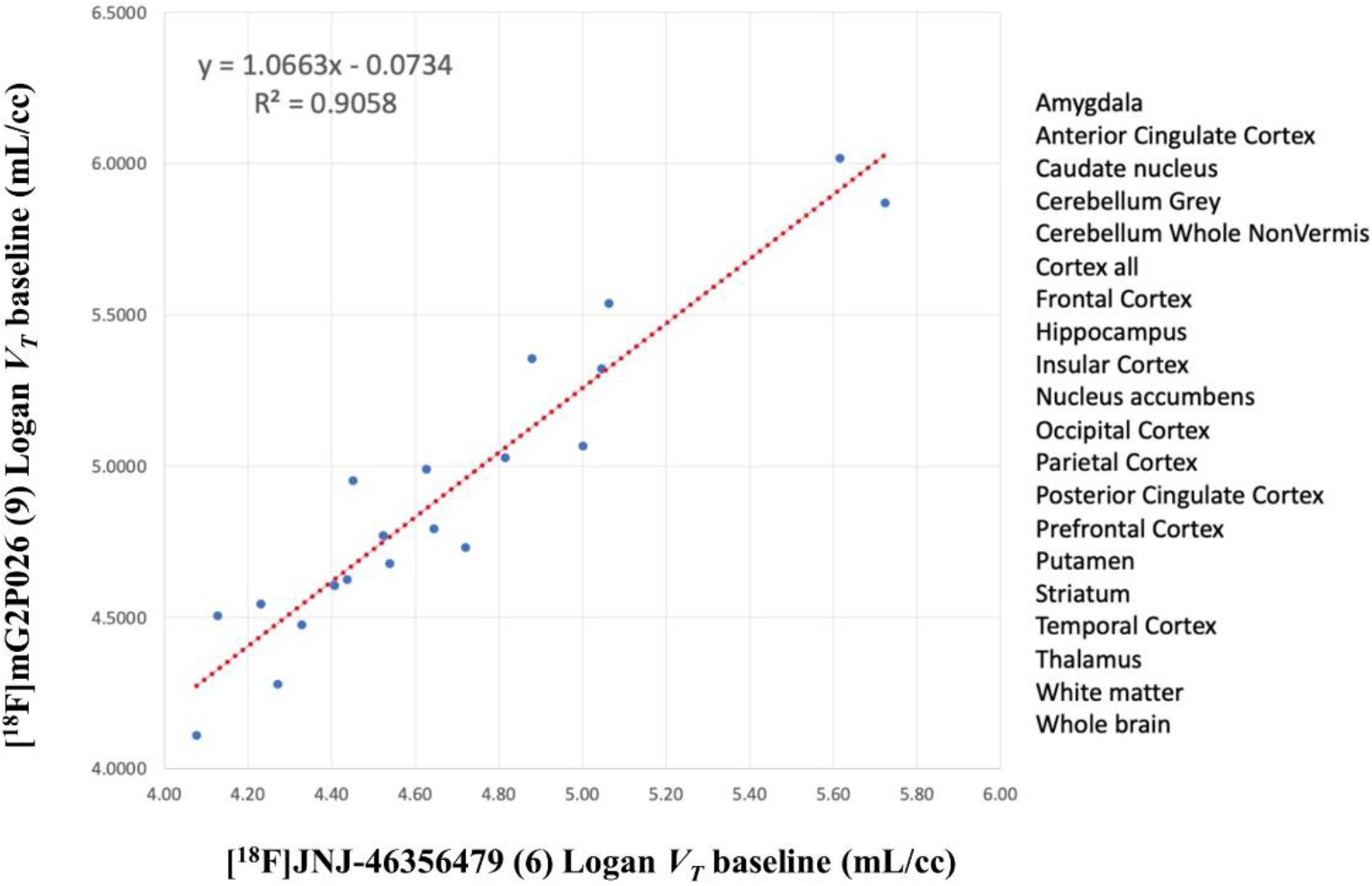
Scatter plots comparing the Logan *V_T_* values obtained for [^18^F]**9** and [^18^F]**6**.

## CONCLUSIONS

We have developed a high contrast PET radioligand [^18^F]mG2P026 (**9**) for imaging mGluR2 in the brain. Compared to compound **6**, the ago-PAM **9** had a 16-fold increase of the PAM activity (EC_50_ = 11 nM) and an apparent mGluR2 agonist activity (IC_50_ = 84 nM). Radiolabeling of compound **9** was achieved under the same conditions as [^18^F]**6**, but with increased radiochemical yield and molar activity. [^18^F]**9** readily entered the rat brain with better imaging characteristics than the piperidine analogue [^18^F]**8**. Distinct from the orthosteric radioligands, pretreatment with mGluR2 PAMs BINA (**2**), compounds **6** and **9** led to an increased [^18^F]**9** uptake in the rat brain, where the self-blocking had the most enhancement. Further characterization of [^18^F]**9** in a non-human primate revealed moderate metabolism stability, fast binding kinetics and feasibility in mapping the mGlu2 receptors. The *V_T_* values derived from the 2T4k1v and Logan models demonstrated a consistent radioactivity enhancement following the pretreatment of compound **2** (0.5 mg/kg, iv.). As a high contrast imaging ligand, [^18^F]**9** was further compared with [^18^F]**6** in the same monkey using a graphical method, where [^18^F]**9** showed an enhanced heterogeneity and a higher specific binding. Therefore, [^18^F]**9** is a promising PET imaging ligand for mGluR2 and can be potentially applied in the disease models and translated to human clinical studies.

## EXPERIMENTAL SECTION

### Animal Procedures

Animal studies were approved and performed following the guidelines of the Subcommittee on Research Animals of the Massachusetts General Hospital and Harvard Medical School in accordance with the Guide of NIH for the Care and Use of Laboratory Animals.

### Chemistry

All chemical reagents and solvents were purchased from the commercial sources and used without further purification. Thin layer chromatography (TLC) was performed on silica gel TLC aluminum foils (Supelco). Flash column chromatography was carried out on silica gel, particle size 60 Å, 230-400 mesh (Supelco). Microwave assisted reactions were done in a CEM Discover microwave synthesizer. Nuclear magnetic resonance (NMR) spectra were collected with a JEOL 500 MHz spectrometer using chloroform-d (CDCl3) and methanol-d_4_ (CD_3_OD) as solvents. Chemical shifts (δ) are assigned as parts in per million (ppm) downfield from tetramethylsiliane. Coupling constants (*J*) are expressed in hertz. Splitting patterns are described as s (singlet), d (doublet), t (triplet), q (quartet), or m (multiplet). Liquid chromatography-mass spectrometry (LCMS) was performed using a 1200 series HPLC system (Agilent Technologies, Canada) comprising a multi-wavelength UV detector, a model 6310 ion trap mass spectrometer (Santa Clara, CA), and an analytical column (Agilent Eclipse C8, 150 mm × 4.6 mm, 5 μm). The samples were subjected to a gradient elution with the mobile phase of 0.1% formic acid solution of water (A) and acetonitrile (B) over 7 min unless otherwise noted at a flow rate of 0.7 mL/min before the MS spectrometer. Purities of all new compounds were determined via the analytical reverse phase HPLC in the LCMS system, using the area percentage method on the UV trace scanning under a wavelength of 254 nm. The compounds were found to have ≥ 95% purity unless otherwise noted. High-Resolution Mass Spectrometry (HRMS) was obtained from Harvard Center for Mass Spectrometry (Cambridge, MA). The electrospray ionization (ESI) technique was used with Thermo_q-Exactive_Plus_I Mass Spectrometer.

### General Procedure for the reductive amination^28, 29^

To a solution of amine (1.1 equiv.) in anhydrous dichloromethane was added triethylamine (4.0 equiv.), magnesium sulfate (10.0 equiv.), and aldehydes (1.0 equiv.). The reaction mixture was stirred at room temperature for 30 min under argon, and then, sodium triacetoxyborohydride (1.5 equiv.) was added. The mixture was stirred at room temperature for another 16 h and was diluted with dichloromethane and washed with water. The aqueous layer was washed two times with dichloromethane and the combined organic layers were dried over magnesium sulfate. The solvent was reduced in vacuo, and the residue was purified by flash column chromatography to give the desired product.

#### 3-(cyclopropylmethyl)-7-((4-(2,4-difluorophenyl)piperidin-1-yl)methyl)-8-(trifluoromethyl)-[1,2,4]triazolo[4,3-a]pyridine (**8**)

Starting from **14** (50 mg, 0.19 mmol) and 4-(2,4-difluorophenyl)piperidine (**15**, 41.2 mg, 0.21 mmol) and following the general procedure described for the reductive amination, compound **8** was obtained as a white solid (40.0 mg, 46.7%). LCMS: m/z 451.2 [M+H]^+^, t_R_ = 3.02 min. ^1^H NMR (500 MHz, CD_3_OD) *δ* ppm 8.52 (d, *J* = 7.2 Hz, 1H), 7.53 (d, *J* = 7.2 Hz, 1H), 7.29-7.30 (m, 1 H), 6.84-6.89 (m, 2H), 3.80 (s, 2H), 3.09 (d, *J* = 6.9 Hz, 2H), 2.96 (d, *J* = 10.9 Hz, 2H), 2.84-2.85 (m, 1H), 2.28 (t, *J* = 11.4Hz, 2H), 1.76-1.84 (m, 4H), 1.24-1.26 (m, 1H), 0.58-0.61 (m, 2H), 0.33-0.35 (m, 2H). ^13^C NMR (125 MHz, CD_3_OD): 166.0 (dd, *J* = 12.0, 99.0 Hz), 164.0 (dd, *J* = 12.0, 100.2 Hz), 152.0, 150.2, 147.2, 132.7 (d, *J* = 3.6 Hz), 132.5 (m), 129.8, 127.3 (q, *J* = 274.7 Hz), 119.4, 118.6 (q, *J* = 33.8 Hz), 114.7 (d, *J* = 18.0 Hz), 107.0 (t, *J* = 26.3 Hz), 61.6, 58.0, 38.9, 35.9, 32.3, 11.9, 8.0. HRMS (ESI^+^) for C_23_H_24_F_5_N_4_^+^ [M+H]^+^ requires m/z = 451.1916, found 451.1920.

#### 7-((4-(2-chloro-4-fluorophenyl)piperazin-1-yl)methyl)-3-(cyclopropylmethyl)-8-(trifluoromethyl)-[1,2,4]triazolo[4,3-a]pyridine (**9**)

Starting from **14** (70 mg, 0.26 mmol) and 1-(2-chloro-4-fluorophenyl)piperazine (**16**, 62.3 mg, 0.29 mmol) and following the general procedure described for the reductive amination, compound **9** was obtained as a white solid (35.0 mg, 28.8%). LCMS: m/z 468.1 [M+H]^+^, t_R_ = 8.49 min (15-min gradient elution). ^1^H NMR (500 MHz, CD_3_OD) *δ* ppm 8.53 (t, *J* = 7.0 Hz, 1H), 7.51 (t, *J* = 7.0 Hz, 1H), 7.12-7.19 (m, 2H), 6.99-7.04 (m, 1H), 3.84 (s, 2H), 3.10 (t, *J* = 7.0 Hz, 2H), 2.98-3.04 (m, 4H), 2.66-2.70 (m, 4H), 1.23-1.27 (m, 1H), 0.58-0.62 (m, 2H), 0.33-0.35 (m, 2H). ^13^C NMR (125 MHz, CD_3_OD): 158.5 (d, *J* = 243.6 Hz), 148.1, 146.2, 145.9, 142.8, 129.5 (d, *J* = 10.6 Hz), 125.9, 123.4 (q, *J* = 274.9 Hz), 121.4 (d, *J* = 9.0 Hz), 117.2 (d, *J* = 25.9 Hz), 115.6, 114.9 (q, *J* = 33.9 Hz), 114.0 (d, *J* = 21.8 Hz), 57.4, 53.2, 51.4, 28.3, 7.9, 4.1. HRMS (ESI^+^) for C_22_H_23_ClF_4_N_5_^+^ [M+H]^+^ requires m/z = 468.1573, found 468.1578.

#### 7-((4-(4-chloro-2-fluorophenyl)piperazin-1-yl)methyl)-3-(cyclopropylmethyl)-8-(trifluoromethyl)-[1,2,4]triazolo[4,3-a]pyridine (**10**)

Starting from **14** (70 mg, 0.26 mmol) and 1-(4-chloro-2-fluorophenyl)piperazine (**17**, 62.3 g, 0.29 mmol) and following the general procedure described for the reductive amination, compound **10** was obtained as a white solid (25.0 g, 20.6%). LCMS: m/z 467.9 [M+H]^+^, t_R_ = 8.63 min (15-min gradient elution). ^1^H NMR (500 MHz, CDCl_3_) *δ* ppm 8.04 (d, *J* = 7.2 Hz, 1H), 7.38 (d, *J* = 7.2 Hz, 1H), 7.03-7.05 (m, 2H), 6.84 (t, *J* = 9.1 Hz, 1H), 3.80 (s, 2H), 3.08-3.10 (m, 6H), 2.67-2.68 (m, 4H), 1.18-1.19 (m, 1H), 0.61-0.65 (m, 2H), 0.33-0.36 (m, 2H). ^13^C NMR (125 MHz, CD_3_OD): 159.3 (d, *J* = 248.8 Hz), 152.1, 150.2, 146.6, 143.0 (d, *J* = 8.7 Hz), 130.6 (d, *J* = 9.8 Hz), 128.3, 126.2, 123.8, 120.2 (d, *J* = 24.8 Hz), 119.5, 118.9 (q, *J* = 33.9 Hz), 61.3, 56.9, 54.3, 30.3, 11.9, 8.0. HRMS (ESI^+^) for C_22_H_23_ClF_4_N_5_^+^ [M+H]^+^ requires m/z = 468.1573, found 468.1578.

#### 3-(cyclopropylmethyl)-7-((4-(2-fluorophenyl)piperazin-1-yl)methyl)-8-(trifluoromethyl)-[1,2,4]triazolo[4,3-a]pyridine (**11**)

Starting from **14** (0.10 g, 0.37 mmol) and 1-(2-fluorophenyl)piperazine (**18**, 64.5 µL, 0.41 mmol) and following the general procedure described for the reductive amination, compound **11** was obtained as a white solid (55.4 mg, 34.6%). LCMS: m/z 434.4 [M+H]^+^, t_R_ = 3.38 min. ^1^H NMR (500 MHz, CD_3_OD) *δ* ppm 8.53 (dd, *J* = 7.5, 9.5 Hz, 1H), 7.51 (dd, *J* = 7.5, 9.5 Hz, 1H), 6.93-7.06 (m, 4H), 3.84 (d, *J* = 8.3 Hz, 2H), 3.08-3.12 (m, 6H), 2.67-2.69 (m, 4H), 1.24-1.26 (m, 1H), 0.59-0.63 (m, 2H), 0.33-0.36 (m, 2H). ^13^C NMR (125 MHz, CD_3_OD): 155.8 (d, *J* = 244.9 Hz), 148.1, 146.2, 142.7, 140.0 (d, *J* = 8.5 Hz), 126.6, 124.4 (d, *J* = 2.9 Hz), 123.4 (q, *J* = 274.9 Hz), 122.6 (d, *J* = 7.9 Hz), 118.9 (d, *J* = 1.9 Hz), 115.6 (d, *J* = 20.6 Hz), 115.5, 114.9 (q, *J* = 33.7 Hz), 57.4, 53.1, 50.5, 28.3, 7.9, 4.1. HRMS (ESI^+^) for C_22_H_24_F_4_N_5_^+^ [M+H]^+^ requires m/z = 434.1962, found 434.1966.

#### 3-chloro-2-hydrazineyl-5-(trifluoromethyl)pyridine (**20**)

To a solution of 2,3-dichloro-5-(trifluoromethyl)pyridine (**19**, 1.0 g, 4.63 mmol) in 1,4-dioxane (7.0 mL) was added hydrazine monohydrate (2.25 mL, 27.8 mmol). The reaction was heated in a sealed reactor at 80 °C for 16 h. The reaction mixture was then cooled to room temperature, quenched with NH_4_OH (32% aqueous solution) and reduced in vacuo. The resulting residue was taken up by ethanol and heated to reflux. After filtering off the solid while hot, the filtrate was concentrated in vacuo to give compound **20** as a white solid (0.3 g, 30.6%). LCMS: m/z 211.9 [M+H]^+^, t_R_ = 2.93 min (8-min gradient elution). ^1^H NMR (500 MHz, CDCl_3_) *δ* ppm 8.33 (s, 1H), 7.65 (d, *J* = 1.8 Hz, 1H), 6.55 (s, 1H), 4.04 (s, 2H). ^13^C NMR (125 MHz, CDCl_3_): 157.1, 143.6 (d, *J* = 4.5 Hz), 133.1 (d, *J* = 2.9 Hz), 123.6 (q, *J* = 271.2 Hz), 117.3 (d, *J* = 33.5 Hz), 114.4. HRMS (ESI^+^) for C_6_H_6_ClF_3_N_3_^+^ [M+H]^+^ requires m/z = 212.0197, found 212.0200.

#### N’-(3-chloro-5-(trifluoromethyl)pyridin-2-yl)-2-cyclopropylacetohydrazide (**21**)

To a solution of **20** (0.3 g, 1.40 mmol) in anhydrous dichloromethane (4.0 mL) was added triethylamine (0.3 mL, 1.28 mmol) and cyclopropylacetyl chloride (0.2 g, 1.70 mmol) at 0 °C. The reaction was stirred at room temperature for 16 h, and then quenched with saturated NaHCO_3_ solution. The resulting solution was extracted three times with dichloromethane and the combined organic layers were dried over magnesium sulfate and reduced in vacuo to give **21** as a white solid (0.17 g, 41.4%). LCMS: m/z 294.0 [M+H]^+^, t_R_ = 4.45 min (8-min gradient elution). ^1^H NMR (500 MHz, CDCl_3_) *δ* ppm 8.32 (s, 1H), 8.29 (d, *J* = 4.5 Hz, 1H), 7.74 (d, *J* = 1.8 Hz, 1H), 7.59 (d, *J* = 4.9 Hz, 1H), 2.31 (d, *J* = 7.2 Hz, 2H), 1.08-1.11 (m, 1H), 0.66-0.69 (m, 2H), 0.28-0.31 (m, 2H). ^13^C NMR (125 MHz, CDCl_3_): 170.7, 154.5, 143.5 (d, *J* = 4.0 Hz), 134.0, 123.3 (q, *J* = 271.4 Hz), 119.3 (q, *J* = 33.8 Hz), 115.2, 39.6, 6.84, 4.84. HRMS (ESI^+^) for C_11_H_11_ClF_3_N_3_O^+^ [M+H]^+^ requires m/z = 294.0616, found 294.0619.

#### 8-chloro-3-(cyclopropylmethyl)-6-(trifluoromethyl)-[1,2,4]triazolo[4,3-a]pyridine (**22**)

To a solution of compound **21** (0.113 g, 0.39 mmol) in anhydrous 1,2-dichloroethane (1.0 mL) was added POCl_3_ (0.073 mL, 0.77 mmol) in a microwave reactor. The reaction mixture was heated at 150 °C for 10 min in the CEM microwave reactor, and then quenched with a solution of saturated NaHCO_3_. The organic layer was isolated, and the aqueous layer was extracted twice with dichloromethane. The combined organic layers were dried with magnesium sulfate, and the volatiles were reduced under vacuo. The residue was purified by flash column chromatography to give **22** as a brown oil (54.0 mg, 50.9%). LCMS: m/z 276.0 [M+H]^+^, t_R_ = 4.39 min (8-min gradient elution). ^1^H NMR (500 MHz, CDCl_3_) *δ* ppm 8.28-8.29 (m, 1H), 7.42 (d, *J* = 1.1 Hz, 1H), 3.12 (d, *J* = 6.8 Hz, 2H), 1.19-1.22 (m, 1H), 0.66-0.70 (m, 2H), 0.35-0.38 (m, 2H). ^13^C NMR (125 MHz, CDCl_3_): 149.8, 147.7, 124.6, 122.4 (dd, *J* = 261.5, 554.0 Hz), 121.3, 120.4 (q, *J* = 5.6 Hz), 118.4 (q, *J* = 35.0 Hz), 29.6, 8.4, 5.4. HRMS (ESI^+^) for C_11_H_10_ClF_3_N_3_^+^ [M+H]^+^ requires m/z = 276.0510, found 276.0513.

#### 3-(cyclopropylmethyl)-6-(trifluoromethyl)-8-vinyl-[1,2,4]triazolo[4,3-a]pyridine (**23**)

In a microwave reactor, **22** (82.0 mg, 0.30 mmol), vinylboronic acid pinacol ester (61.0 μL, 0.36 mmol), Pd(PPh3)4 (17.3 mg, 0.015 mmol), and saturated aqueous NaHCO_3_ solution (0.62 mL) were mixed in 1,4-dioxane (3.2 mL). The reaction mixture was heated at 150 °C for 15 min in the CEM microwave reactor. After cooling to room temperature, the reaction mixture was diluted with ethyl acetate/water. The aqueous layer was extracted three times with ethyl acetate. The combined organic layers were washed with brine and dried over magnesium sulfate. The solvent was reduced in vacuo and the residue was purified by flash column chromatography to give **23** as a yellow solid (47.0 mg, 58.6%). LCMS: m/z 268.0 [M+H]^+^, t_R_ = 4.73 min (8-min gradient elution). ^1^H NMR (500 MHz, CDCl_3_) *δ* ppm 8.22 (s, 1H), 7.25 (s, 1H), 6.99 (d, *J* = 6.0 Hz, 2H), 5.84 (t, *J* = 6.1 Hz, 1H), 3.11 (d, *J* = 6.8 Hz, 2H), 1.21-1.25 (m, 1H), 0.65-0.68 (m, 2H), 0.34-0.37 (m, 2H). ^13^C NMR (125 MHz, CDCl_3_): 148.2, 130.3, 128.0, 124.5, 123.1 (q, *J* = 271.6 Hz), 119.9, 119.8, 119.7, 118.5 (q, *J* = 34.1 Hz), 29.4, 8.5, 5.3. HRMS (ESI^+^) for C_13_H_13_F_3_N_3_^+^ [M+H]^+^ requires m/z = 268.1056, found 268.1061.

#### 3-(cyclopropylmethyl)-6-(trifluoromethyl)-[1,2,4]triazolo[4,3-a]pyridine-8-carbaldehyde (**24**)

To a solution of **23** (0.54 g, 2.02 mmol) in water (5.4 mL) and 1,4-dioxane (21.6 mL) was added sodium periodate (1.30 g, 6.08 mmol) and osmium tetroxide (2.5% in tert-butanol, 1.03 mL, 0.08 mmol). The reaction mixture was stirred at room temperature for 2 h. After the reaction was completed, the mixture was diluted with ethyl acetate/water. The aqueous layer was extracted three times with ethyl acetate. The combined organic layers were washed with brine and dried over magnesium sulfate. The solvent was reduced in vacuo and the residue was purified by flash column chromatography to give **24** as a brown oil (0.224 g, 41.2%). LCMS: m/z 270.0 [M+H]^+^, t_R_ = 3.94 min (8-min gradient elution). ^1^H NMR (500 MHz, CDCl_3_) *δ* ppm 10.78 (s, 1H), 8.55 (d, *J* = 1.1 Hz, 1H), 7.99 (d, *J* = 1.3 Hz, 1H), 3.17 (d, *J* = 6.8 Hz, 2H), 1.23-1.24 (m, 1H), 0.69-0.72 (m, 2H), 0.37-0.40 (m, 2H). ^13^C NMR (125 MHz, CDCl_3_): 186.3 (d, *J* = 4.2 Hz), 148.7, 147.7, 125.7, 124.1, 123.9, 122.6 (q, *J* = 271.7 Hz), 118.3 (q, *J* = 35.2 Hz), 29.5, 8.4, 5.5. HRMS (ESI^+^) for C_12_H_11_F_3_N_3_O^++^ [M+H]^+^ requires m/z = 270.0849, found 270.0853.

#### 3-(cyclopropylmethyl)-8-((4-(2,4-difluorophenyl)piperazin-1-yl)methyl)-6-(trifluoromethyl)-[1,2,4]triazolo[4,3-a]pyridine (**12**)

Starting from **24** (50 mg, 0.19 mmol) and 4-(2,4-difluorophenyl)piperidine (41.2 mg, 0.21 mmol) and following the procedure described for the reductive amination, compound **12** was obtained as a white solid (22.0 mg, 25.6%). LCMS: m/z 452.1 [M+H]^+^, t_R_ = 4.28 min (8-min gradient elution). ^1^H NMR (300 MHz, CD_3_OD) *δ* ppm 9.16 (s, 1H), 7.90 (s, 1H), 7.02-7.10 (m, 1H), 6.82-6.93 (m, 2H), 4.02 (s, 2H), 3.07-3.10 (m, 4H), 2.84 (d, *J* = 7.0 Hz, 2H), 2.75-2.79 (m, 4H), 1.21-1.31 (m, 1H), 0.53-0.59 (m, 2H), 0.28-0.33 (m, 2H). ^13^C NMR (75 MHz, CD_3_OD): 168.2, 151.5, 136.6 (d, *J* = 9.0 Hz), 127.3, 126.3 (d, *J* = 5.4 Hz), 124.6 (d, *J* = 2.7 Hz), 123.4 (d, *J* = 270.5 Hz), 119.8 (dd, *J* = 4.1, 9.3 Hz), 117.5 (d, *J* = 34.7 Hz), 110.5 (d, *J* = 3.8 Hz), 110.2 (d, *J* = 3.8 Hz), 103.8 (d, *J* = 51.6 Hz), 103.9, 55.9, 53.0, 50.6, 32.8, 9.2, 3.8. HRMS (ESI^+^) for C_22_H_23_F_5_N_5_^+^ [M+H]^+^ requires m/z = 452.1868, found 452.1873.

#### Diethyl 2-(((1-(cyclopropylmethyl)-1H-pyrazol-5-yl)amino)methylene)malonate (**27**)

A solution of diethyl 2-(ethoxymethylene)malonate (**25**, 1.62 mL, 8.02 mmol) and 1-(cyclopropylmethyl)-1H-pyrazol-5-amine (**26**, 1.0 g, 7.29 mmol) in ethanol was heated to reflux overnight. The solvent was removed in vacuo. The residue was purified via flash column chromatography to give **27** as a pale yellow solid (3.94 g, 88.0%). LCMS: m/z 308.0 [M+H]^+^, t_R_ = 3.58 min. ^1^H NMR (500 MHz, CDCl_3_) *δ* ppm 11.02 (d, *J* = 12.8 Hz, 1H), 8.16 (d, *J* = 12.8 Hz, 1H), 7.41 (d, *J* = 1.9 Hz, 1H), 6.06 (d, *J* = 1.9 Hz, 1H), 4.31 (q, *J* = 7.1 Hz, 2H), 4.22 (q, *J* = 7.1 Hz, 2H), 3.96 (d, *J* = 6.9 Hz, 2H), 1.37 (t, *J* = 7.1 Hz, 3H), 1.30 (t, *J* = 7.1 Hz, 3H), 1.22-1.26 (m, 1H), 0.59-0.63 (m, 2H), 0.39-0.43 (m, 2H). ^13^C NMR (125 MHz, CD3Cl_3_): 169.1, 165.1, 154.1, 139.2, 138.9, 95.46, 94.31, 60.9, 60.4, 53.2, 14.4, 14.3, 11.2, 4.0. HRMS (ESI^+^) for C_15_H_22_N_3_O_4_^+^ [M+H]^+^ requires m/z = 308.1605, found 308.1608.

#### Ethyl 4-chloro-1-(cyclopropylmethyl)-1H-pyrazolo[3,4-b]pyridine-5-carboxylate (**28**)

A solution of compound **27** (1.57 g, 5.10 mmol) in POCl_3_ (30.0 mL) was refluxed at 120 °C overnight. The reaction mixture was quenched with water at 0 °C and extracted with ethyl acetate. The aqueous layer was further extracted with two times extraction with ethyl acetate. The combined organic layers were washed with brine and dried over magnesium sulfate. The solvent was reduced in vacuo and the residue was purified by flash column chromatography to give **28** as a colorless waxy solid (1.35 g, 67.5%). LCMS: m/z 280.0 [M+H]^+^, t_R_ = 5.54 min (8-min gradient elution). ^1^H NMR (500 MHz, CDCl_3_) *δ* ppm 9.01 (s, 1H), 8.19 (s, 1H), 4.45 (q, *J* = 7.1 Hz, 2H), 4.38 (d, *J* = 7.2 Hz, 2H), 1.43 (t, *J* = 7.1 Hz, 3H), 1.36-1.40 (m, 1H), 0.53-0.57 (m, 2H), 0.44-0.47 (m, 2H). ^13^C NMR (125 MHz, CD3Cl_3_): 164.5, 151.7, 151.1, 139.7, 132.6, 118.4, 116.2, 61.8, 52.4, 14.4, 11.3, 4.0. HRMS (ESI^+^) for C_13_H_15_ClN_3_O_2_^+^ [M+H]^+^ requires m/z = 280.0847, found 280.0852.

#### Ethyl 1-(cyclopropylmethyl)-4-(trifluoromethyl)-1H-pyrazolo[3,4-b]pyridine-5-carboxylate (**29**)

To a solution of **28** (0.10 g, 0.36 mmol) in acetonitrile (5.7 mL) was added sodium iodide (0.108 g, 0.72 mmol) and acetylchloride (0.57 mL). The reaction mixture was stirred at 50 °C overnight. The reaction mixture was diluted with dichloromethane, washed with saturated aqueous NaHCO_3_ solution, and brine. The organic layer was dried over magnesium sulfate, and the solvent was reduced under vacuo. The resulting residue was dissolved in DMF (1.4 mL) and mixed with fluorosulfonyl-difluoro-acetic acid methyl ester (51.0 μL, 0.40 mmol) and copper(I) iodide (76.0 mg, 0.40 mmol) in a sealed tube. The mixture was stirred at 100 °C for 6 h. After cooling, the mixture was diluted with water and extracted with ethyl acetate. The organic layer was washed with brine and reduced under vacuo. The resulting residue was purified by flash column chromatography to give **29** as a white solid (65.0 mg, 57.6% over two steps). LCMS: m/z 313.9 [M+H]^+^, t_R_ = 5.85 min (8-min gradient elution). ^1^H NMR (300 MHz, CDCl_3_) *δ* ppm 8.96 (s, 1H), 8.26 (q, *J* = 2.3 Hz, 1H), 4.42-4.49 (m, 4H), 1.42 (t, *J* = 7.1 Hz, 3H), 1.36-1.41 (m, 1H), 0.54-0.58 (m, 2H), 0.47-0.50 (m, 2H). ^13^C NMR (75 MHz, CDCl_3_): 165.4, 151.2, 150.2, 132.6 (q, *J* = 4.0 Hz), 131.0 (q, *J* = 35.7 Hz), 122.7 (d, *J* = 275.1 Hz), 119.6, 110.9 (d, *J* = 2.4 Hz), 62.4, 52.2, 13.9, 11.2, 3.9. HRMS (ESI^+^) for C_14_H_15_F_3_N_3_O_2_^+^ [M+H]^+^ requires m/z = 314.1111, found 314.1116.

#### 1-(cyclopropylmethyl)-4-(trifluoromethyl)-1H-pyrazolo[3,4-b]pyridine-5-carboxylic acid (**30**)

To a solution of **29** (50 mg, 0.16 mmol) in tetrahydrofuran/water (2.0 mL/0.15 mL, v/v) was added lithium hydroxide monohydrate (20.2 mg, 0.48 mmol). The reaction mixture was stirred at room temperature for 12 h. After the reaction was completed, 1N HCl was added to quench the reaction and the mixture was extracted with ethyl acetate. The organic layer was washed with brine and dried over magnesium sulfate. The solvent was removed under vacuo and the residue was purified by flash column chromatography to give **30** as a white solid (40.0 mg, 87.7%). LCMS: m/z 286.0 [M+H]^+^, t_R_ = 4.70 min (8-min gradient elution). ^1^H NMR (500 MHz, CDCl_3_) *δ* ppm 9.16 (s, 1H), 8.33 (q, *J* = 2.4 Hz, 1H), 4.47 (d, *J* = 7.2 Hz, 2H), 1.40-1.45 (m, 1H), 0.56-0.60 (m, 2H), 0.48-0.51 (m, 2H). ^13^C NMR (125 MHz, CD_3_OD): 166.6, 151.0, 150.4, 132.0 (d, *J* = 3.8 Hz), 130.4 (q, *J* = 35.8 Hz), 122.9 (q, *J* = 274.3 Hz), 120.4, 110.6, 51.8, 10.7, 2.9. HRMS (ESI^+^) for C_12_H_11_F_3_N_3_O_2_^+^ [M+H]^+^ requires m/z = 286.0798, found 286.0802.

#### (1-(cyclopropylmethyl)-4-(trifluoromethyl)-1H-pyrazolo[3,4-b]pyridin-5-yl)(4-(2,4-difluorophenyl)piperazin-1-yl)methanone (**13**)

A solution of **30** (0.12 g, 0.42 mmol) in DMF (4.5 mL) was added HATU (0.16 g, 0.42 mmol) and DIPEA (0.15 mL, 0.84 mmol). After stirring 10 min at room temperature, 4-(2,4-difluorophenyl)piperidine (93.0 mg, 0.465 mmol) was added and stirred for 16 h. The reaction was then quenched with water and extracted with ethyl acetate. The organic layer was dried over magnesium sulfate and the solvent was removed under vacuo. The residue was purified by flash column chromatography to give **13** as a yellow solid (0.15 mg, 76.9%). LCMS: m/z 466.2 [M+H]^+^, t_R_ = 4.38 min. ^1^H NMR (500 MHz, CDCl_3_) *δ* ppm 8.52 (s, 1H), 8.20 (d, *J* = 1.6 Hz, 1H), 6.87-6.92 (m, 1H), 6.79-6.84 (m, 2H), 4.44-4.45 (m, 2H), 4.05-4.09 (m, 1H), 3.96-4.01 (m, 1H), 3.39-3.41 (m, 2H), 3.11-3.14 (m, 2H), 2.92-2.94 (m, 2H), 1.40-1.43 (m, 1H), 0.56-0.58 (m, 2H), 0.47-0.49 (m, 2H). ^13^C NMR (125 MHz, CD3Cl_3_): 165.6, 158.5 (d, *J* = 233.0 Hz), 155.9 (d, *J* = 249.7 Hz), 150.4, 147.1 (d, *J* = 30.4 Hz), 136.0, 131.4, 128.0 (d, *J* = 35.3 Hz), 122.9 (t, *J* = 137.5 Hz), 120.2 (d, *J* = 9.4 Hz), 110.9, 110.8 (d, *J* = 58.4 Hz), 105.1 (d, *J* = 52.3 Hz), 105.0, 52.4, 51.0, 50.7, 47.6, 42.3, 11.2, 4.1. HRMS (ESI^+^) for C_22_H_21_F_5_N_5_O^+^ [M+H]^+^ requires m/z = 466.1661, found 466.1665.

#### 4-(2-fluoro-4-(4,4,5,5-tetramethyl-1,3,2-dioxaborolan-2-yl)phenyl)piperidine (**35**)

To a solution of **34** (0.30 g, 0.74 mmol) in ethanol (10.0 mL) was added Pd/C (0.05 g, 10 wt. %). The resulting solution was subjected to a hydrogen environment at 43 psi at room temperature overnight. The Pd/C was filtered out and the solvent was reduced under vacuo to give a yellow oil LCMS: m/z 350.1 [M+H]^+^ (*t*-butyl cleaved mass was observed), t_R_ = 5.17 min. The yellow oil was then taken up by a solution of 4N HCl in dioxane (2.67 mL) and stirred at room temperature for 1 h. The solvent was removed under vacuo and the residue was used for the next step without further purification. LCMS: m/z 306.1 [M+H]^+^, t_R_ = 2.78 min.

#### 7-((4-(2-chloro-4-(4,4,5,5-tetramethyl-1,3,2-dioxaborolan-2-yl)phenyl)piperazin-1-yl)methyl)-3-(cyclopropylmethyl)-8-(trifluoromethyl)-[1,2,4]triazolo[4,3-a]pyridine (**37**)

Starting from **14** (78.0 mg, 0.29 mmol) and 1-(2-chloro-4-(4,4,5,5-tetramethyl-1,3,2-dioxaborolan-2-yl)phenyl)piperazine·HCl salt (**35**, 99.1 mg, 0.29 mmol) and following the general procedure described for the reductive amination, compound **37** was obtained as a waxy white solid (54.0 mg, 33.3%). LCMS: m/z 559.2 [M+H]^+^, t_R_ = 3.32 min. ^1^H NMR (500 MHz, CDCl_3_) *δ* ppm 8.04 (d, J = 7.2 Hz, 1H), 7.52 (d, J = 7.5 Hz, 1H), 7.42-7.47 (m, 2H), 7.23-7.26 (m, 1 H), 3.75 (d, J = 1.4 Hz, 2H), 3.09 (d, J = 6.7 Hz, 2H), 2.90-2.92 (m, 3H), 2.27-2.32 (m, 2H), 1.79-1.83 (m, 4H), 1.33 (s, 12H), 1.15-1.19 (m, 1H), 0.62-0.64 (m, 2H), 0.34-0.36 (m, 2H). ^13^C NMR (125 MHz, CDCl_3_): 160.6 (d, *J* = 246.2 Hz), 147.7, 147.1, 146.7, 141.5, 135.9, 135.8, 130.6, 127.2, 124.3, 124.1, 116.8, 115.1 (m), 84.1, 57.9, 54.5, 35.5, 32.0, 29.3, 24.8, 8.5, 5.3. HRMS (ESI^+^) for C_29_H_36_BF_4_N_4_O_2_^+^ [M+H]^+^ requires m/z = 559.2862, found 559.2871.

#### 3-(cyclopropylmethyl)-7-((4-(2-fluoro-4-(4,4,5,5-tetramethyl-1,3,2-dioxaborolan-2-yl)phenyl)piperidin-1-yl)methyl)-8-(trifluoromethyl)-[1,2,4]triazolo[4,3-a]pyridine (**38**)

Starting from **14** (100.0 mg, 0.37 mmol) and 4-(2-fluoro-4-(4,4,5,5-tetramethyl-1,3,2-dioxaborolan-2-yl)phenyl)piperidine (**36**, 150.8 mg, 0.42 mmol) and following the general procedure described for the reductive amination, compound **38** was obtained as a waxy brown solid (20.0 mg, 9.4%). LCMS: m/z 576.2 [M+H]^+^, t_R_ = 4.32 min. ^1^H NMR (500 MHz, CDCl_3_) *δ* ppm 8.04 (d, J = 7.3 Hz, 1H), 7.78 (d, J = 1.4 Hz, 1H), 7.63 (dd, J = 1.4, 8.0 Hz, 1H), 7.40-7.45 (m, 1H), 6.97-7.04 (m, 1 H), 3.81 (d, J = 1.5 Hz, 2H), 3.10-3.13 (m, 4H), 3.09 (d, J = 6.7 Hz, 2H), 2.70 (t, J = 4.5 Hz, 4H), 1.32 (s, 12H), 1.17-1.20 (m, 1H), 0.61-0.64 (m, 2H), 0.34-0.36 (m, 2H). ^13^C NMR (125 MHz, CD_3_OD): 151.6, 148.2, 146.1, 136.4, 135.8, 134.1, 133.2, 127.7, 126.1, 123.3 (q, *J* = 274.7 Hz), 119.6, 115.7, 115.2 (d, *J* = 34.5 Hz), 83.9, 57.3, 53.1, 50.5, 28.3, 23.8, 7.9, 4.1. HRMS (ESI^+^) for C_28_H_35_BClF_3_N_5_O_2_^+^ [M+H]^+^ requires m/z = 576.2519, found 576.2530.

### Radiochemistry

The fully automated radiosyntheses of [^18^F]**8** and [^18^F]**9** followed the same procedures described for [^18^F]**6** in GE TRACERLab™ FX_FN_ platform.^32, 49^ Briefly, no carrier added [^18^F]fluoride ion was produced via the ^18^O(p, n)^18^F reaction by irradiating ^18^O-enriched water (Isoflex Isotope, San Francisco, CA) in a GE PETtrace 16.5 MeV cyclotron (GE Healthcare, Waukesha, WI, USA). The [^18^F]fluoride solution was enriched with a QMA Sep-Pak Cartridge (Sep-Pak plus light, Waters, Milford, MA) before released into the reactor by a solution of tetraethylammonium bicarbonate (TEAB, 2.7 mg, 14.1 μmol) in acetonitrile/water (0.7 mL/0.3 mL). Azeotropic drying of the [^18^F]fluoride ion was performed at 80 °C for 10 min and then 100 °C for 3 min with addition of another 1.0 mL anhydrous acetonitrile (1 mL). Then, 0.4 mL *n*-BuOH (0.4 mL) was added to the reactor followed by a solution of **37** or **38** (6.2 μmol) and [Cu(OTf)_2_Py_4_] (9.0 mg, 13.3 μmol) in dimethylacetamide (DMA, 0.8 mL). The reaction mixture was heated at 130 °C for 10 min and then quenched by adding 4.0 mL water at 50 °C. [^18^F]**8** or [^18^F]**9** was then separated with semi-preparative HPLC equipped with an Xbridge BEH C18 OBD column (130 Å, 5 µm, 10×250 mm) eluted with acetonitrile: 0.1M ammonium formate solution [55:45 (v/v)] at 6 mL/min. The fraction containing [^18^F]**8** or [^18^F]**9** was collected, diluted with 25 mL high purity water, and passed through a C18 cartridge (light Sep-Pak, Waters, Milford, MA). The cartridge was then washed with 10 mL water and eluted with 0.6 mL ethanol into a product collection vial. Then, 5.4 mL saline was passed the cartridge into the collection vial to afford the formulated solution. The radiochemical identity, molar activity (A_m_) and purity of the injected radioligands were determined by the radio-HPLC system (Waters 4000) using an XBridge analytical column (C18, 3.5 μm, 4.6 × 150 mm) eluted with acetonitrile: 0.1M ammonium formate solution [60:40 (v/v)] at 1 mL/min and a UV wavelength of 254 nm.

### Molecular modeling

The molecular docking experiments of compounds **6** and **8**-**11** in the allosteric site were done as previously described for compound **7** using the in-house made mGluR2 homology model.^34^ The ligands were prepared and optimized in Avogadro 1.2^54^ before docking. Molecular docking was performed was performed into the homology model using Extra precision Induced Fit Docking in Glide.^55–57^

### In vitro binding assays

The PAM activity of compounds **8**-**11** toward mGluR2 and their selectivity against other mGlu subtypes were determined as previously described for compounds **6** and **7**.^32, 34^ Briefly, the HEK-293 stable cell lines that express mGluR1, mGluR2, mGluR4, mGluR6 or mGluR8 were maintained in complete Dulbecco’s modified Eagle’s medium (DMEM) containing Hygromycin B (100 μg/mL) and Blasticidin (15 μg/mL). The temperature was kept at 37°C with the presence of 5% CO_2_. The ligand binding toward mGluR1 and mGluR5 were measured by using Ca^2+^ mobilization assays. In these assays, the mGluR1 stable cell lines were plated into poly-L-lysine (PLL) coated 384-well black clear bottom cell culture plates with complete Basal Medium Eagle (BME) buffer. The mGluR5 expressing HEK-293 cells were plated into the plate with complete BME buffer. The density was 20,000 cells in 40 µL per well. The ligand binding toward G_i/o_ coupled receptors of mGluR2, mGluR3, mGluR4, mGluR6 and mGluR8 were determined using cAMP assay. The ligand PAM functional activity was tested with Promega’s split luciferase based GloSensor cAMP biosensor assay. In this assay, the changes of the intracellular cAMP concentration were determined in the presence of EC_20_ amount of L-glutamate and compound **2** was used as a reference compound. The glutamate binding co-operativity with compound **11** was also determined using this assay. Each assay was repeated at least three times.

## Data Analysis

The concentration-response curves for each compound were generated using the Prism GraphPad software 9.0 (Graph Pad Inc., San Diego, CA, U.S.). The curves were fitted to a four-parameter logistic equation to determine the EC_50_/IC_50_ values. The EC_50_/IC_50_ are the concentration of a compound that causes a half-maximal potentiation or suppression of the corresponding response in each assay. The glutamate-response curves in the absence or presence of compound **11** were fitted to Black-Leff Ehlert equation using general least squares fit in Prism to determine the glutamate binding co-operativity parameters.

### *In vivo* characterization in rats

The PET imaging studies in rats were performed in the Triumph II Preclinical Imaging System (Trifoil Imaging, LLC, Northridge, CA). During the scan, rats were anesthetized with a flow of 1.0-1.5% isoflurane in oxygen (1-1.5 L/min), and the tail vein was catheterized to inject the radioligands. A dynamic 60-min dynamic acquisition was started from the injection of a radioligand (20-41 MBq iv). The vital signs were monitored throughout the imaging. A CT scan was performed after each PET acquisition to obtain anatomical information and correction for attenuation.

Altogether, five normal Sprague Dawley rats (male, 275-500 g) were used in eight studies to investigate *in vivo* imaging characteristics of [^18^F]**8**. Three rats had control studies followed by the “blocking” studies and one rat had only control or blocking study. The unlabeled compound **8** was used to investigate selectivity and sensitivity of [^18^F]**8** for imaging mGluR2. Compound **8** was formulated in a solution of 10% DMSO, 5% Tween-20 and 85% PBS with a pH under 5.5 and was administered (4 mg/kg, iv) 10 min before the radioactivity. Similarly, six normal Sprague Dawley rats (male, 275-500 g) were used in fourteen studies to investigate *in vivo* imaging characteristics of [^18^F]**9**. Four rats had control studies followed by ten “blocking” studies with additional two rats in the blocking studies. Pretreatments with compounds **2** (four studies), **7** (three studies) and **9** (three studies) were used to investigate the tracer selectivity and sensitivity. All of these compounds were formulated into the 10% DMSO, 5% Tween-20 and 85% PBS with a pH under 5.5 and injected 10 min before the tracer injection. Compound **2** was injected at 1.0 mg/kg and 2.0 mg/kg doses, compound **6** was administered at 4 mg/kg dose, and compound **9** was studied at 2 mg/kg and 4 mg/kg doses.

The PET imaging data were corrected for uniformity, scatter, and attenuation and processed by using maximum-likelihood expectation-maximization (MLEM) algorithm with 30 iterations to dynamic volumetric images (9×20”, 7×60”, 6×300”, 2×600”). The CT data were reconstructed via the modified Feldkamp algorithm using matrix volumes of 512×512×512 and a pixel size of 170 μm. The ROIs, i.e., striatum, thalamus, cortex, hypothalamus, hippocampus, cerebellum, and whole brain, were drawn onto coronal PET slices based on the rat brain atlas. PMOD 3.2 (PMOD Technologies Ltd., Zurich, Switzerland) was used to create the corresponding TACs (time-activity curves) for each tracer. The percent changes in selected brain regions between the baseline and blocking studies were calculated at the time window of 2-20 min after radiotracer injection. The statistical significance was calculated by the two-sample t-test.

### *In vivo* characterization of [^18^F]**9** in primate

#### PET monkey scans

[^18^F]**9** was characterized in a non-human primate (Cynomolgus fascicularis) via the same protocol as described for [^18^F]**6** and [^11^C]**7**.^32, 36^ Briefly, the PET/CT imaging studies were conducted in the same monkey at a Discovery MI (GE Healthcare) PET/CT scanner. Before the scan, the animal was anesthetized with ketamine/xylazine (10/0.5 mg/kg IM) and then maintained in anesthesia using isoflurane (1-2% in 100% O_2_). At baseline, PET acquisition started immediately prior to a 3-min intravenous tracer infusion (138.898 MBq, 375.4 GBq/μmol) and were acquired in 3D list mode for 120 min. Under pretreatment conditions, compound **2** (0.5 mg/kg) was administered intravenously 10 min before [^18^F]**9** (187.368 MBq, 477.6 GBq/μmol) in the same monkey two weeks later.

In each study, a CT scan was carried out to center the head in the imaging field and acquire data for attenuation correction. Arterial blood samples were drawn for radiometabolite analysis, plasma free fraction determination and radiometabolite-corrected arterial input function. Images were reconstructed via a fully 3D time-of-flight iterative reconstruction algorithm using 3 iterations and 34 subsets. The raw PET data were framed into dynamic series of 6x10, 8x15, 6x30, 8x60, 8x120 and 18x300-s frames. All PET images were corrected for photon attenuation and scatter, system dead time, radioactive decay, random coincident events, and detector inhomogeneity.

During the PET image processing and analyses, MATLAB and FSL^58^ were used for PET data processing and registration, respectively. A 3D structural T1-weighted magnetization-prepared rapid gradient-echo (MEMPRAGE) imaging from a 3T Biograph mMR (Siemens Medical Systems) was used as anatomical reference during the registration to obtain regional time-activity curves^32^.

Regional total volume of distribution (*V_T_*) was estimated for the extracted TACs by reversible one-(1T) and two-(2T) tissue compartment model configurations with the metabolite-corrected arterial plasma input function. The Logan graphical analysis technique was also investigated to generate *V_T_* estimates.

#### Analysis of radiometabolite

The arterial blood sampling and processing of [^18^F]**9** was performed with the same procedures as described previously.^32, 36^ Briefly, twenty-two arterial blood samples were drawn during the 120-min PET scans. The plasma samples were obtained as the supernatants following vigorous centrifugation of the whole-blood samples. The radiometabolism profile of [^18^F]**9** was mapped using the selected plasma samples of 3, 5, 10, 15, 30, 60, 90, and 120 minutes. The percent parent in plasma (%PP) was determined via the same automated column switching radioHPLC system as described for [^18^F]JNJ-46356479 (**6**). Similarly, the plasma free fraction *f*_p_ of [^18^F]**9** was determined via ultracentrifugation.

The concentration of radioactivity in whole-blood (WB) and plasma (PL) was measured in a well counter and was expressed as kBq/cc. The time courses of %PP(t) were fitted to a sum of two decaying exponentials plus a constant. By multiplying the resulting model fits with the time course of total plasma radioactivity concentration, the metabolite corrected arterial input function was generated for tracer kinetic modeling.

## ASSOCIATED CONTENT

### Supporting Information

The Supporting Information is available free of charge on the ACS Publications website at DOI: Molecular formular strings (CSV)

*In vitro* mGluRs binding selectivity for compounds **8**-**13**, molecular docking scores and poses for compounds **8**-**11**, automated radiosynthesis, purification and characterization of [^18^F]**8** and [^18^F]**9**, kinetic modeling of [^18^F]**9** in monkey, and ^1^H, ^13^C NMR spectra for synthesized compounds (PDF)

## AUTHOR INFORMATION

### Author Contributions

The manuscript was written through contributions of all authors.

### Notes

The authors declare no conflict interest.

## ACKNOWLEDGEMENT

This project was supported by NIH grants [R01EB021708, R01NS100164, 1S10RR023452-01 and 1S10OD025234-01] for development and characterization of the imaging ligands and by the NIH grants [S10OD018035 and P41EB022544] for the blood counting and metabolite analysis equipment. The functional data were generously provided by the National Institute of Mental Health’s Psychoactive Drug Screening Program, Contract # HHSN-271-2013-00017-C (NIMH PDSP). This program is directed by Bryan L. Roth (mail to) at the University of North Carolina at Chapel Hill and Project Officer Jamie Driscoll (mail to) at NIMH, Bethesda MD, USA. The experimental details can be found at the PDSP website (https://pdspdb.unc.edu/pdspWeb/).

## ABBREVIATIONS USED

PAM: positive allosteric modulator
7-TM: seven transmembrane
VFTD: Venus flytrap domain
PDB: Protein Data Bank
BBB: blood-brain barrier
RCY: radiochemical yield
A_m_: molar activity
Bpin: boronic acid pinacol
TEAB: tetraethyl ammonium bicarbonate
DMA: dimethylacetamide
EOS: end of synthesis
LCMS: Liquid chromatography-mass spectrometry
TAC: time-activity curve
*F_p_*: plasma free fraction
WB: whole-blood
PL: plasma
AIC: Akaike information criteria
NMT: NIMH macaque template
% ID/g: percentage of injected dose per gram of wet tissue
SUV: standardized uptake value
*V_T_*: regional total volume of distribution
MEMPRAGE: magnetization-prepared rapid gradient-echo
DMEM: Dulbecco’s modified Eagle’s medium
PLL: poly-L-lysine
BME: Basal Medium Eagle
USP: United States Pharmacopeia
MLEM: maximum-likelihood expectation-maximization

