## Supporting information for "Design, synthesis, and characterization of [^18^F]mG2P026 as a high contrast PET imaging ligand for metabotropic glutamate receptor 2"

### Table of Contents

1. *In vitro* mGluRs binding selectivity for compounds **8-13**……………………..……………S2

2. Molecular docking scores and poses for compounds **8-11**………………………...…….…S2

3. Automated synthesis, purification and characterization of [$^{18}$F]**8** and [$^{18}$F]**9** …...……...S6

4. Kinetic modeling of [$^{18}$F]**9** in monkey ...…….……………………………...….…S10

5. NMR spectra ……………………..……………………………….....…………….…S14

6. References……………………………...……..……...……....….……...……S48

**1. Table S1**. Binding selectivity of compounds **6** and **8-13** toward other mGluRs

| | mGluR1 | | mGluR2 | mGluR3 | mGluR4 | | mGluR5 | | mGluR6 | | mGluR8 | |
|---|---|---|---|---|---|---|---|---|---|---|---|---|
| **Comp.** | **Ago** | **Anta** | **Anta** | **Anta** | **Ago** | **Anta** | **Ago** | **Anta** | **Ago** | **Anta** | **Ago** | **Anta** |
| **6** | >10 µM | 8.5 µM | >10 µM | ND | >10 µM | >10 µM | >10 µM | 15.1 µM | >10 µM | >10µM | ND | >10 µM |
| **8** | >10 µM | >10 µM | >10 µM | >10 µM | >10 µM | >10 µM | >10 µM | >10 µM | >10 µM | >10 µM | >10 µM | >10 µM |
| **9** | >10 µM | >10 µM | >10 µM | >10 µM | >10 µM | >10 µM | >10 µM | >10 µM | >10 µM | >10 µM | >10 µM | >10 µM |
| **10** | >10 µM | >10 µM | >10 µM | >10 µM | >10 µM | >10 µM | >10 µM | >10 µM | >10 µM | >10 µM | >10 µM | >10 µM |
| **11** | >10 µM | >10 µM | >10 µM | >10 µM | >10 µM | >10 µM | >10 µM | >10 µM | >10 µM | >10 µM | >10 µM | >10 µM |
| **12** | >10 µM | >10 µM | >10 µM | >10 µM | >10 µM | >10 µM | >10 µM | >10 µM | >10 µM | >10 µM | >10 µM | >10 µM |
| **13** | >10 µM | >10 µM | >10 µM | >10 µM | 1.8 µM | >10 µM | >10 µM | >10 µM | >10 µM | >10 µM | 7.4 µM | >10 µM |

Ago: agonist activity; Anta: antagonist activity; The $EC_{50}$ value of agonist activity or the $IC_{50}$ value of antagonist activity >10 µM indicates that no curve was noted in the dose-response up to 10 µM; ND: Not determined.

### 2. Molecular docking scores and poses for compounds 8-11

**Table S2**: Docking scores for compounds **6** and **8-11**

| Compound # | Docking Score (kcal/mol) |
|---|---|
| **Compound 6** | 13.08 |
| **Compound 8** | 11.89 |
| **Compound 9** | 13.20 |
| **Compound 10** | 14.46 |
| **Compound 11** | 11.35 |

**(a) Compound 8**

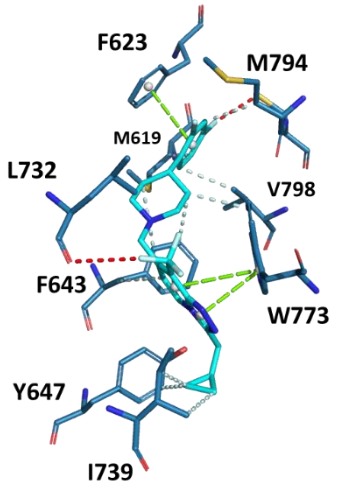


**(b) Compound 9**

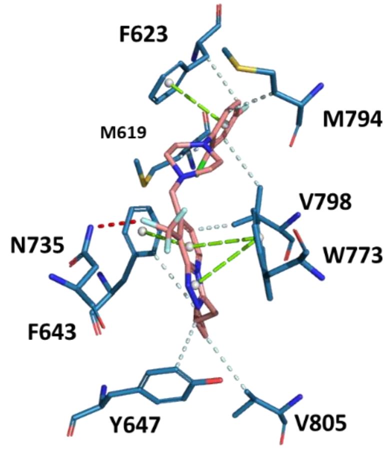


**(c) Compound 10**

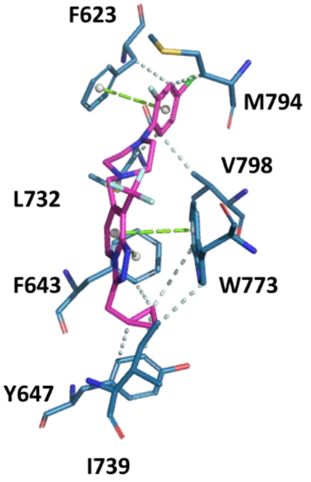


**(d) Compound 11**

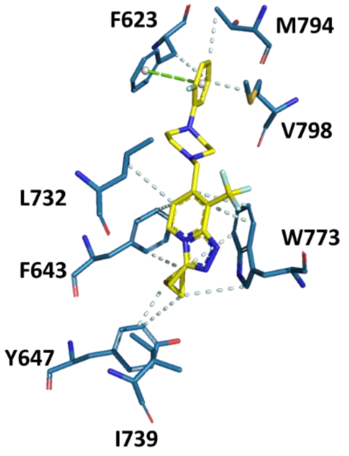

**(e) Compound 6**

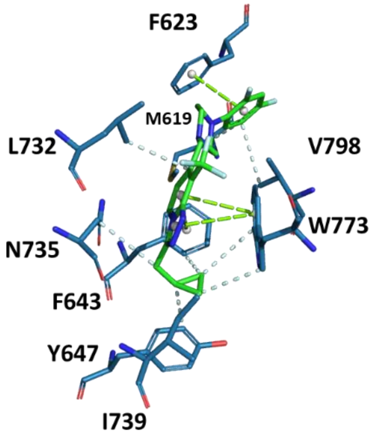


**Figure S1**. Ligand-protein interaction diagram of compounds **8-11** (**a-d**) in comparison with compound **6** (**e**). Pictures were rendered in Protein Ligand Interaction Profiler (PLIP) using Pymol 2.3.3. The binding pocket residues are shown in teal with oxygen in red, nitrogen in blue and ligands with carbons in cyan (**a**), orange (**b**), magenta (**c**), yellow (**d**) and green (**e**) oxygen in red, nitrogen in blue and fluorine in light cyan and chlorine in green. Light cyan dotted lines show hydrophobic interactions, green dotted lines show π-π stacking and red dotted lines show halogen bonds.

### 3. RADIOCHEMISTRY

#### Isolation and analysis of [$^{18}$F]8 and [$^{18}$F]9:

Both [$^{18}$F]**8** and [$^{18}$F]**9** were isolated via the same semi-Prep HPLC conditions: mobile phase: 55% $CH_3CN$ in 0.1M AMF water, 6 mL/min, Waters Xbridge BEH C18, 5 μm, 10 x 250 mm.

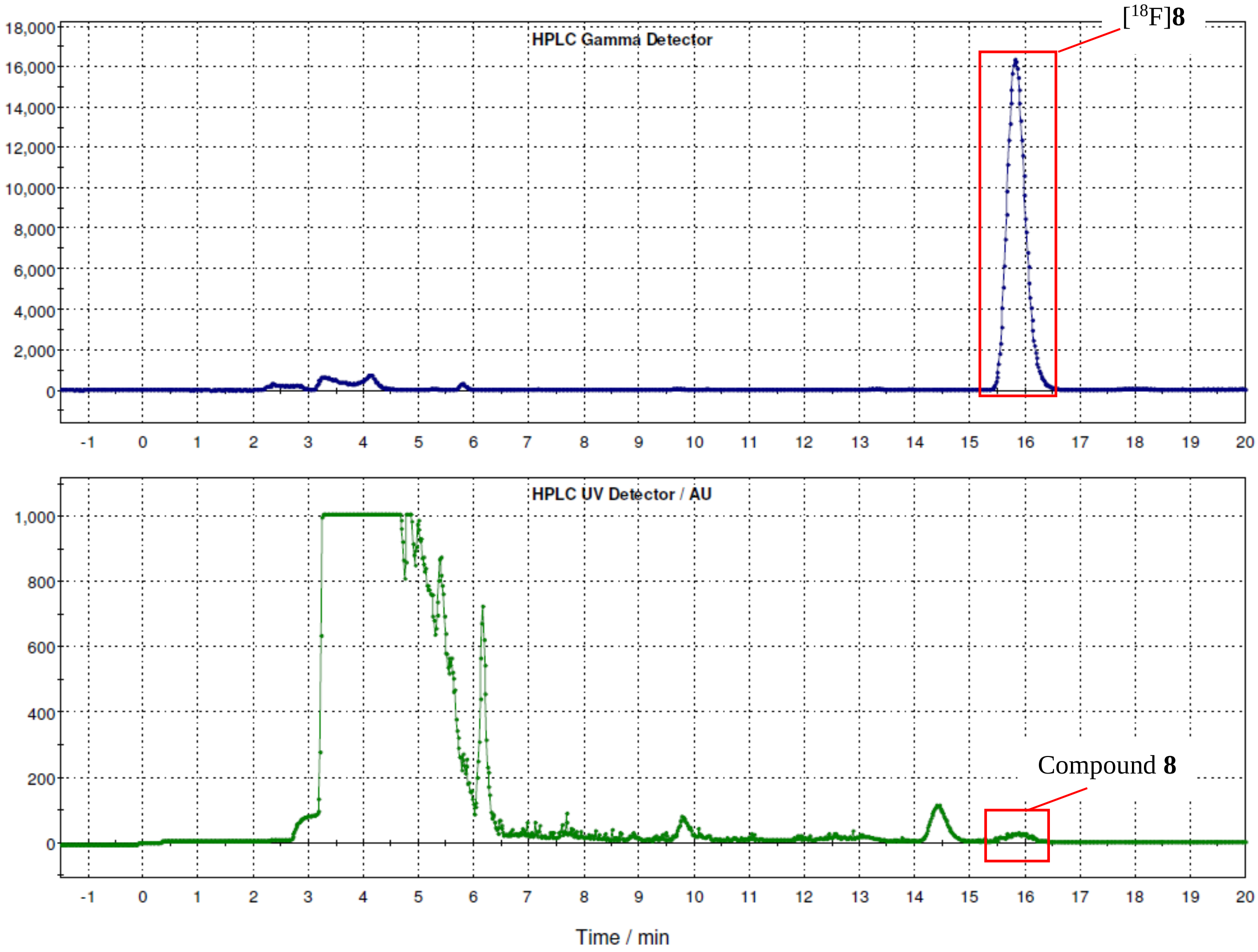


**Figure S2**. Purification chromatography of [$^{18}$F]**8** from the reaction mixture via semi-preparative HPLC.

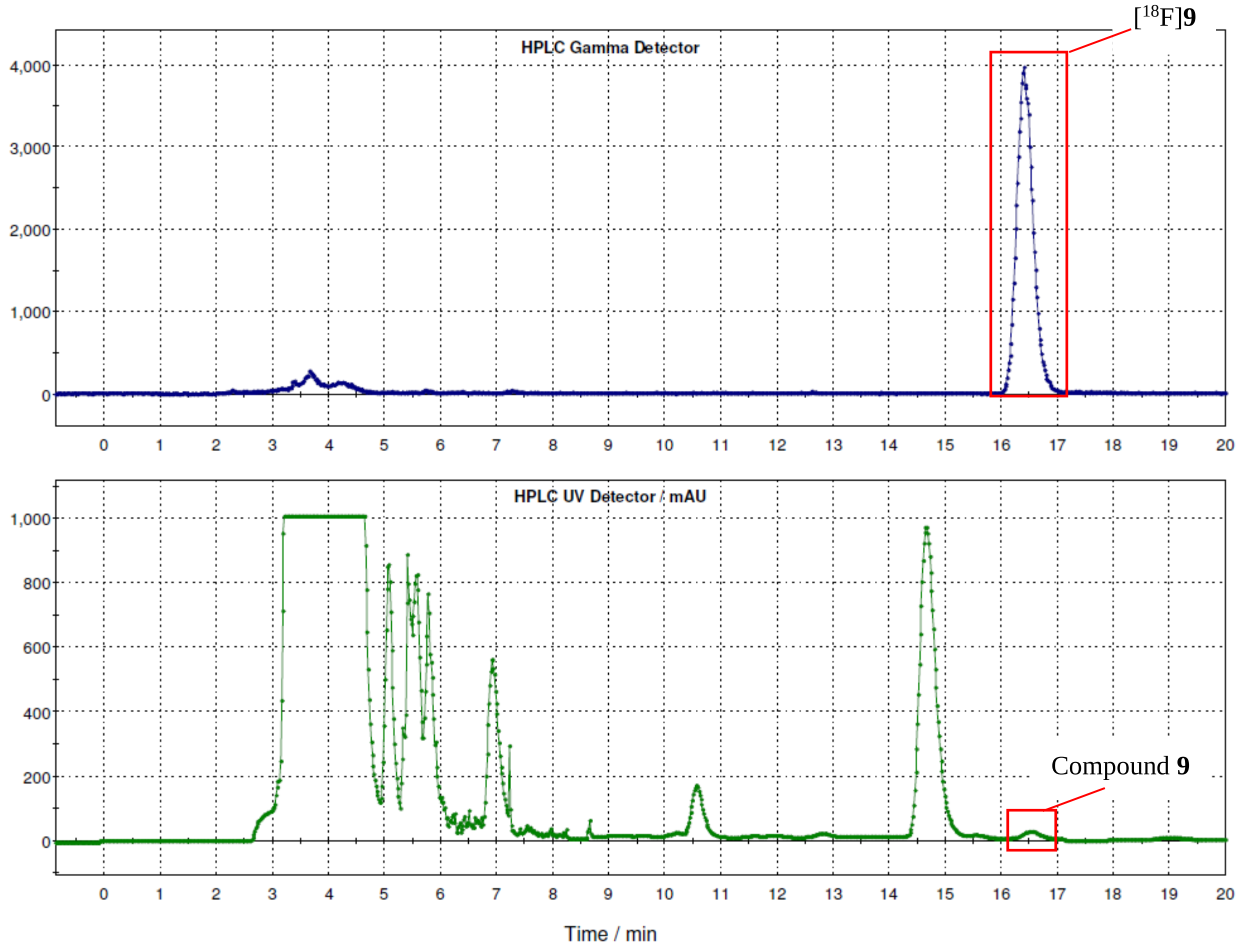


**Figure S3**. Purification chromatography of [$^{18}$F]**9** from the reaction mixture via semi-preparative HPLC.

The [18F]**8** and [18F]**9** were analyzed via the same analytical HPLC conditions: mobile phase: 60% $CH_3CN$ in 0.1M AMF water, 1 mL/min, Waters Xbridge BEH C18, 3.5 μm, 4.6 x 150 mm.

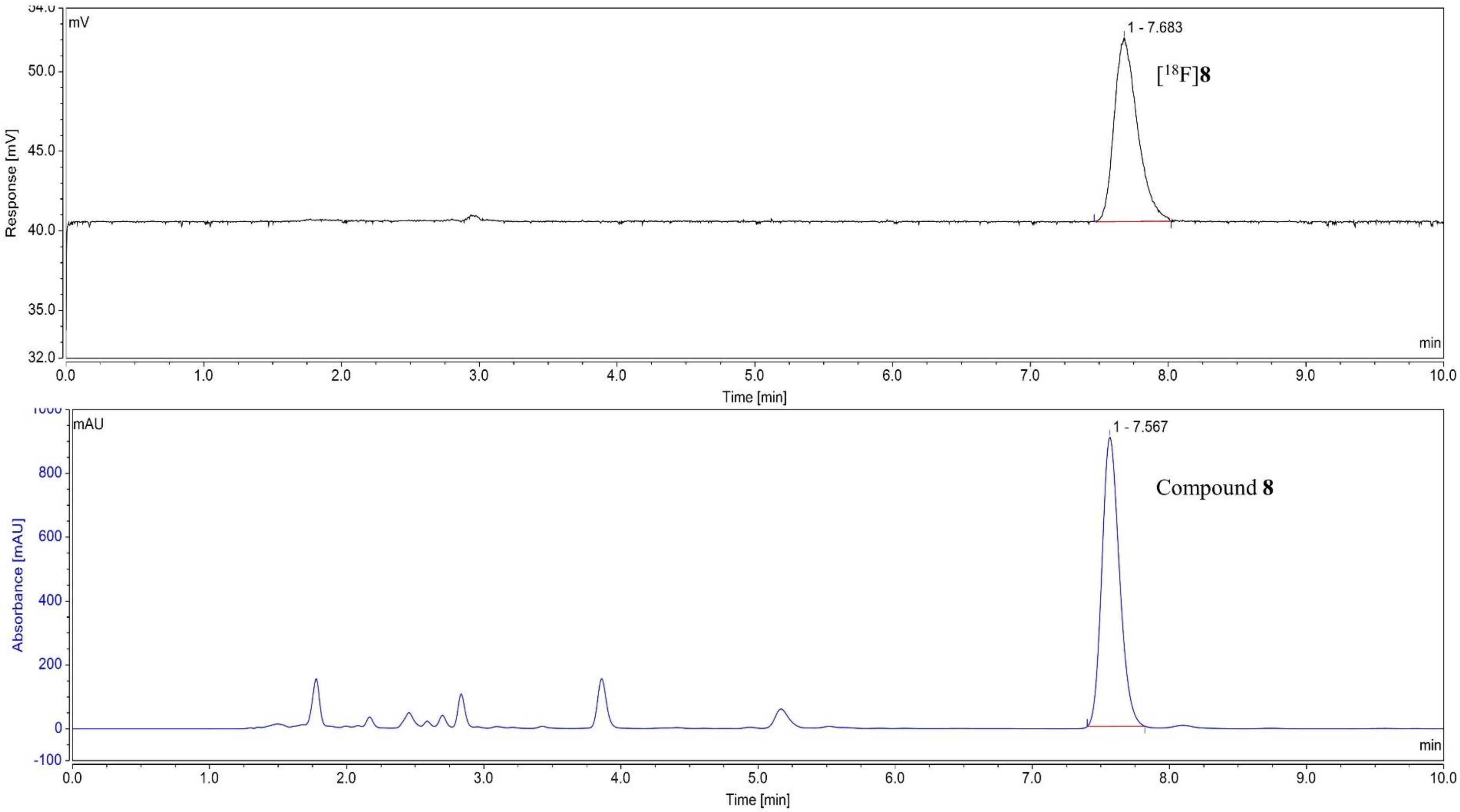


**Figure S4**. Confirmation of the [18F]**8** via HPLC analysis. The radioHPLC trace is shown in black (top) and the UV trace is shown in blue (bottom) under a wavelength of 254 nM.

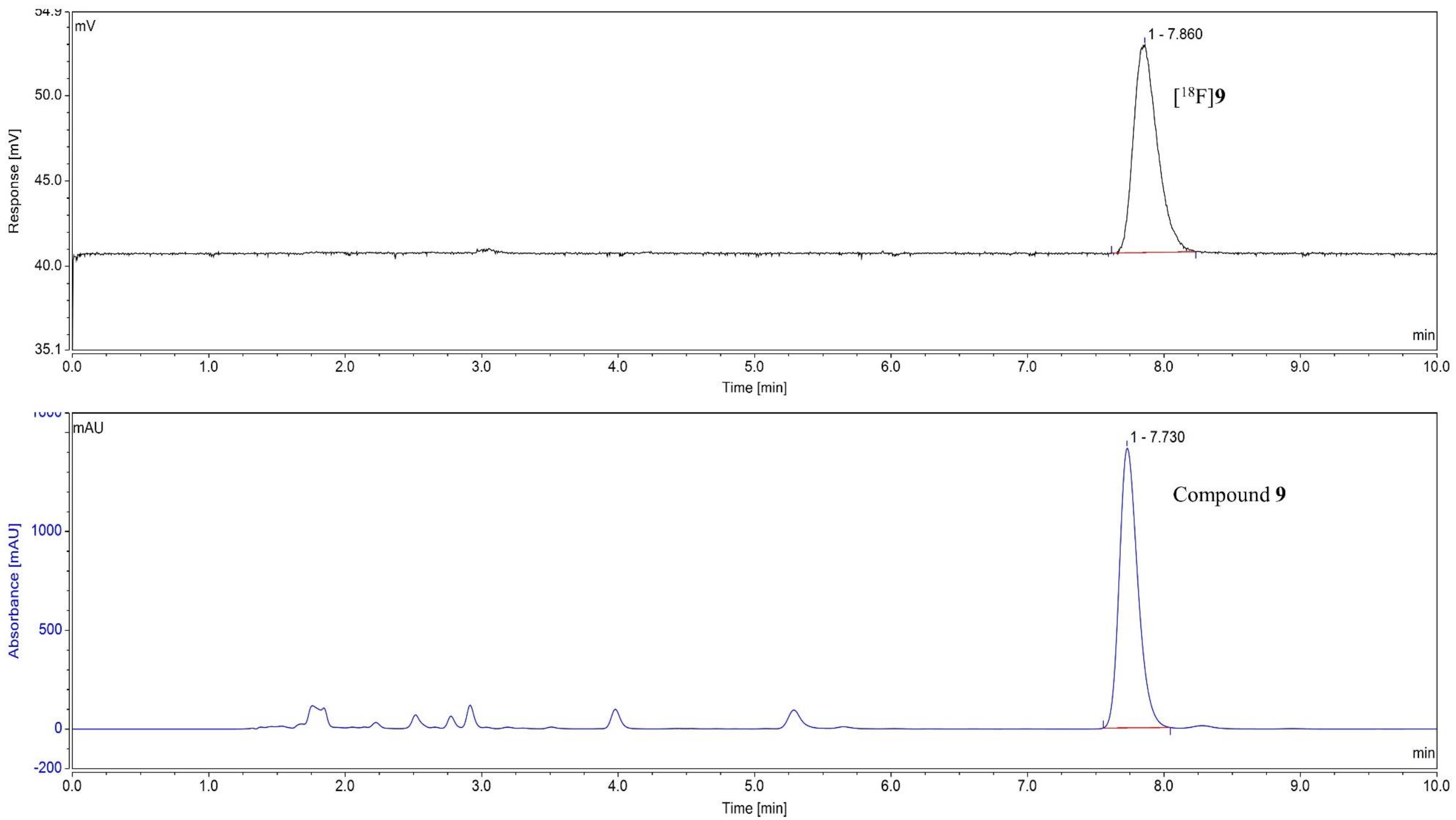


**Figure S5**. Confirmation of the [$^{18}$F]**9** via HPLC analysis. The radioHPLC trace is shown in black (top) and the UV trace is shown in blue (bottom) under a wavelength of 254 nM.

### 4. Kinetic modeling of [$^{18}$F]9 in monkey

**Table S3.** 2T4k1v model parameters for baseline study

| Regions | $K_1$ mL/min/cc | $k_2$ ($min^{-1}$) | $k_3$ ($min^{-1}$) | $k_4$ ($min^{-1}$) | $V_f$ | $V_T$ (mL/cc) |
|---|---|---|---|---|---|---|
| Whole Brain | 0.232682 | 0.07077 | 0.016874 | 0.021486 | 0.131423 | 5.870075 |
| Caudate | 0.243262 | 0.065665 | 0.012053 | 0.021812 | 0.1248 | 5.751756 |
| Cerebellum | 0.330021 | 0.089772 | 0.01226 | 0.024191 | 0.201576 | 5.539202 |
| Cortex all | 0.251348 | 0.070741 | 0.012762 | 0.023429 | 0.129177 | 5.488489 |
| Thalamus | 0.264995 | 0.068123 | 0.031486 | 0.045743 | 0.103222 | 6.567432 |
| Striatum | 0.309946 | 0.072011 | 0.011109 | 0.023213 | 0.16875 | 6.364027 |
| White Matter | 0.185404 | 0.060126 | 0.02847 | 0.023947 | 0.110975 | 6.749583 |

**Table S4.** $V_T$ estimates using Logan graphical method for baseline study

| Regions | $V_T$ t*=30 min (mL/cc) |
|---|---|
| Whole Brain | 5.0281 |
| Caudate | 4.9896 |
| Cerebellum | 4.6049 |
| Cortex all | 4.7932 |
| Thalamus | 6.0173 |
| Striatum | 5.3553 |
| White Matter | 5.8696 |

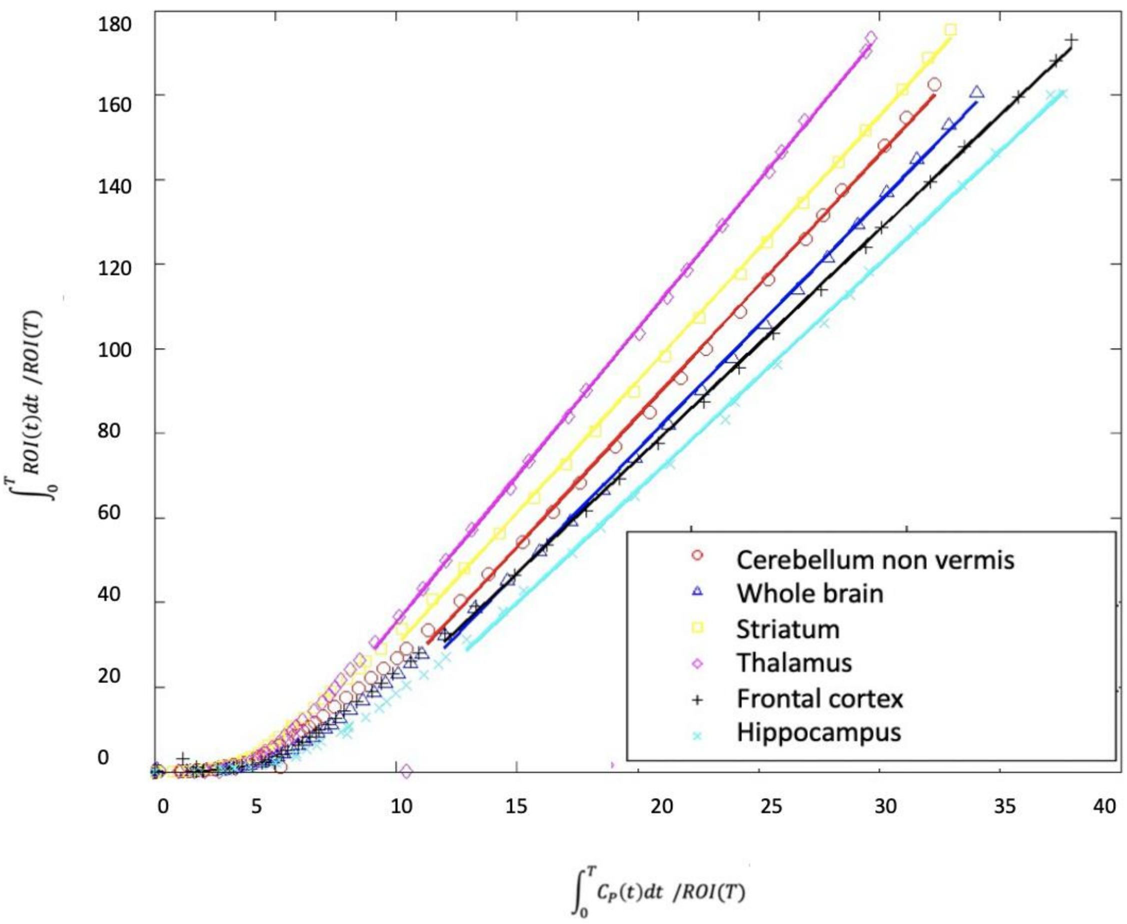


**Figure S6**. Moolyg baseline Logan model fits (30 min t*)

**a**

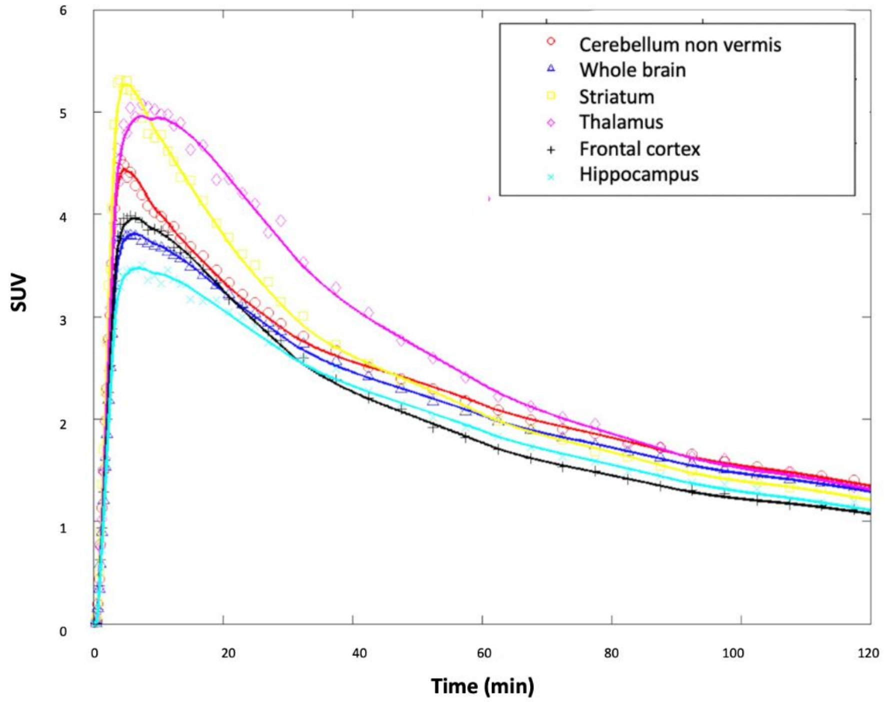


**b**

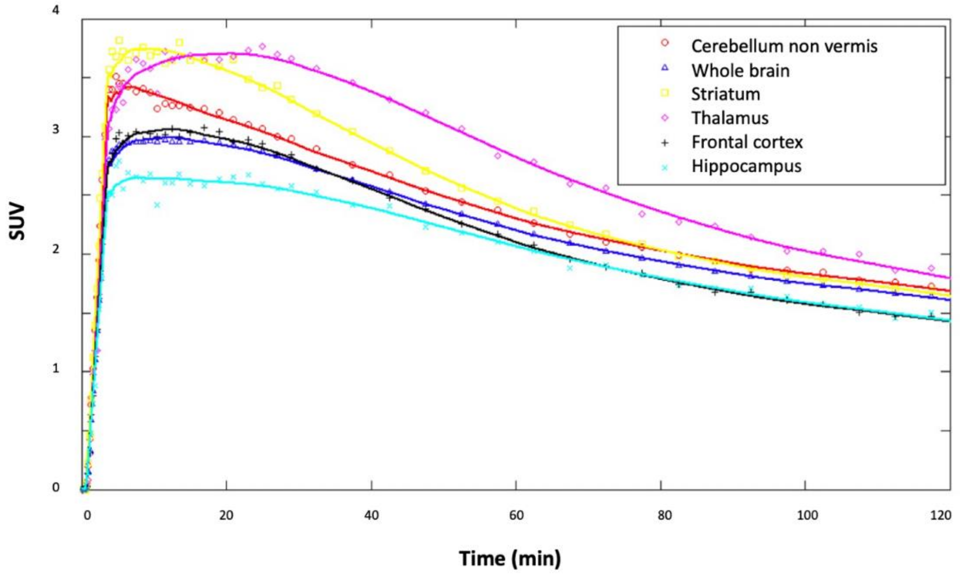


**Figure S7**. Comparison of the time-activity curve between [$^{18}$F]JNJ-46356479 (**6**, top) and [$^{18}$F]mG2P026 (**9**, bottom).

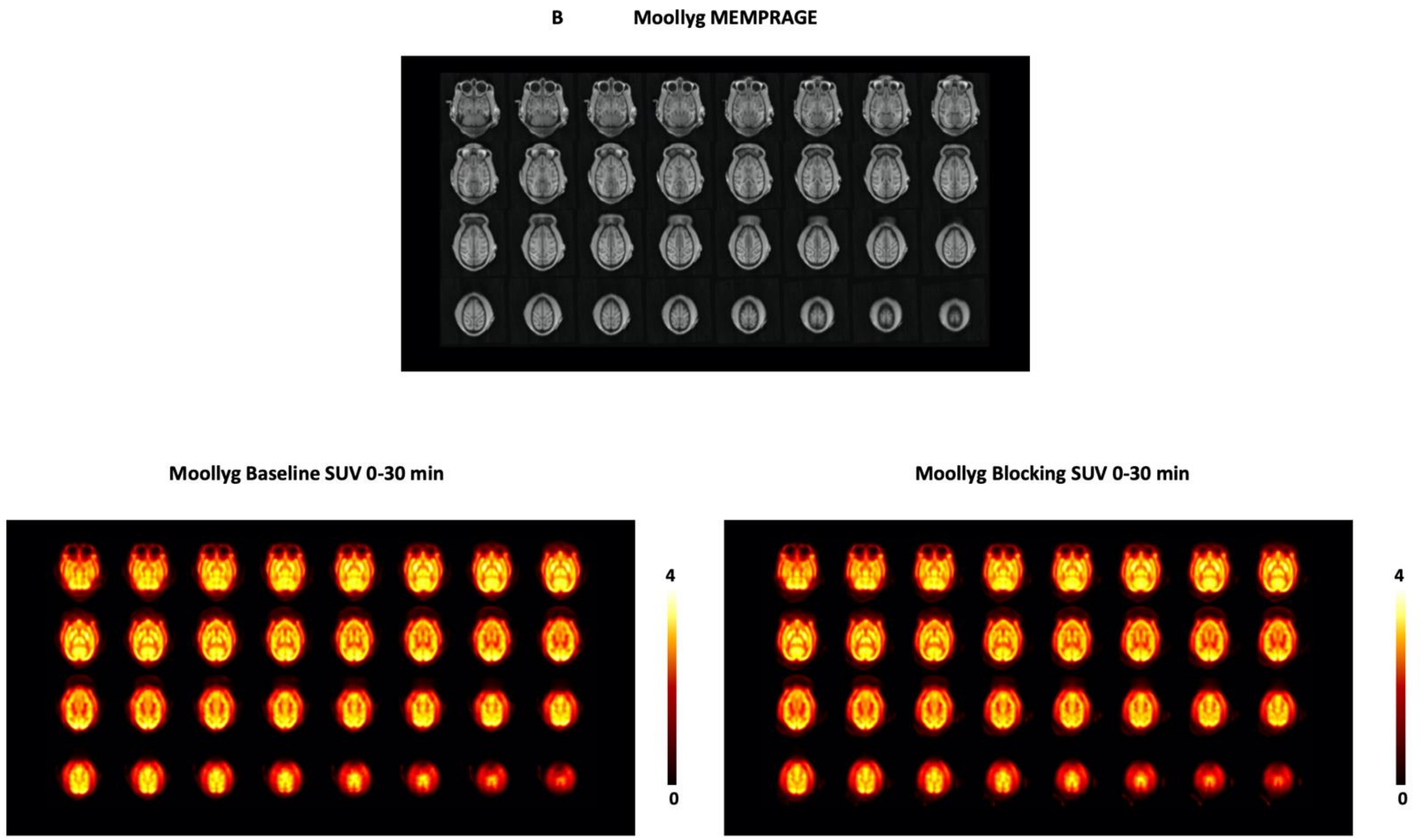


**Figure S8**. Structural MRI (MEMPRAGE) and [[18]F]**9** SUV images calculated from 0 to 30 min post tracer injection ($SUV_{0\text{-}30min}$) for the baseline (left) and blocking conditions (right). Images are presented in the NIMH Macaque Template (NMT)[1] space.

### 5. NMR spectra

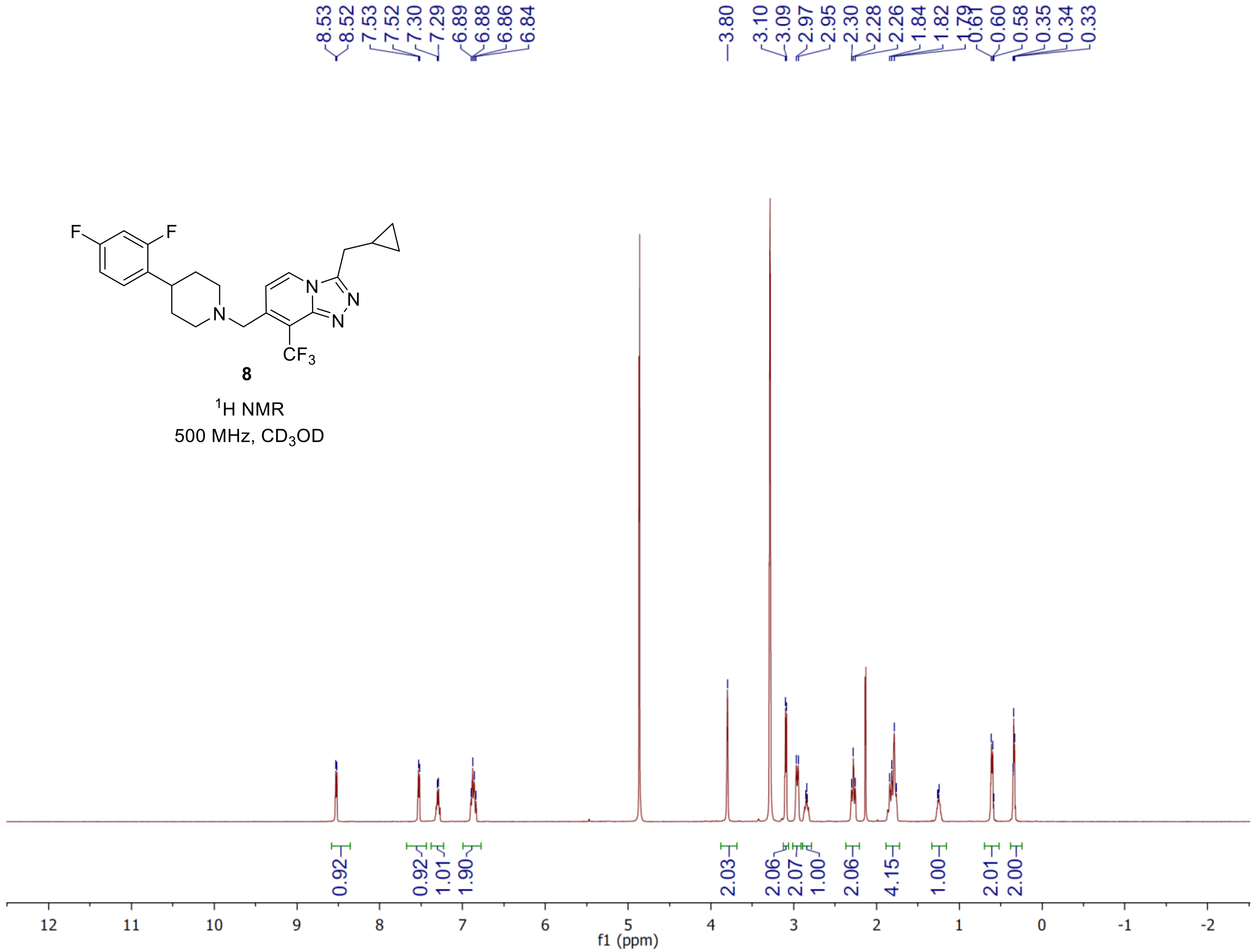

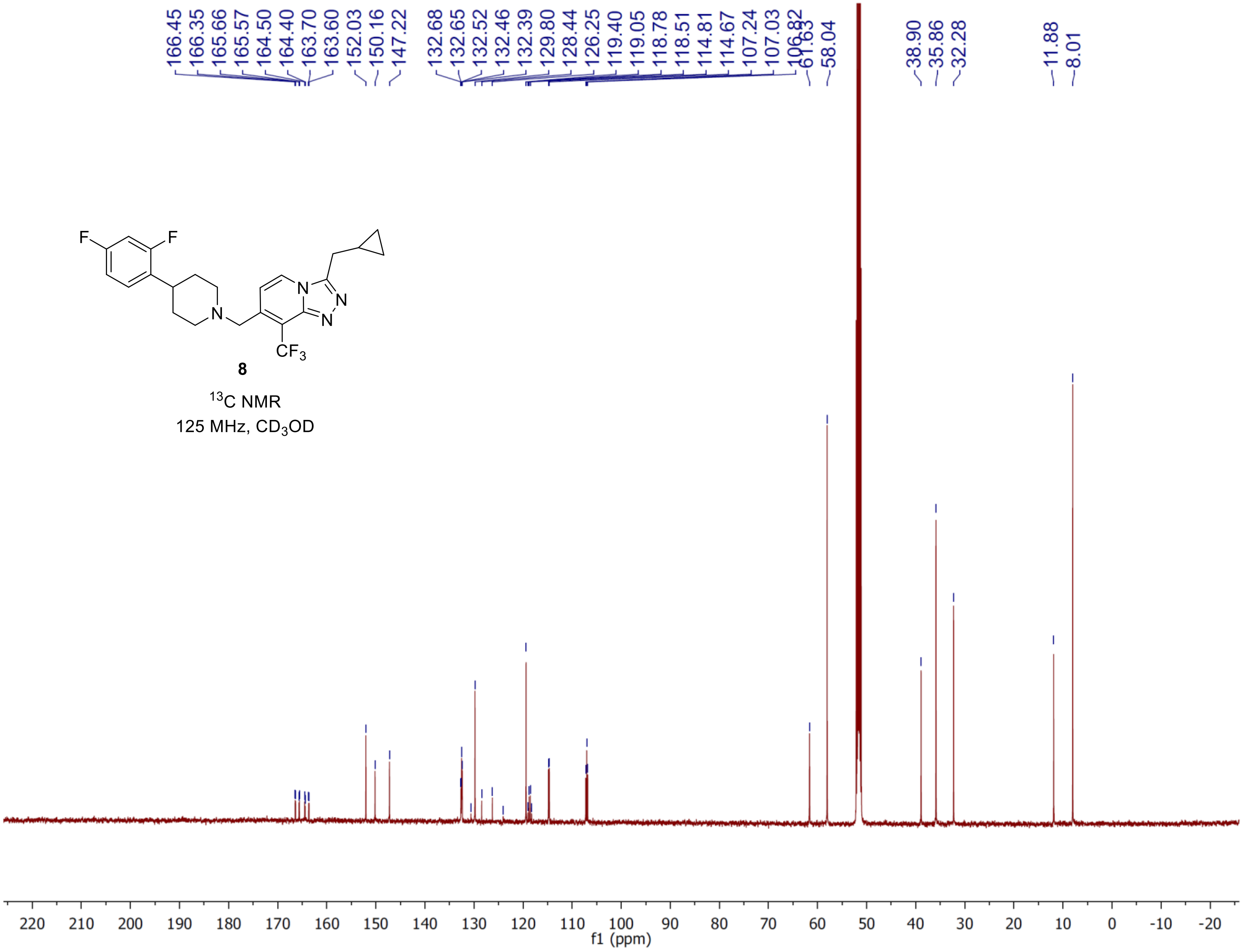
166.45
166.35
165.66
165.57
164.50
164.40
163.70
163.60
152.03
150.16
147.22
132.68
132.65
132.52
132.46
132.39
129.80
128.44
126.25
119.40
119.05
118.78
118.51
114.81
114.67
107.24
107.03
106.82
61.63
58.04
38.90
35.86
32.28
11.88
8.01
F
F
N
N
N
N
CF3
8
13C NMR
125 MHz, CD3OD
220 210 200 190 180 170 160 150 140 130 120 110 100 90 80 70 60 50 40 30 20 10 0 -10 -20
f1 (ppm)

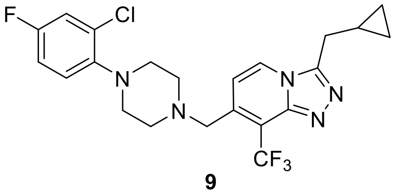


**9**

$^{1}H$ NMR

500 MHz, $CD_3OD$

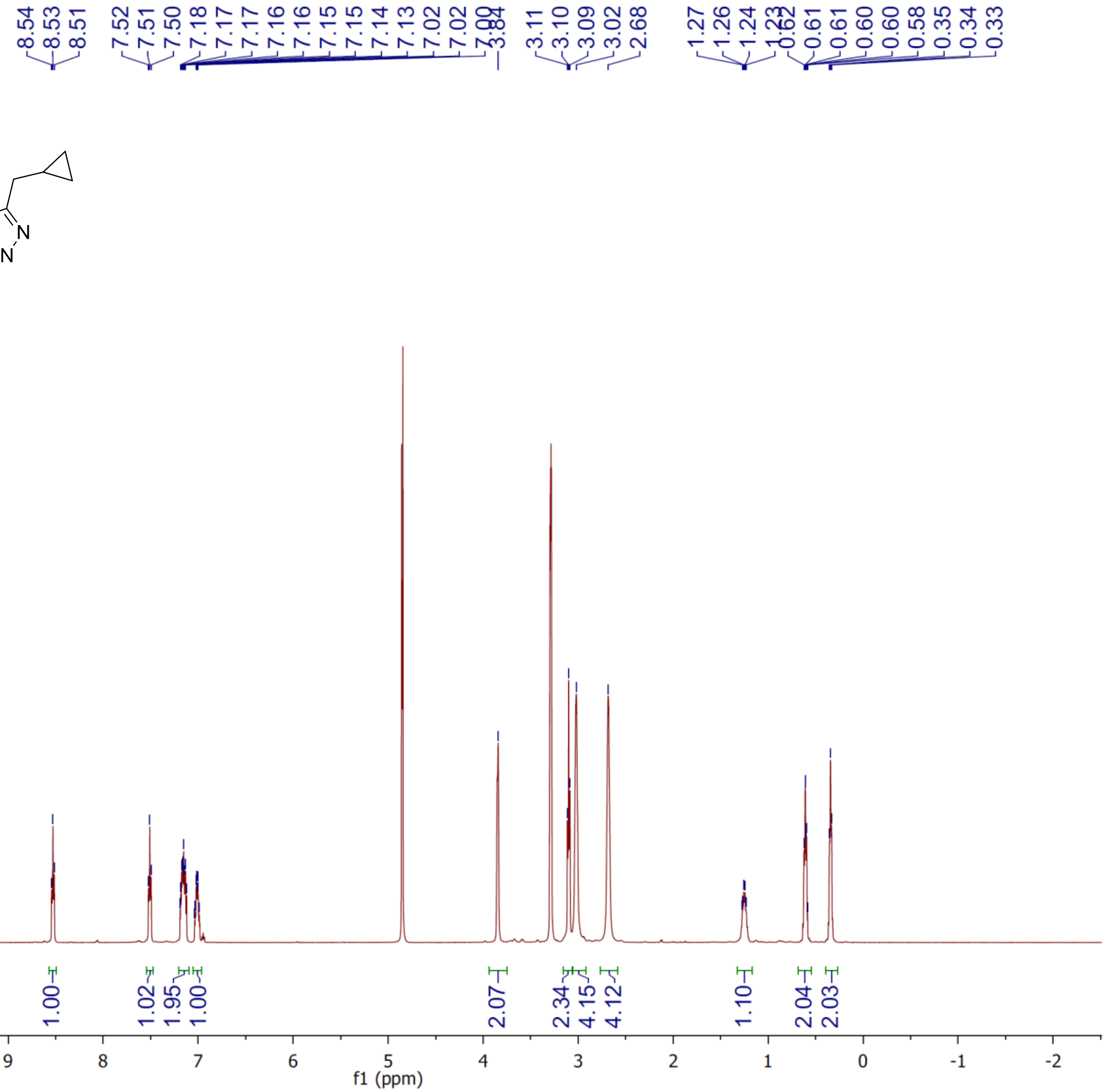

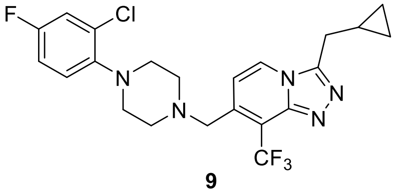


**9**

$^{13}C$ NMR

125 MHz, $CD_3OD$

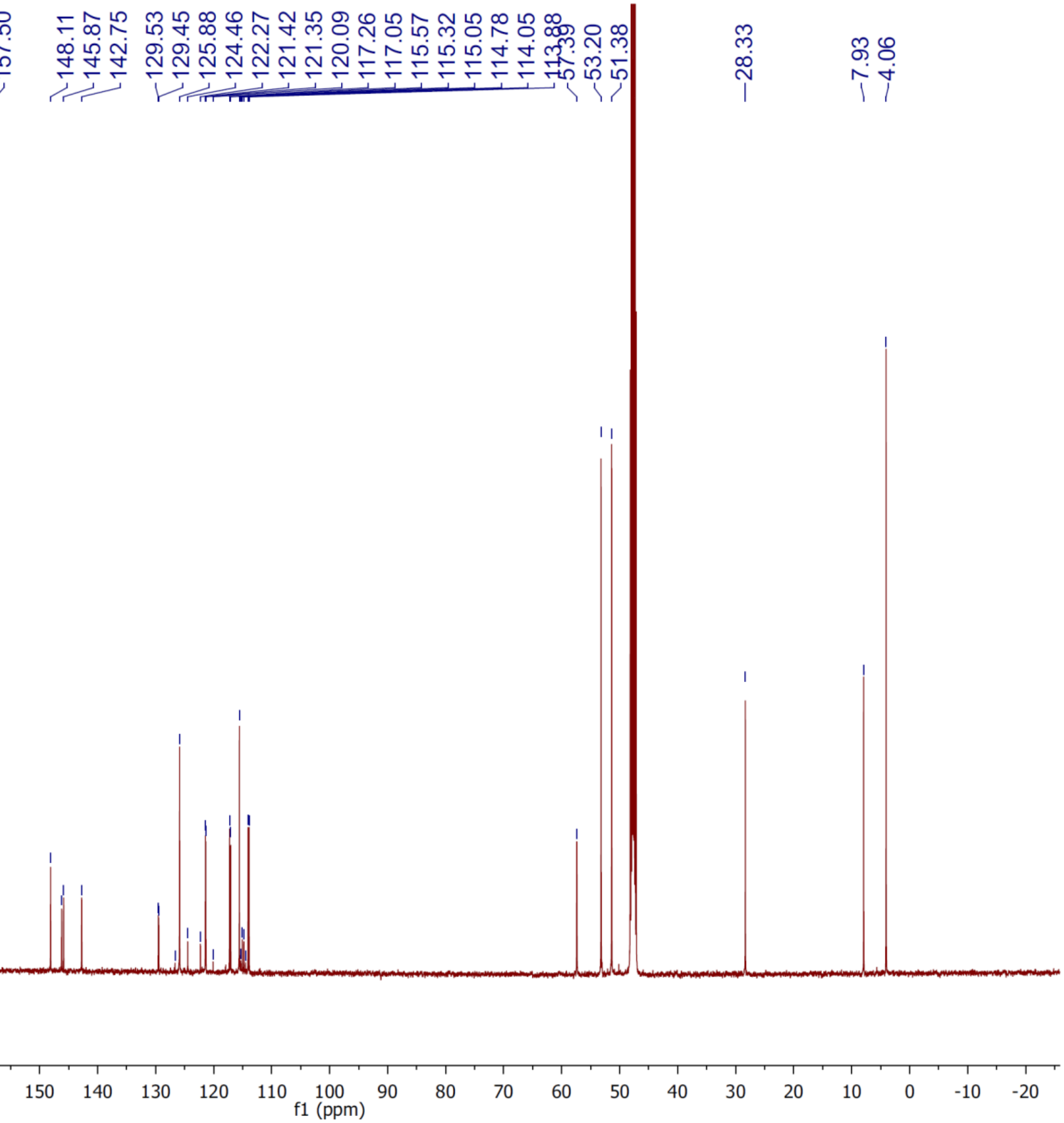

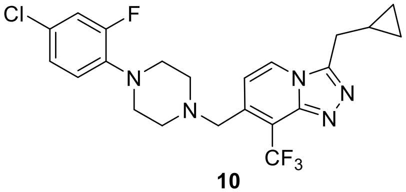


**10**

$^{1}H$ NMR

500 MHz, $CDCl_3$

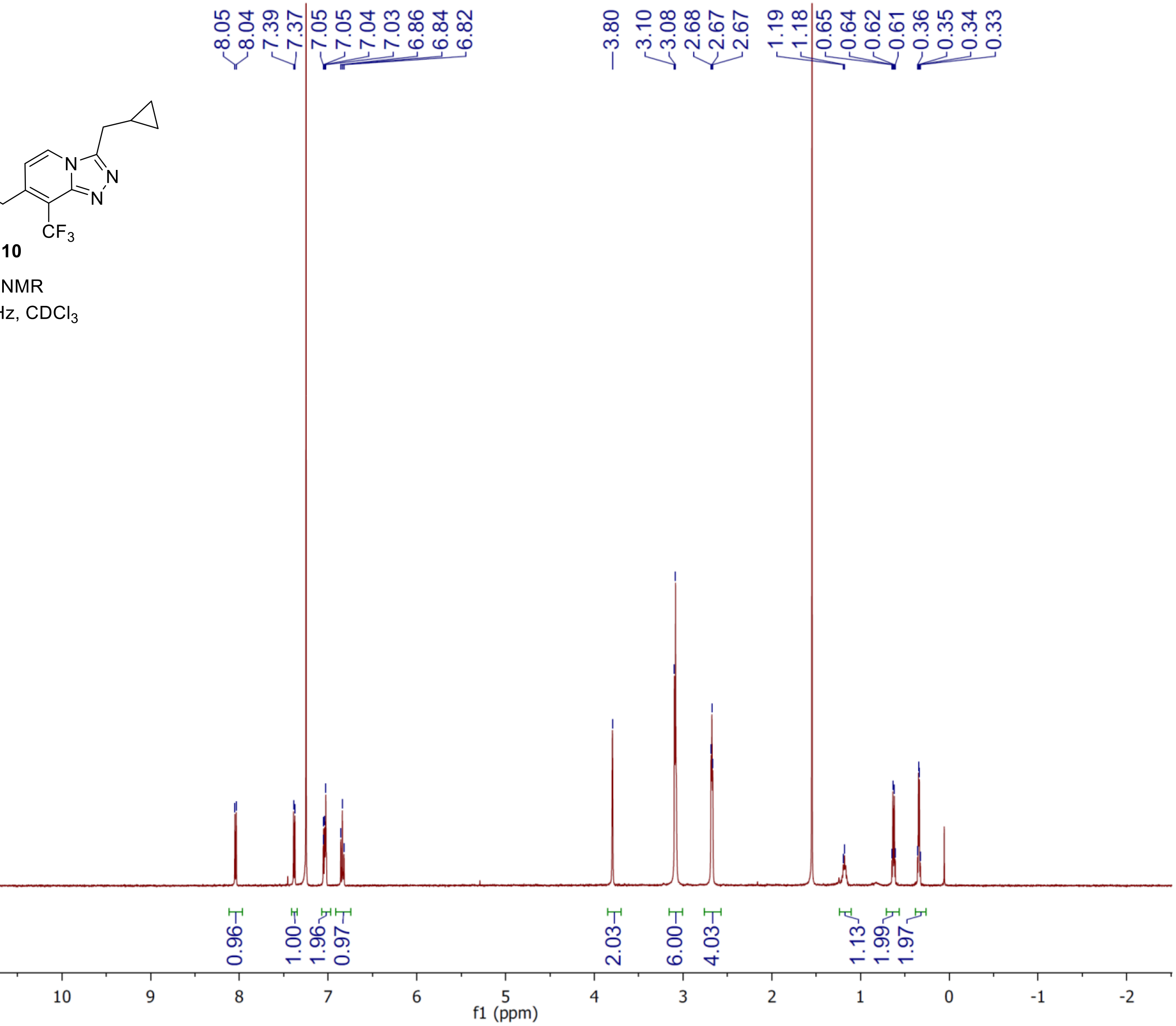

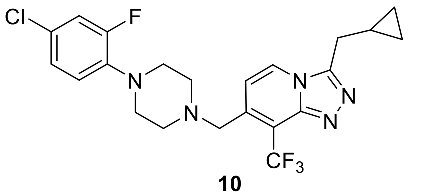

**10**

$^{13}C$ NMR
125 MHz, $CDCl_3$

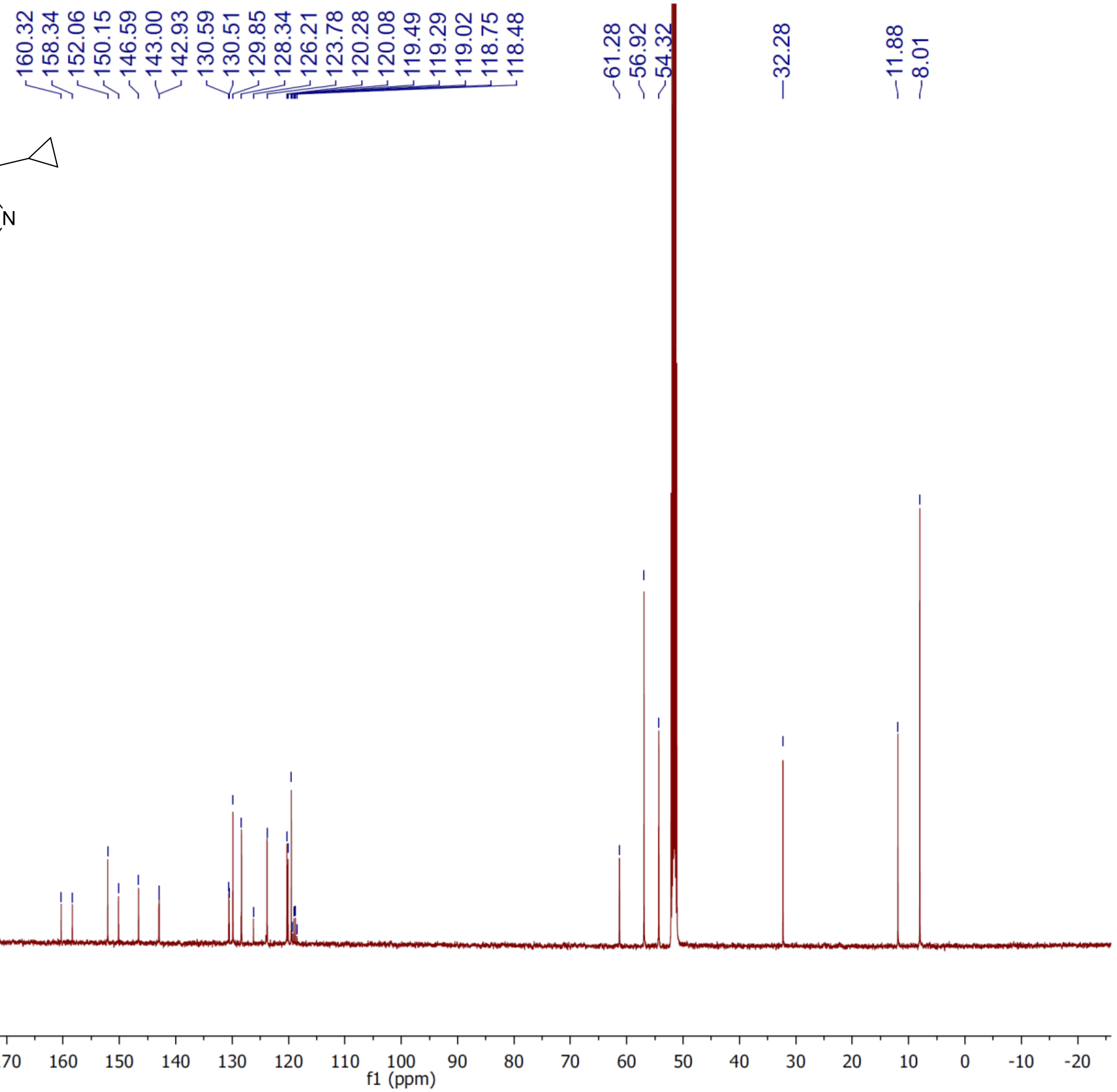

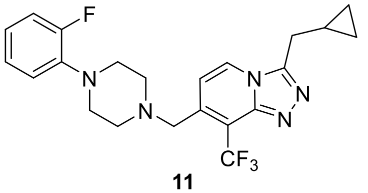


**11**

$^1H$ NMR

500 MHz, $CD_3OD$

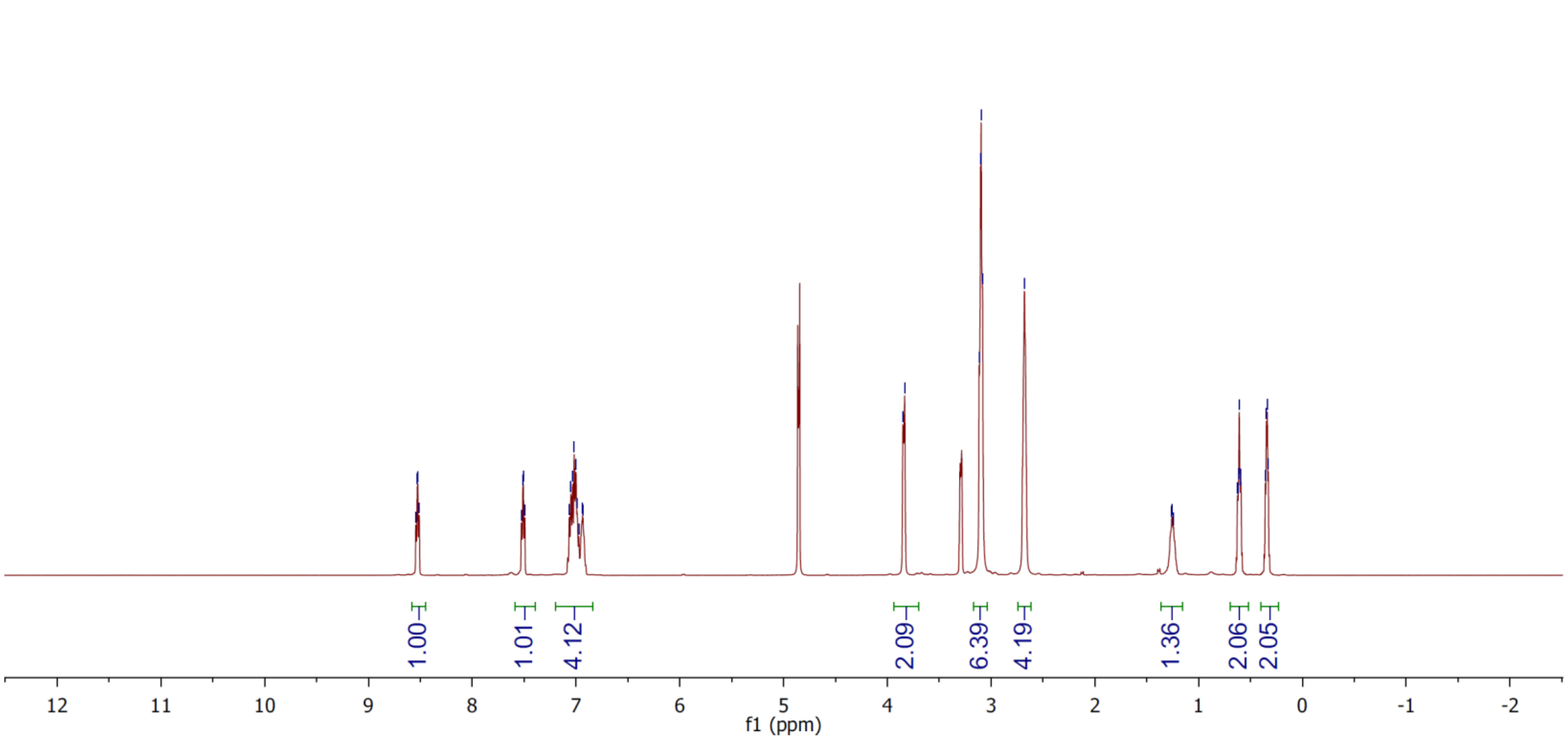

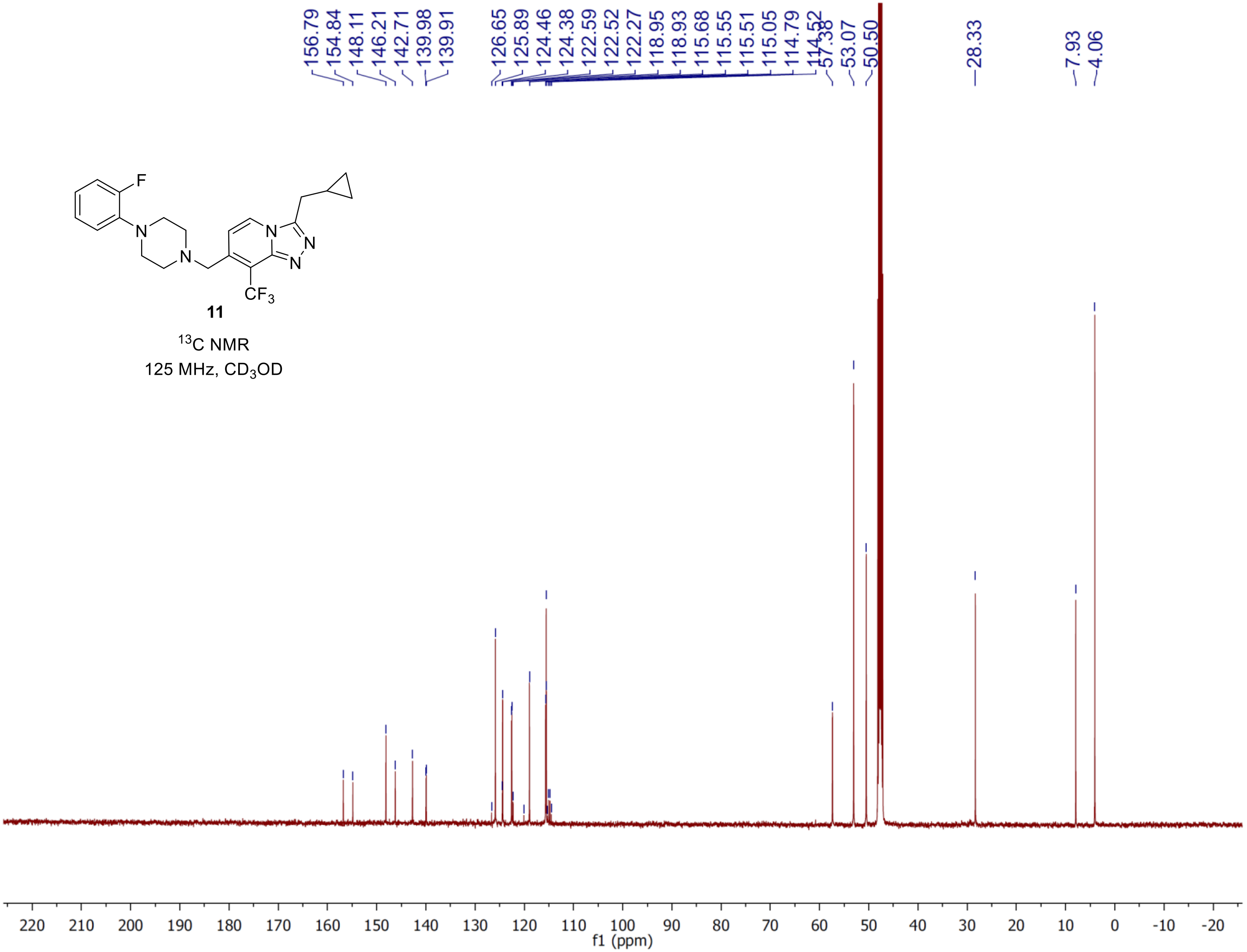


**11**

$^{13}C$ NMR

125 MHz, $CD_3OD$

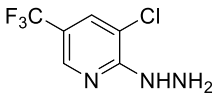


**20**

$^1$H NMR

500 MHz, $CDCl_3$

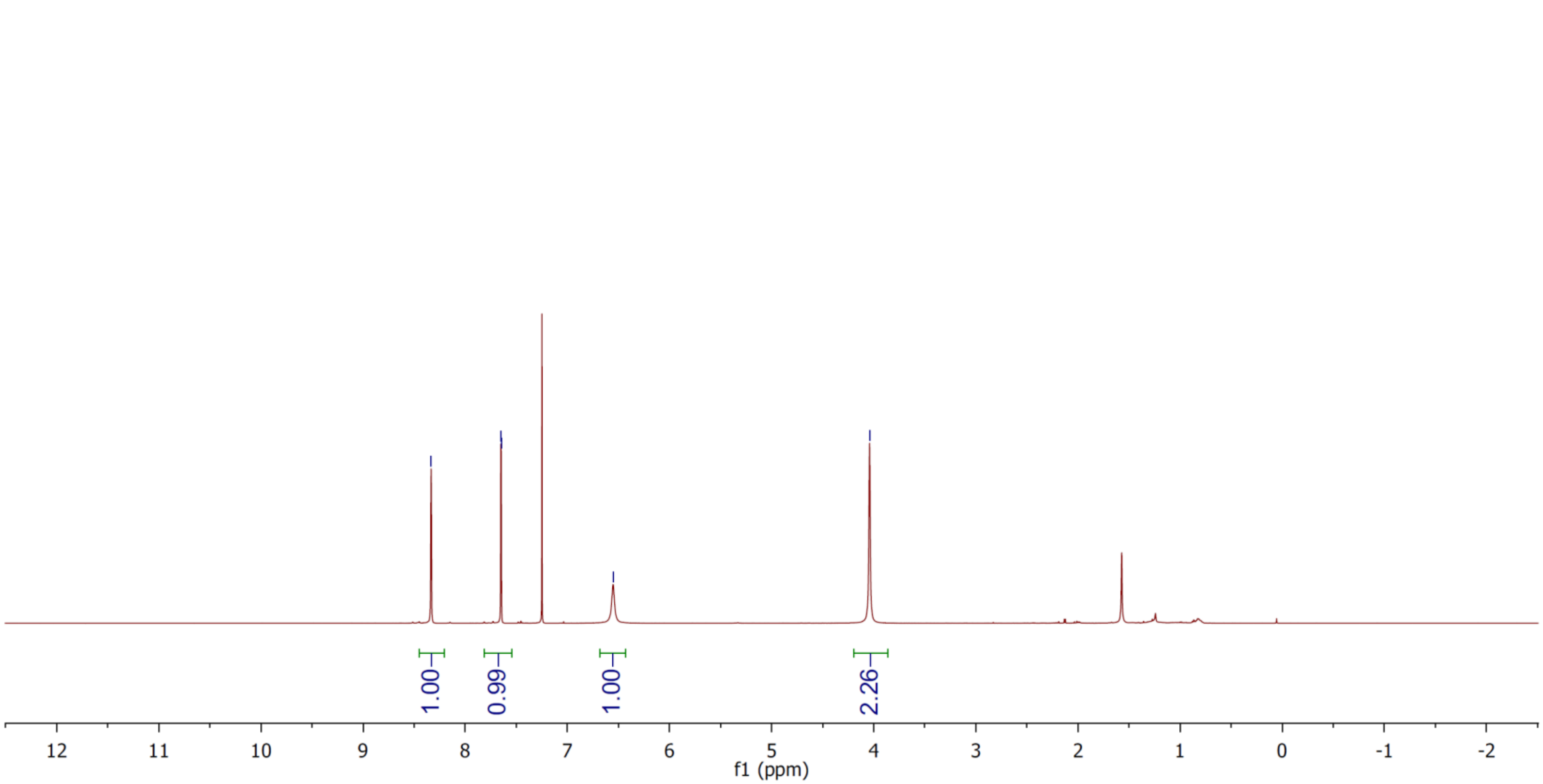

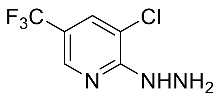

**20**

$^{13}C$ NMR
125 MHz, $CDCl_3$

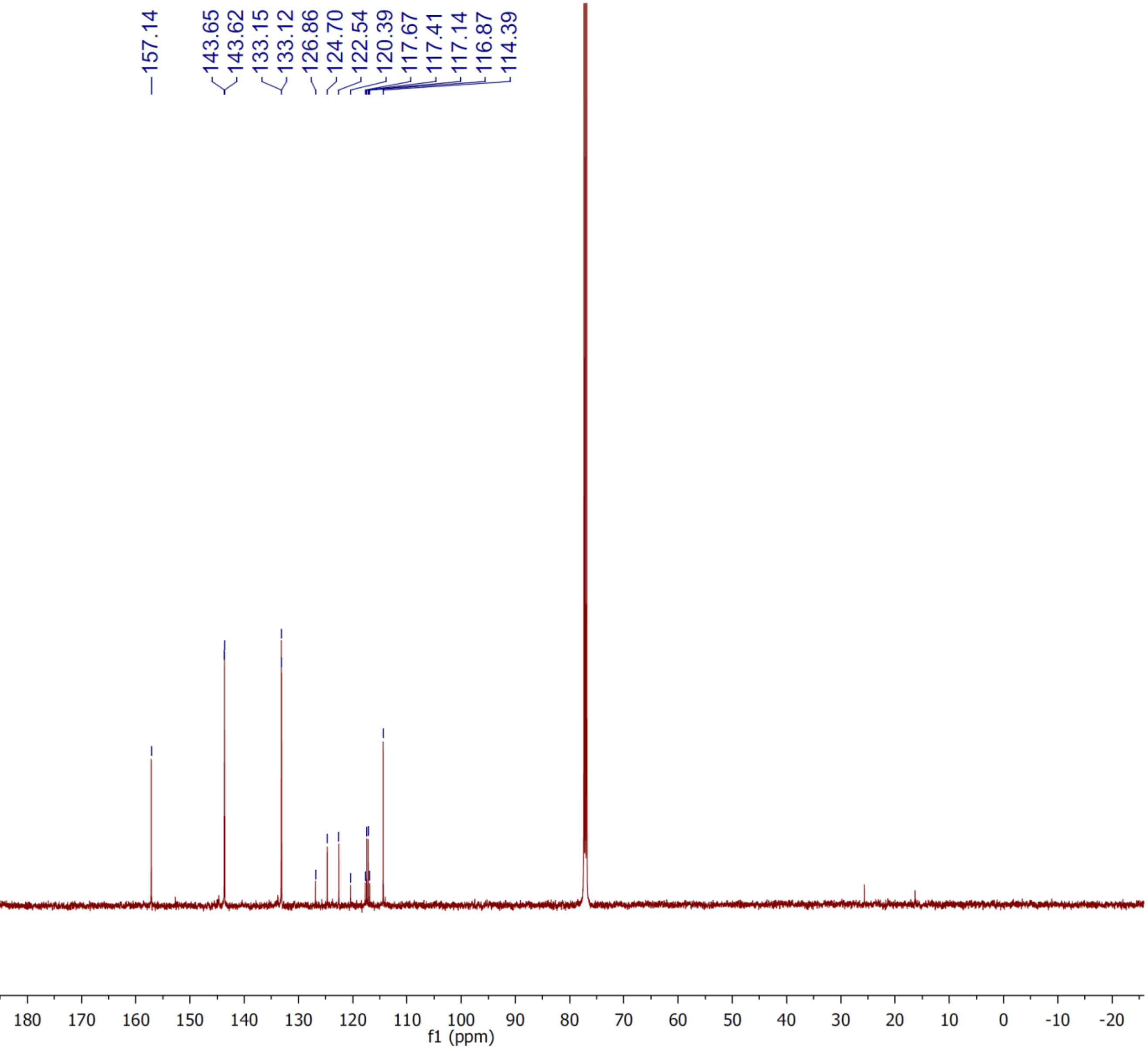

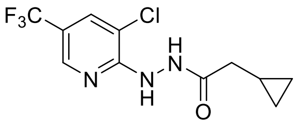


**21**

$^{1}$H NMR

500 MHz, $CDCl_3$

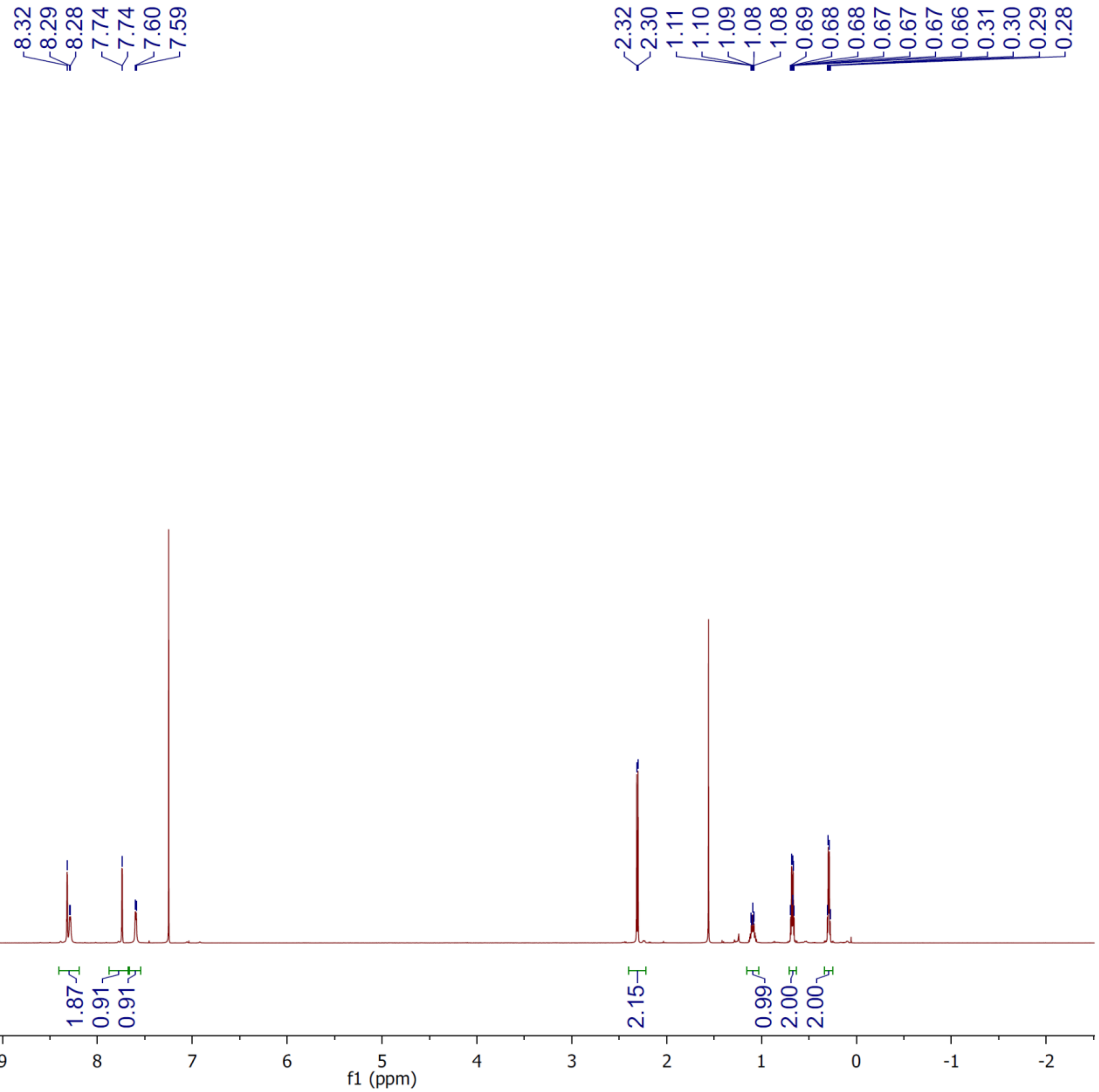

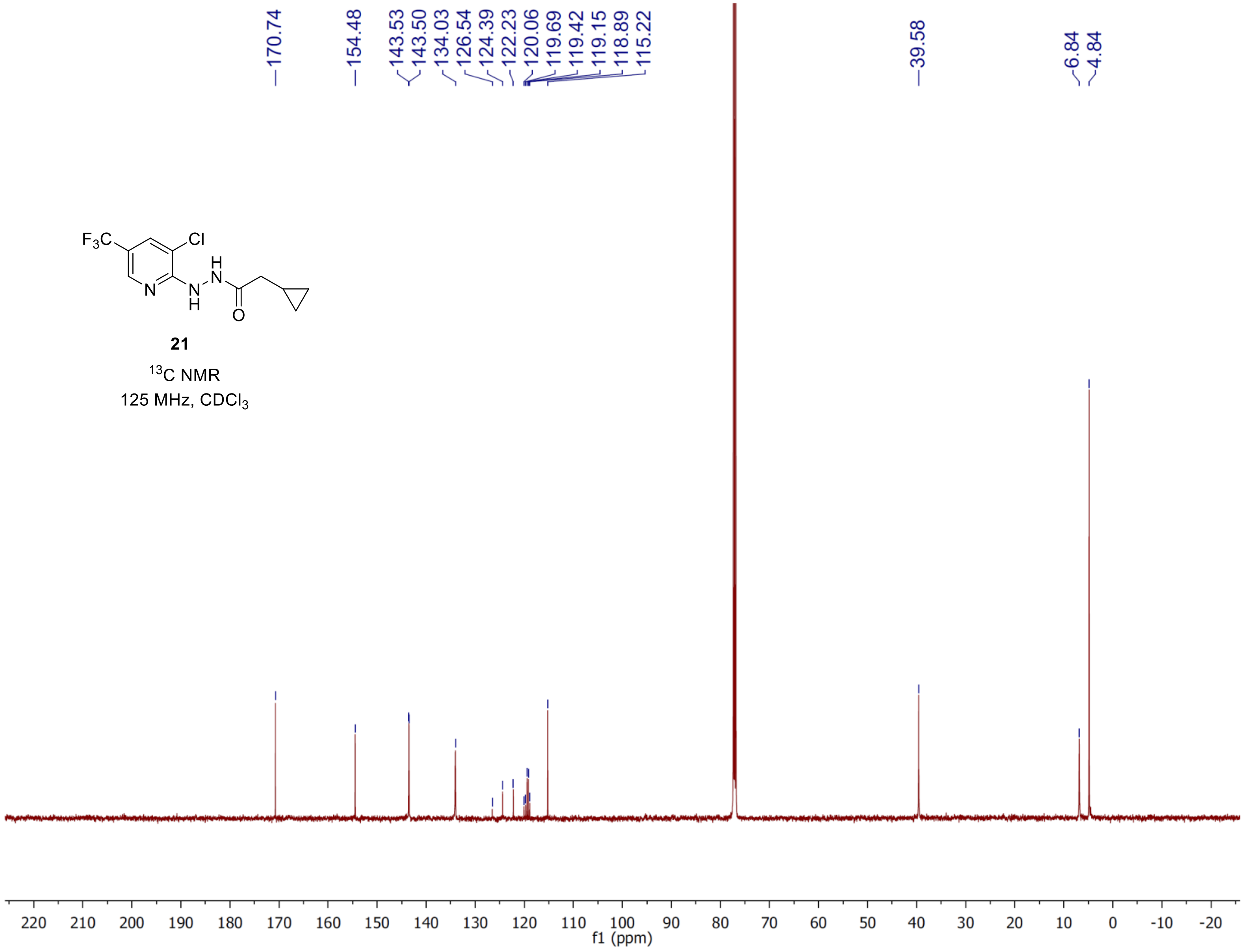


**21**

$^{13}C$ NMR

125 MHz, $CDCl_3$

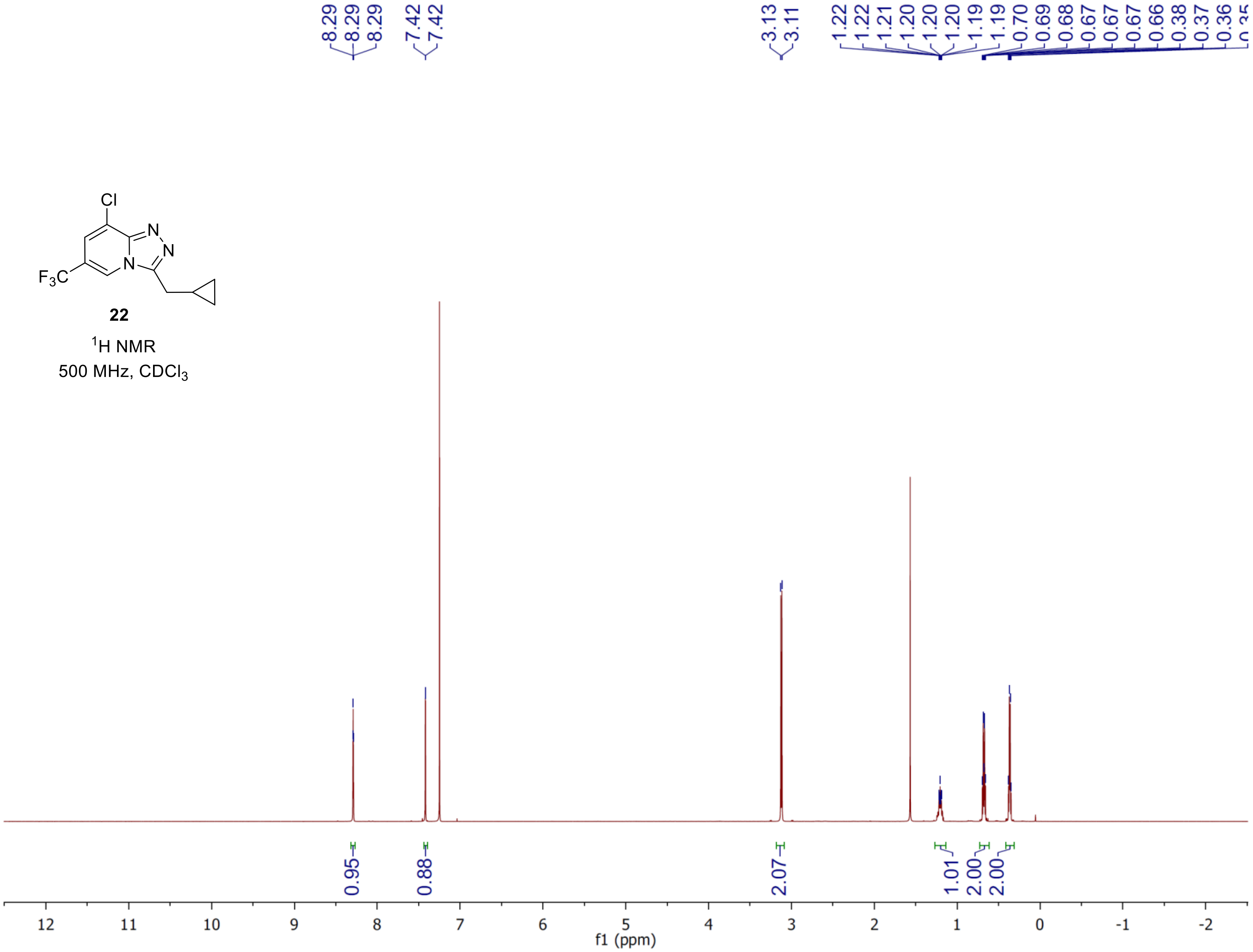

8.29
8.29
8.29
7.42
7.42
3.13
3.11
1.22
1.22
1.21
1.20
1.20
1.20
1.19
1.19
0.70
0.69
0.68
0.67
0.67
0.67
0.66
0.38
0.37
0.36
Cl
N
N
N
F3C
22
1H NMR
500 MHz, CDCl3
0.95
0.88
2.07
1.01
2.00
2.00
12
11
10
9
8
7
6
5
4
3
2
1
0
-1
-2
f1 (ppm)

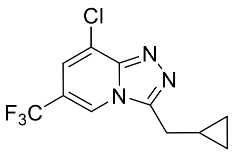


**22**

$^{13}$C NMR

125 MHz, $CDCl_3$

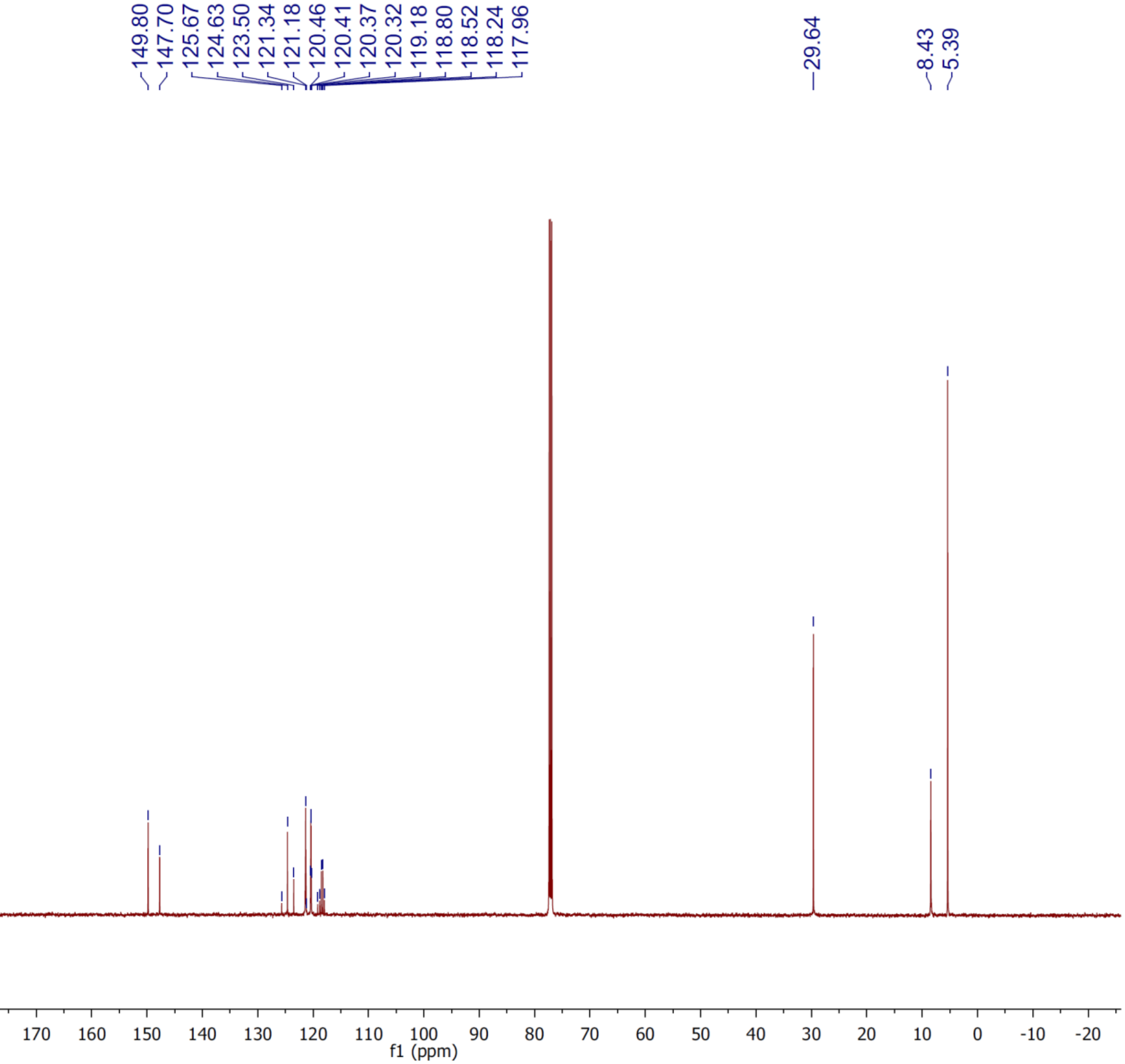

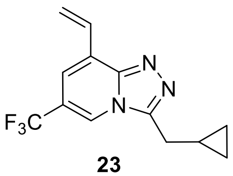


**23**

$^1$H NMR

500 MHz, $CDCl_3$

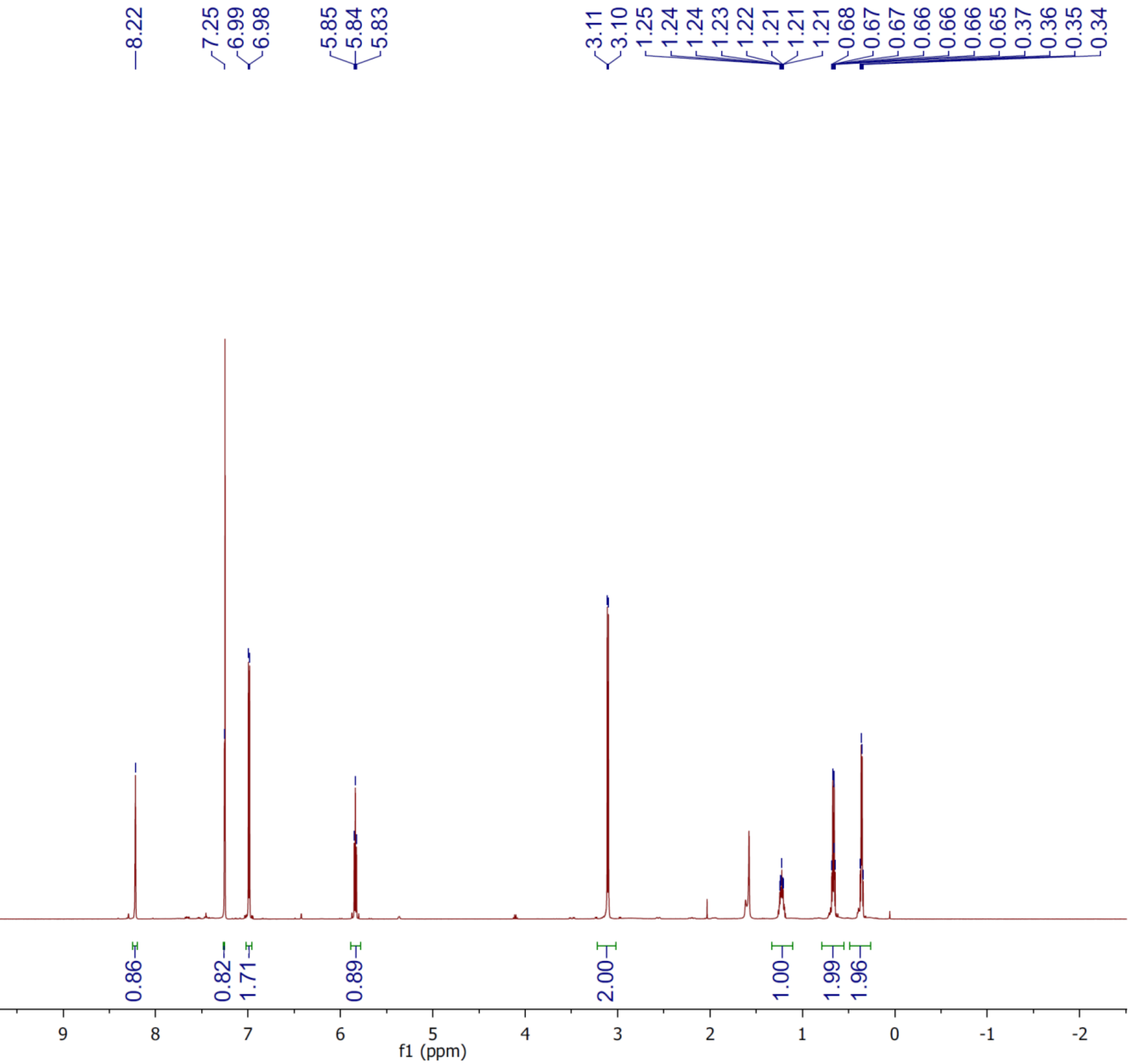

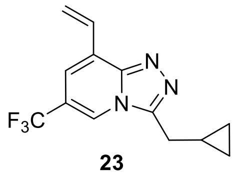

**23**

$^{13}$C NMR

125 MHz, $CDCl_3$

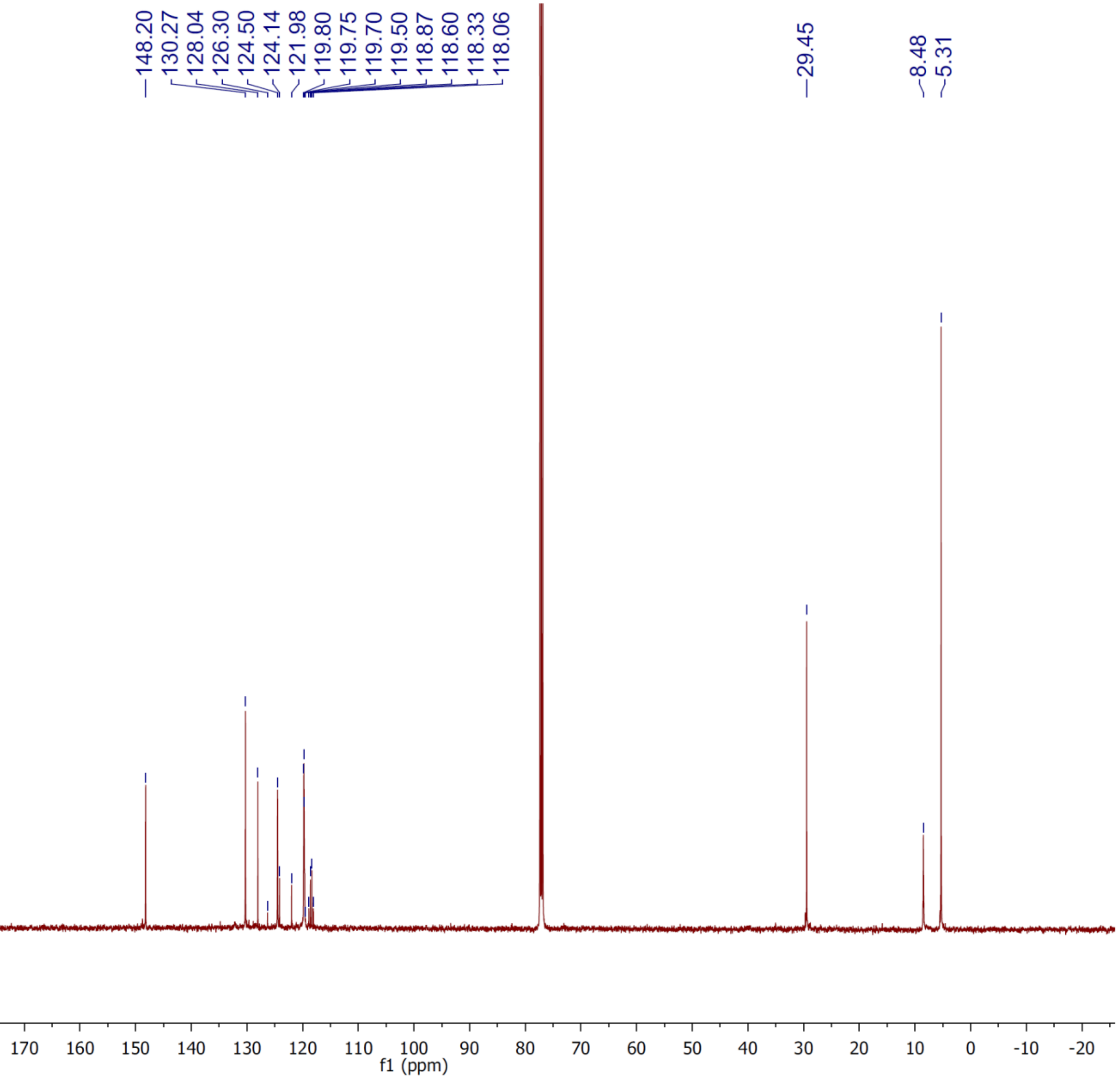

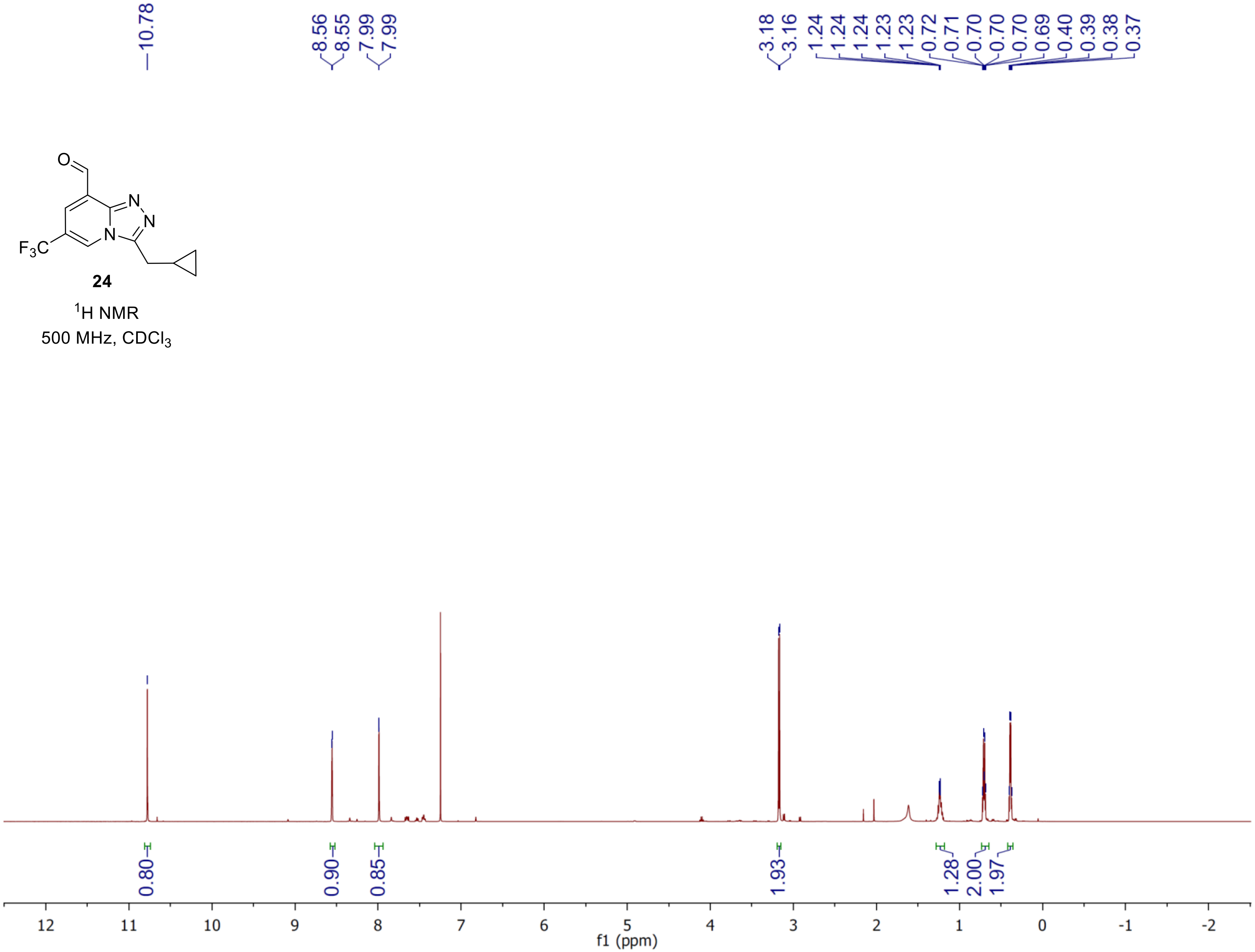
10.78
8.56
8.55
7.99
7.99
3.18
3.16
1.24
1.24
1.24
1.23
1.23
0.72
0.71
0.70
0.70
0.70
0.69
0.40
0.39
0.38
0.37
O
N
N
N
$F_3C$
24
$^1$H NMR
500 MHz, $CDCl_3$
0.80
0.90
0.85
1.93
1.28
2.00
1.97
12
11
10
9
8
7
6
5
4
3
2
1
0
-1
-2
f1 (ppm)

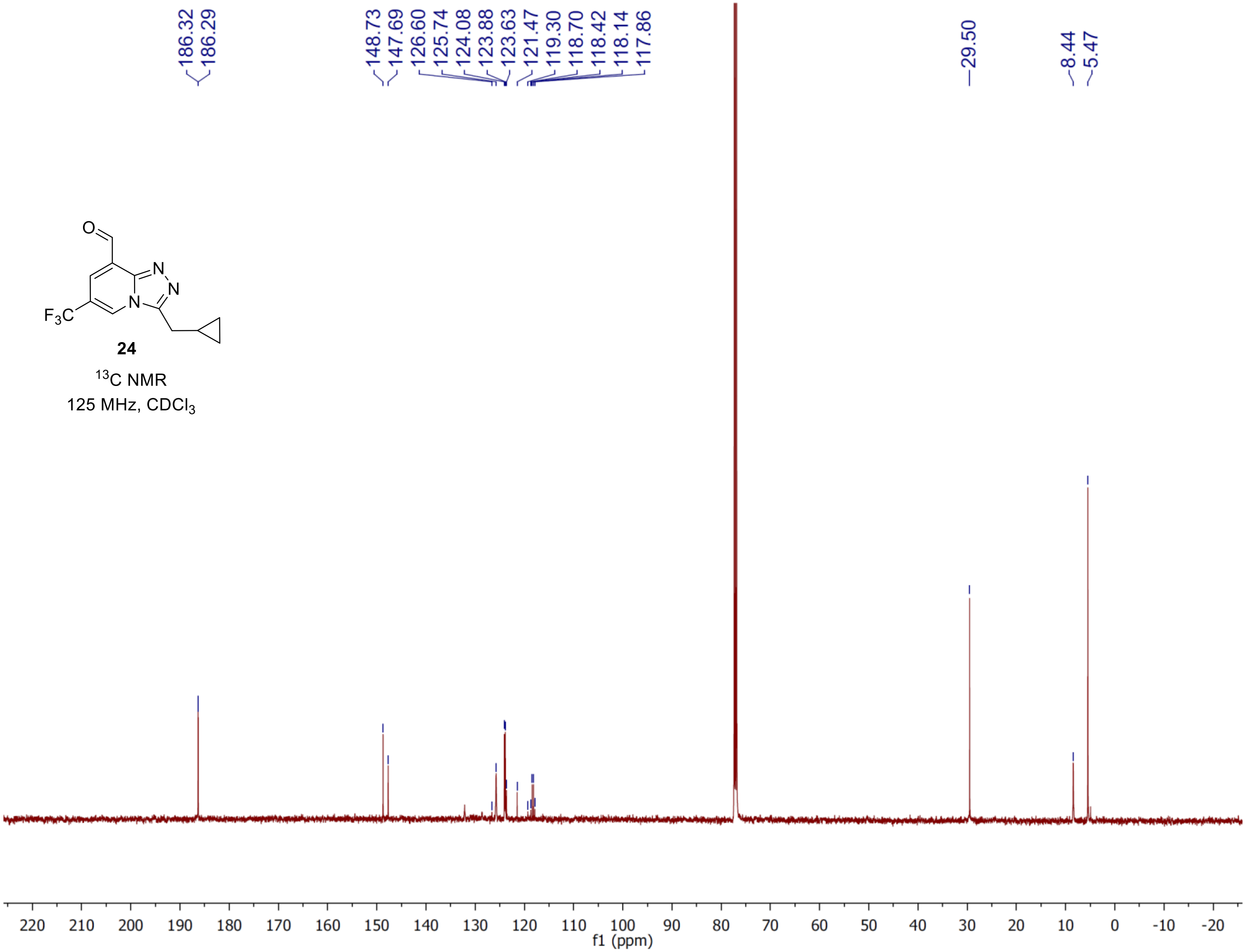



**24**

$^{13}C$ NMR

125 MHz, $CDCl_3$

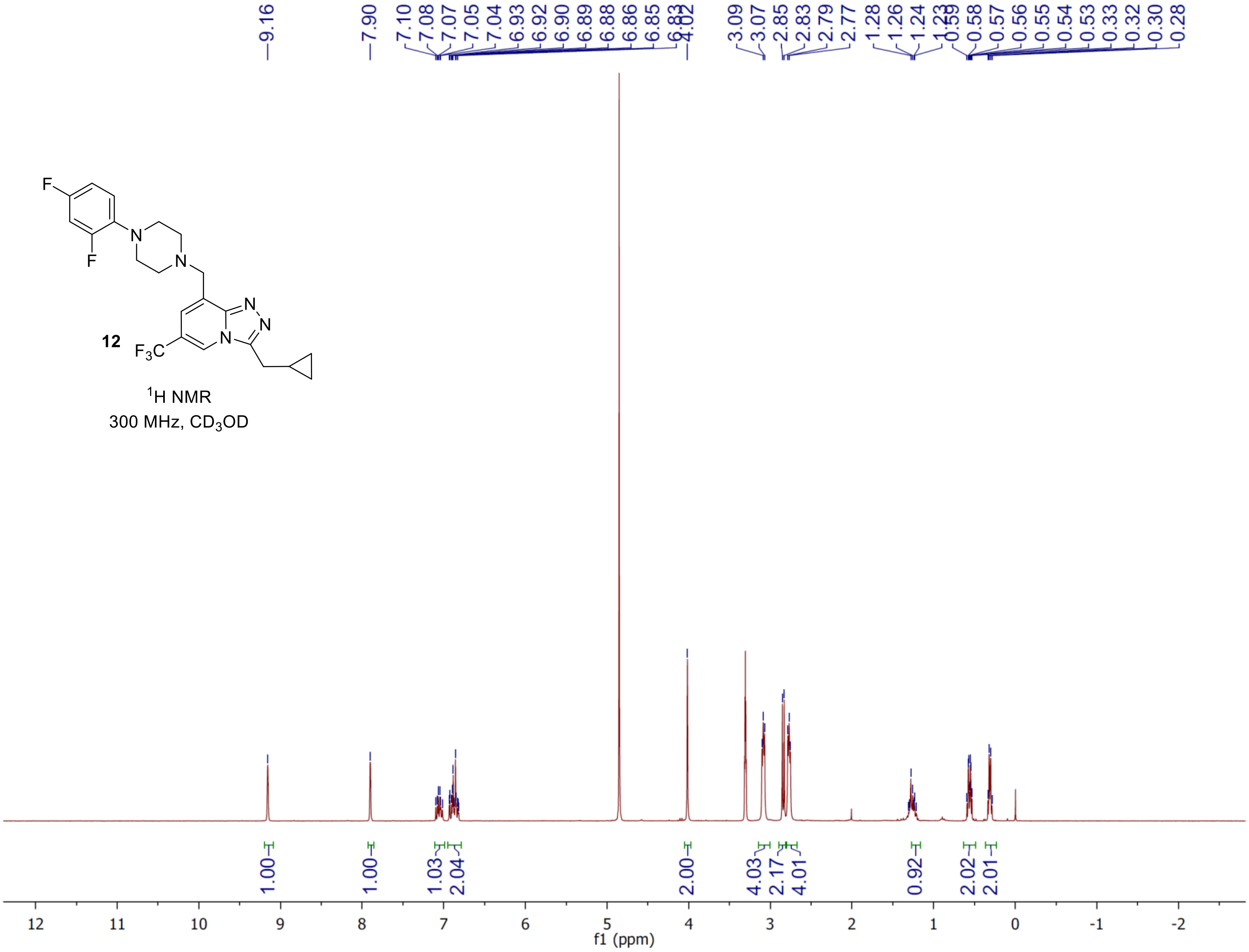
9.16
7.90
7.10
7.08
7.07
7.05
7.04
6.93
6.92
6.90
6.89
6.88
6.86
6.85
6.83
4.02
3.09
3.07
2.85
2.83
2.79
2.77
1.28
1.26
1.24
1.23
0.59
0.58
0.57
0.56
0.55
0.54
0.53
0.33
0.32
0.30
0.28
F
F
N
N
N
N
N
N
$F_3C$
12
$^1$H NMR
300 MHz, $CD_3OD$
1.00
1.00
1.03
2.04
2.00
4.03
2.17
4.01
0.92
2.02
2.01
12 11 10 9 8 7 6 5 4 3 2 1 0 -1 -2
f1 (ppm)

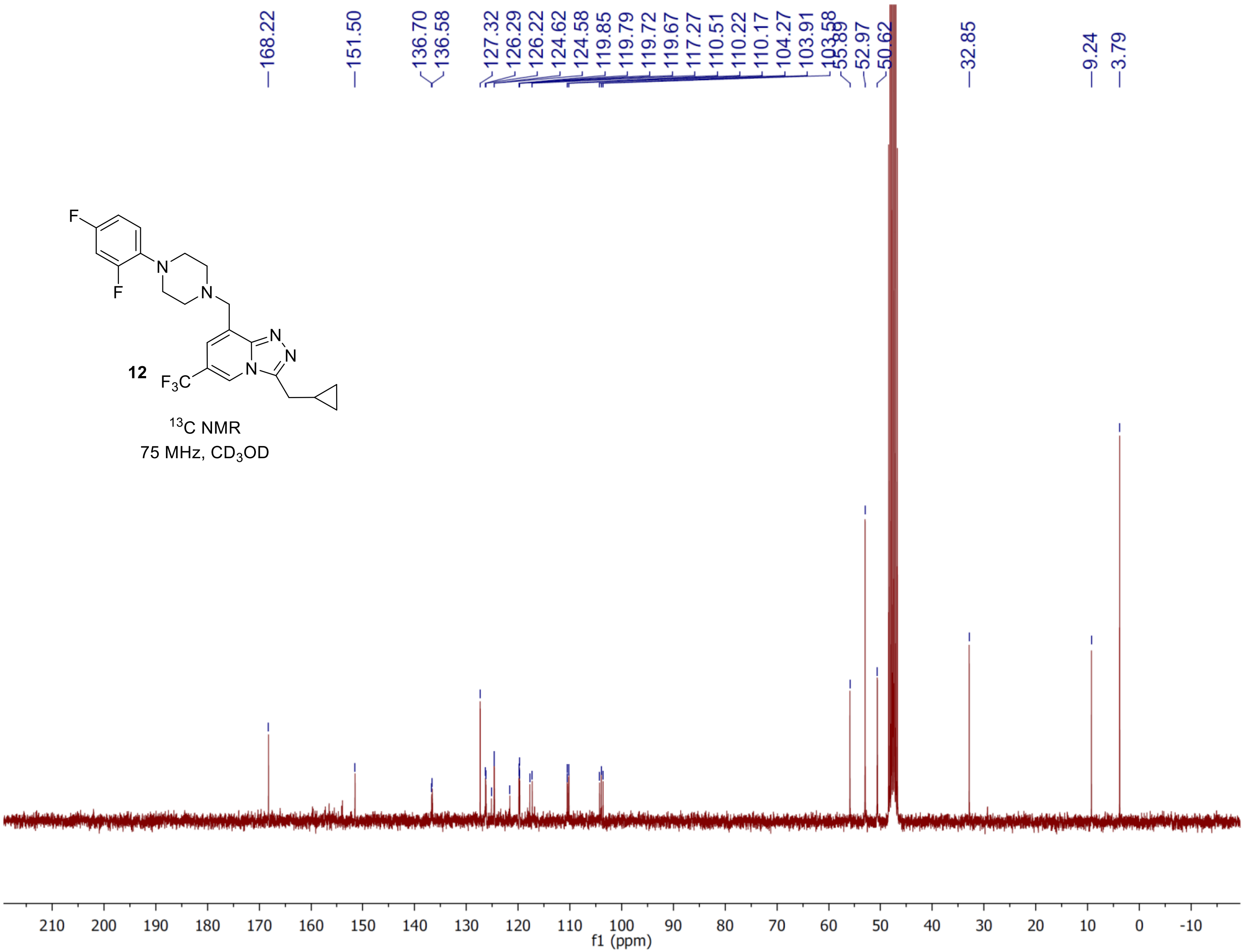
12
$^{13}$C NMR
75 MHz, $CD_3OD$
168.22
151.50
136.70
136.58
127.32
126.29
126.22
124.62
124.58
119.85
119.79
119.72
119.67
117.27
110.51
110.22
110.17
104.27
103.91
103.58
55.89
52.97
50.62
32.85
9.24
3.79
f1 (ppm)

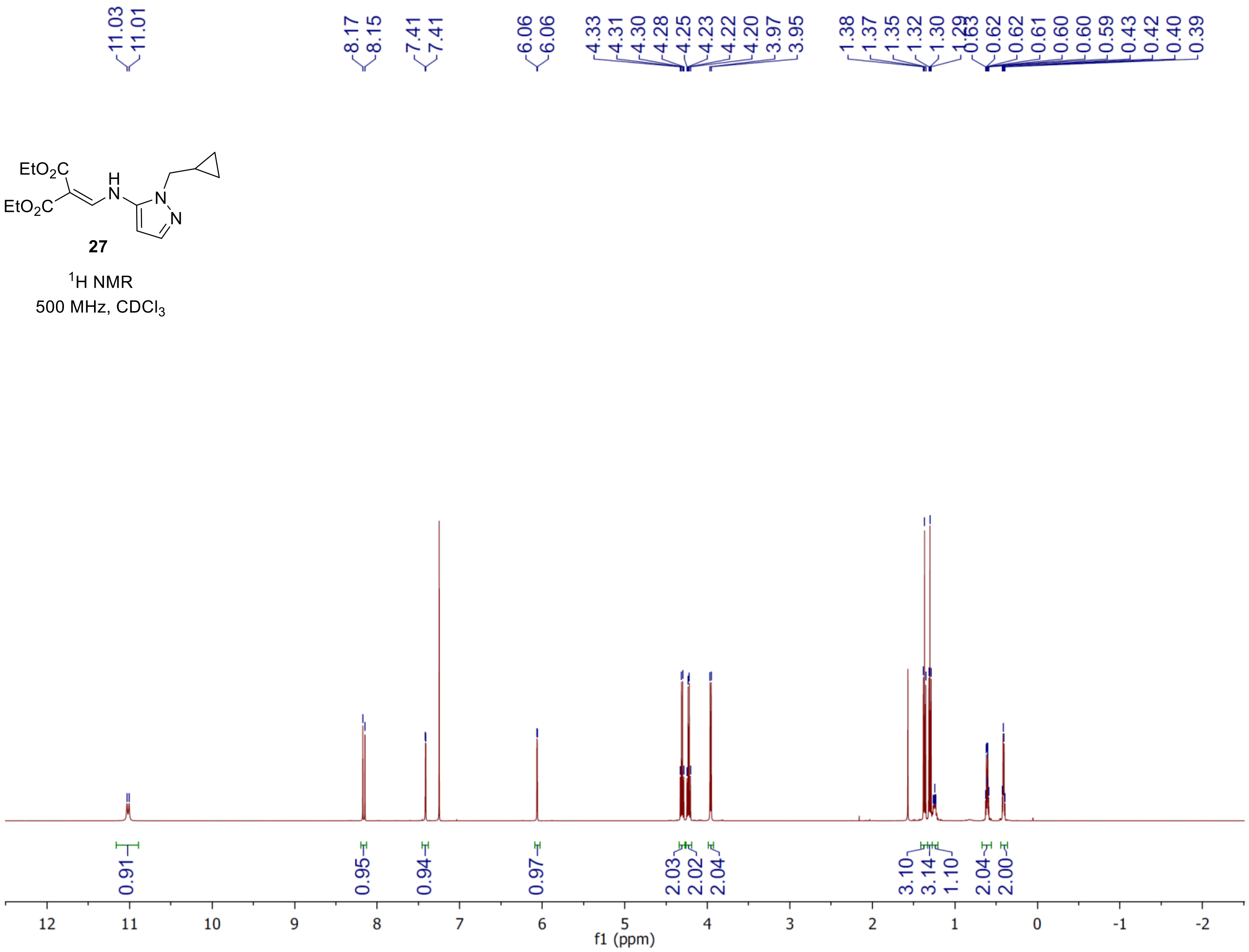
EtO2C
EtO2C
H
N
N
N
27
1H NMR
500 MHz, CDCl3
11.03
11.01
8.17
8.15
7.41
7.41
6.06
6.06
4.33
4.31
4.30
4.28
4.25
4.23
4.22
4.20
3.97
3.95
1.38
1.37
1.35
1.32
1.30
1.29
0.63
0.62
0.62
0.61
0.60
0.60
0.59
0.43
0.42
0.40
0.39
0.91
0.95
0.94
0.97
2.03
2.02
2.04
3.10
3.14
1.10
2.04
2.00
12
11
10
9
8
7
6
5
4
3
2
1
0
-1
-2
f1 (ppm)

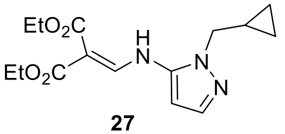


**27**

$^{13}C$ NMR

125 MHz, $CDCl_3$

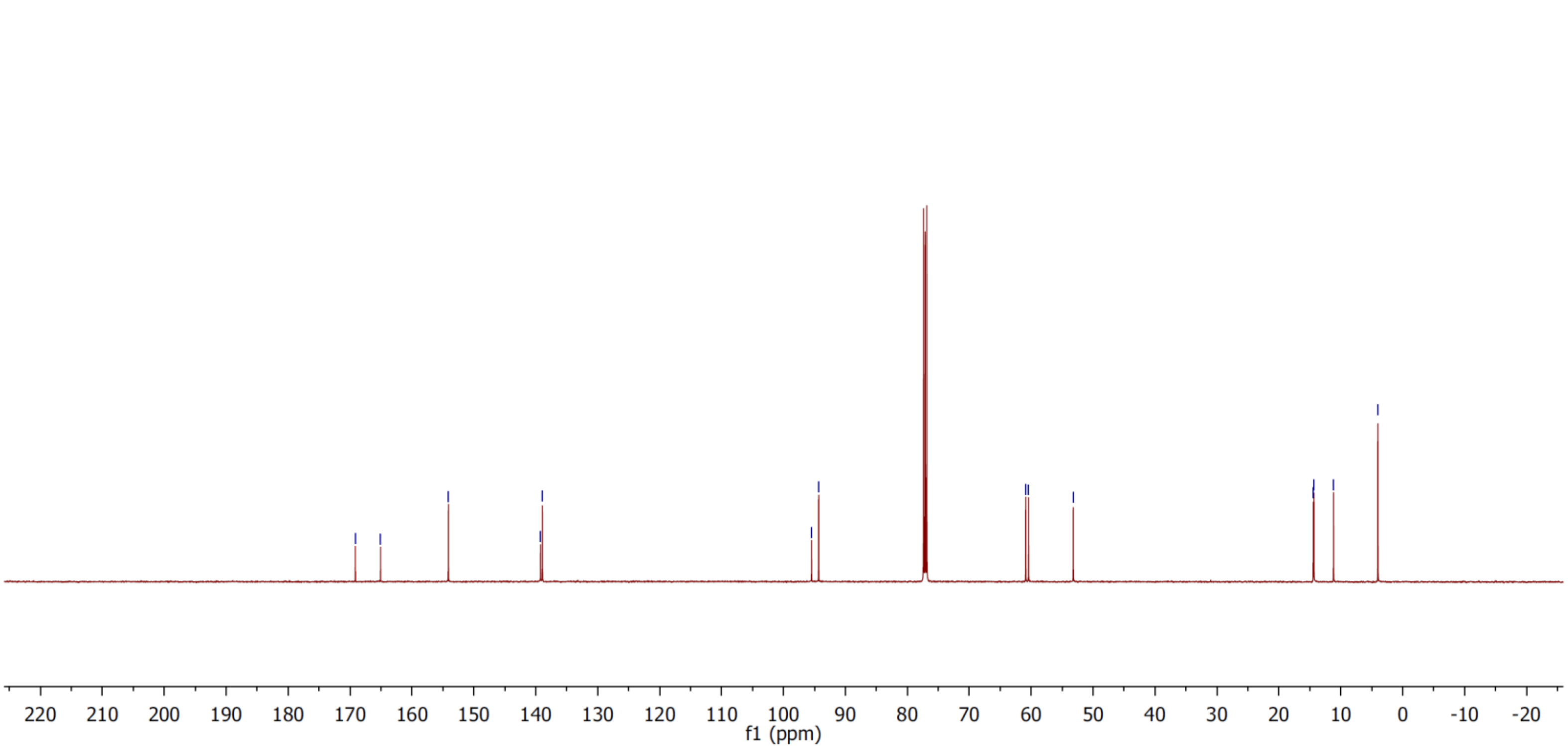

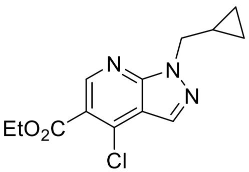


**28**

$^1$H NMR

500 MHz, $CDCl_3$

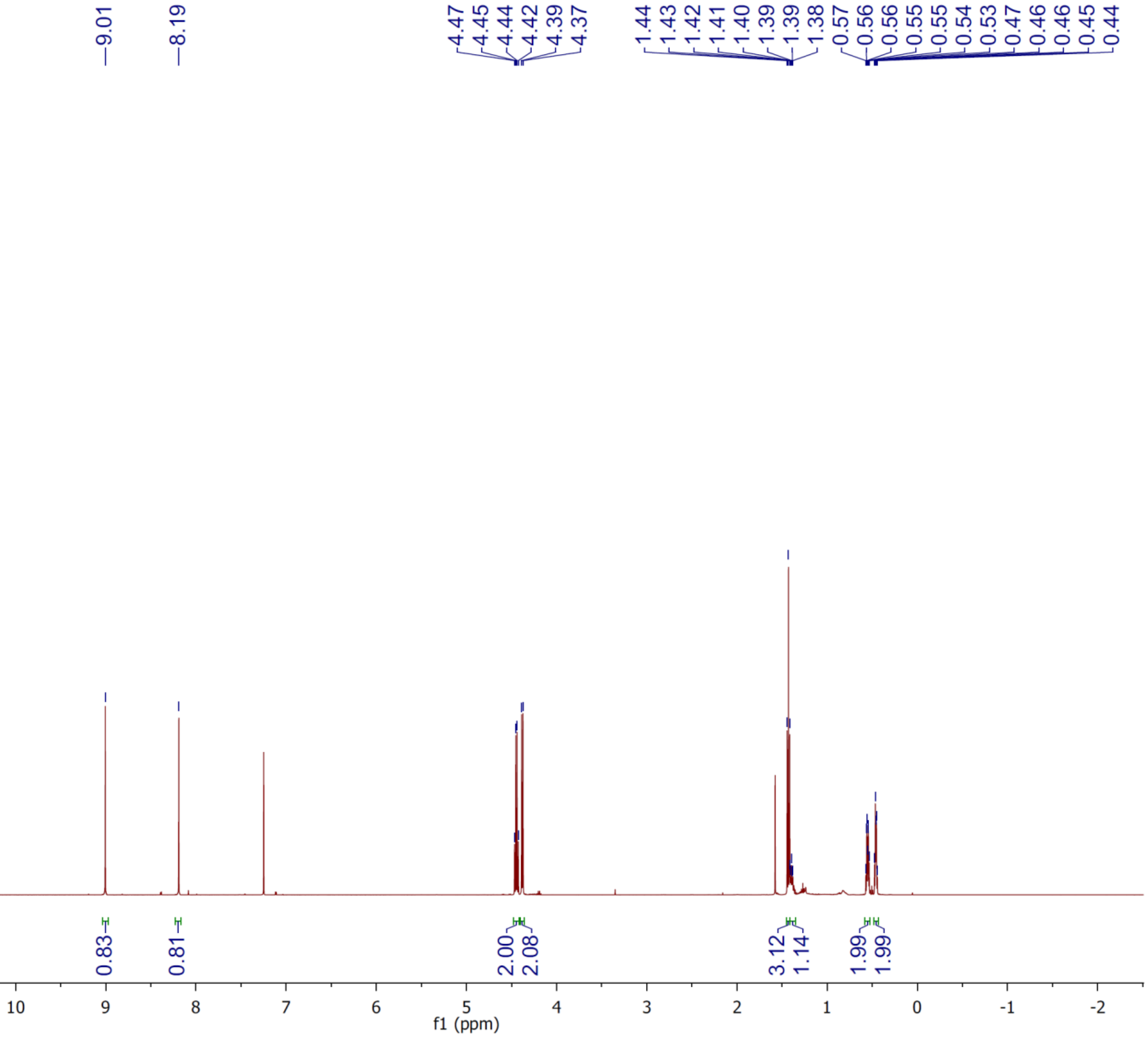

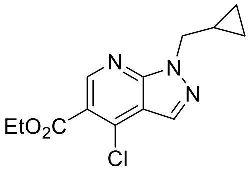

**28**

$^{13}C$ NMR

125 MHz, $CDCl_3$

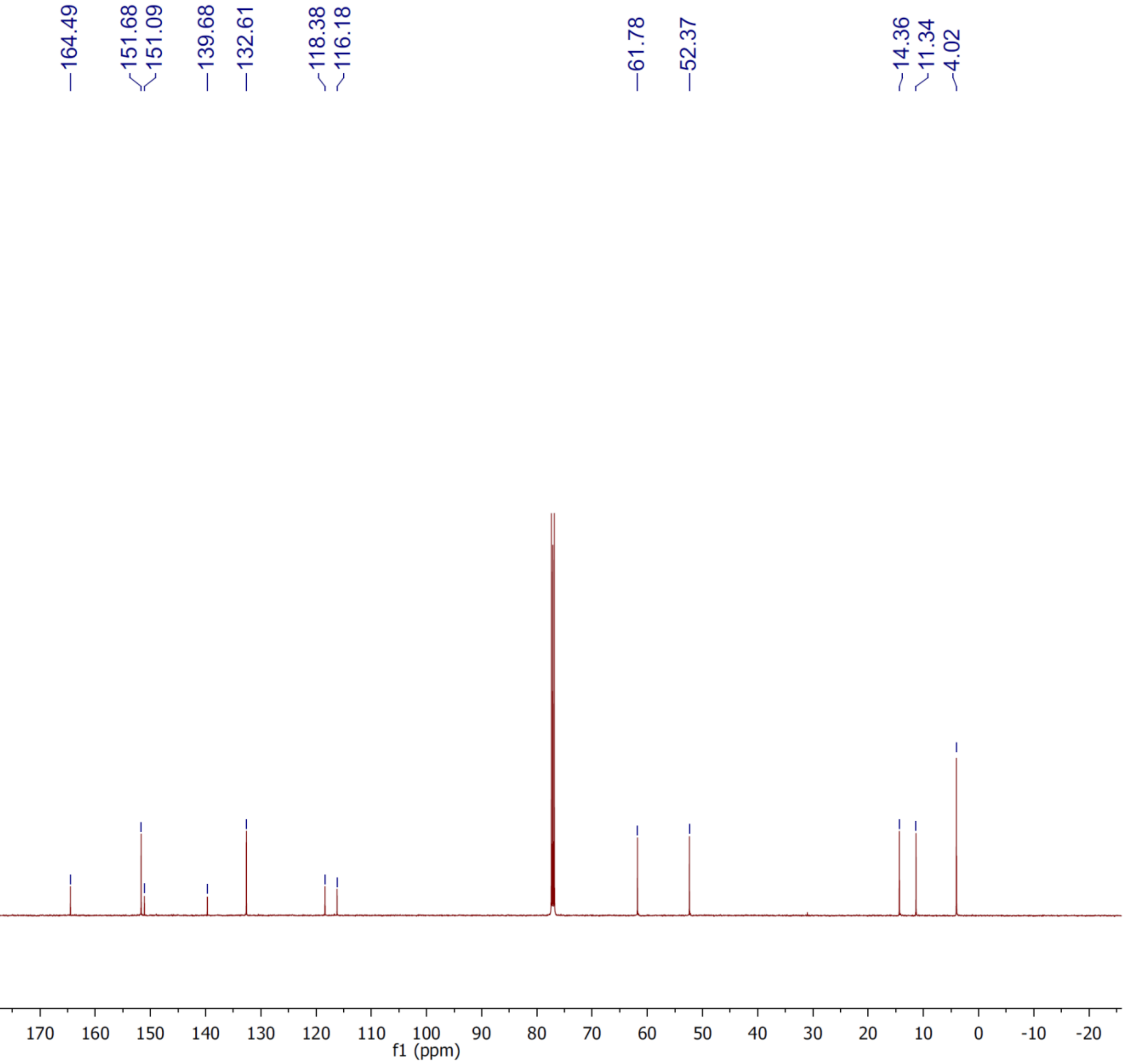

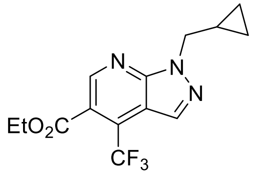


**29**

$^{1}$H NMR

300 MHz, $CDCl_3$

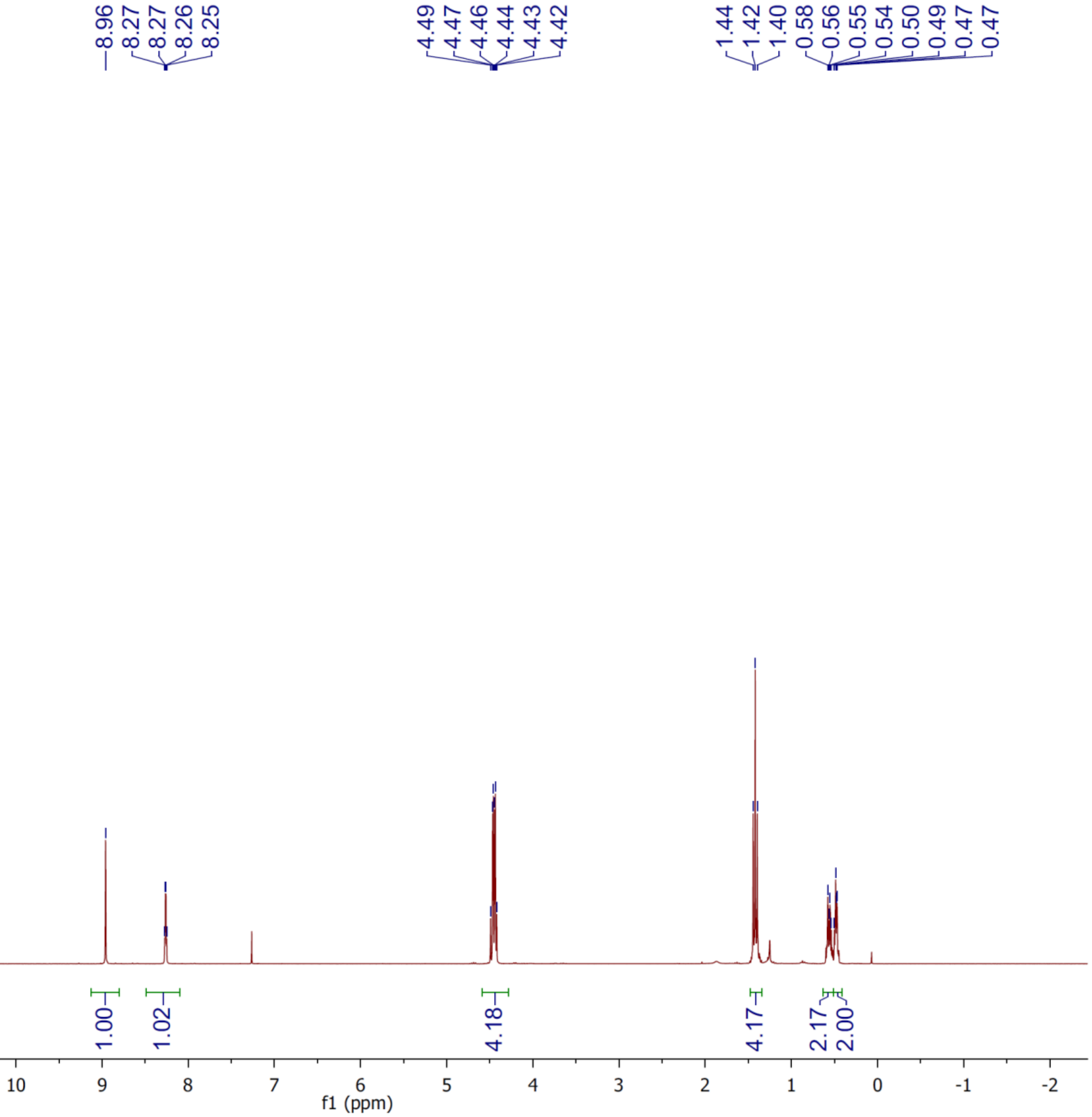

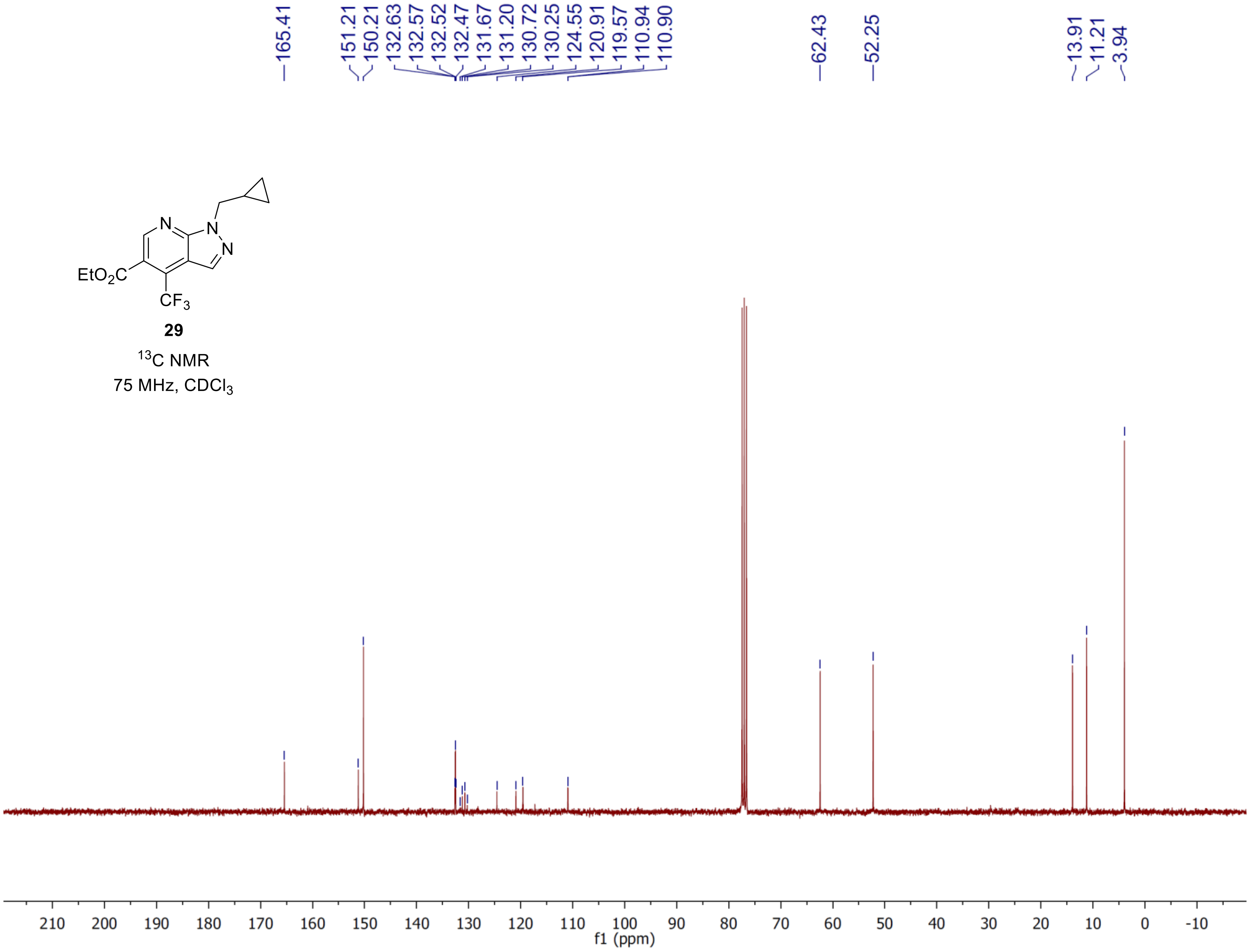

165.41
151.21
150.21
132.63
132.57
132.52
132.47
131.67
131.20
130.72
130.25
124.55
120.91
119.57
110.94
110.90
62.43
52.25
13.91
11.21
3.94
EtO2C
CF3
N
29
13C NMR
75 MHz, CDCl3
f1 (ppm)

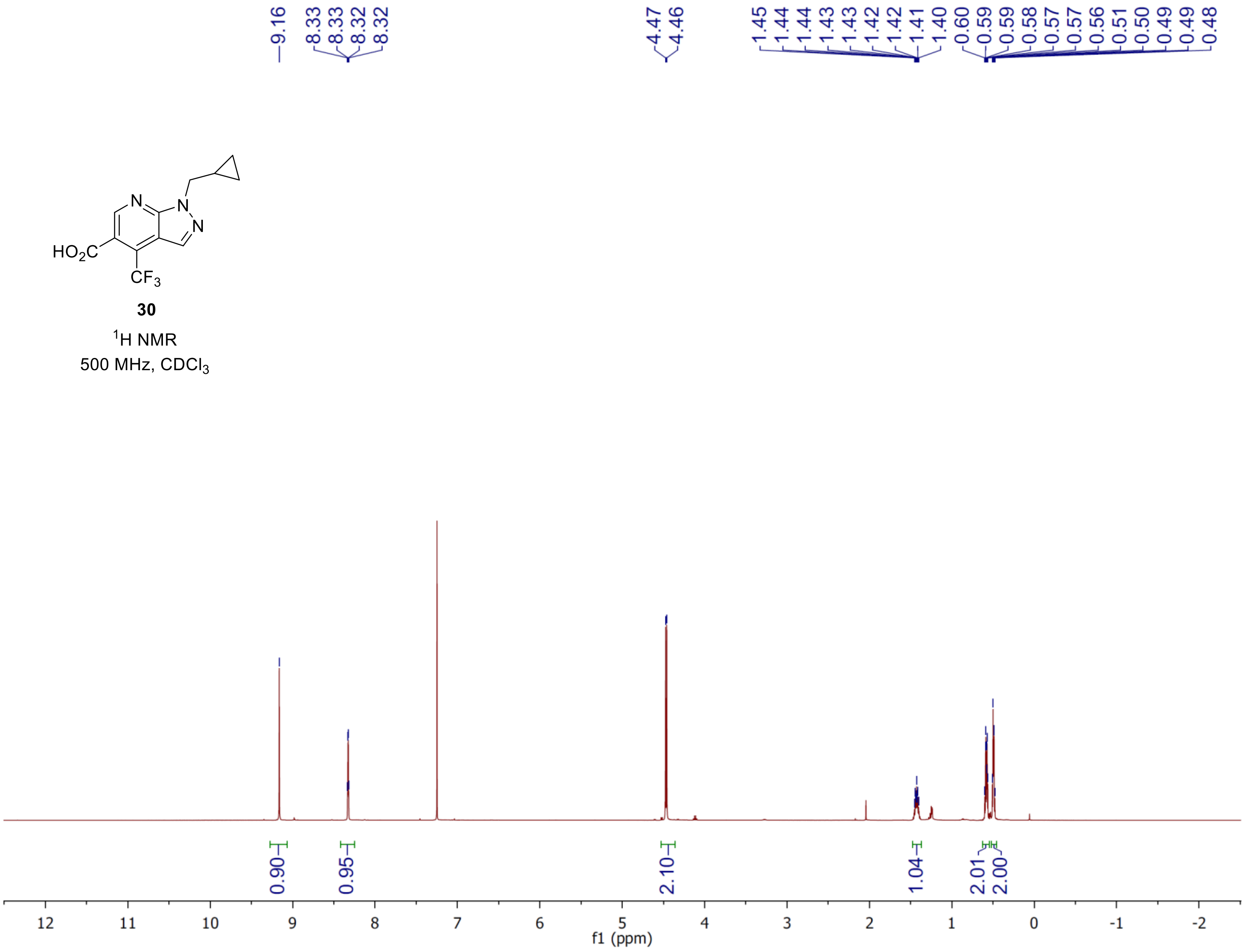
9.16
8.33
8.33
8.32
8.32
4.47
4.46
1.45
1.44
1.44
1.43
1.43
1.42
1.42
1.41
1.40
0.60
0.59
0.59
0.58
0.57
0.57
0.56
0.51
0.50
0.49
0.49
0.48
HO2C
CF3
N
N
N
30
1H NMR
500 MHz, CDCl3
0.90
0.95
2.10
1.04
2.01
2.00
12
11
10
9
8
7
6
5
4
3
2
1
0
-1
-2
f1 (ppm)

**30**

$^{13}C$ NMR

125 MHz, $CD_3OD$

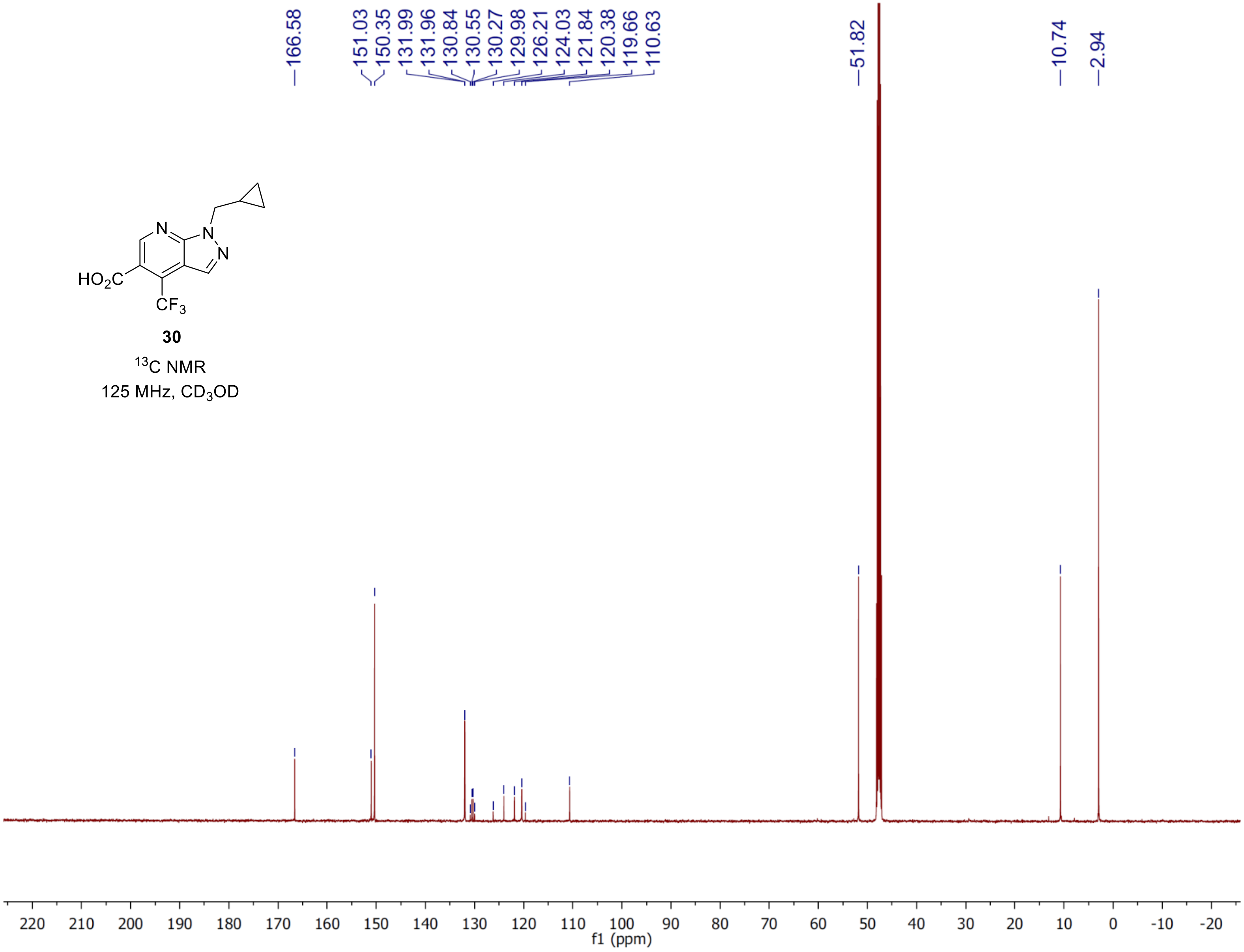

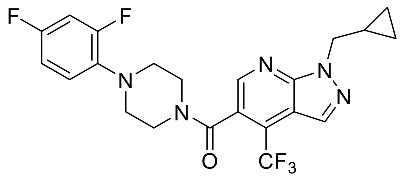


**13**

$^{1}$H NMR

500 MHz, $CDCl_3$

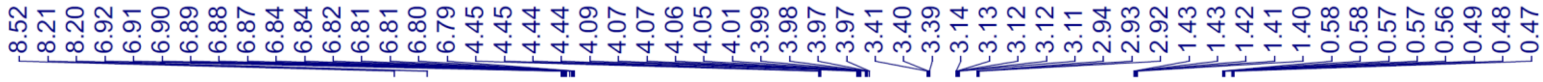


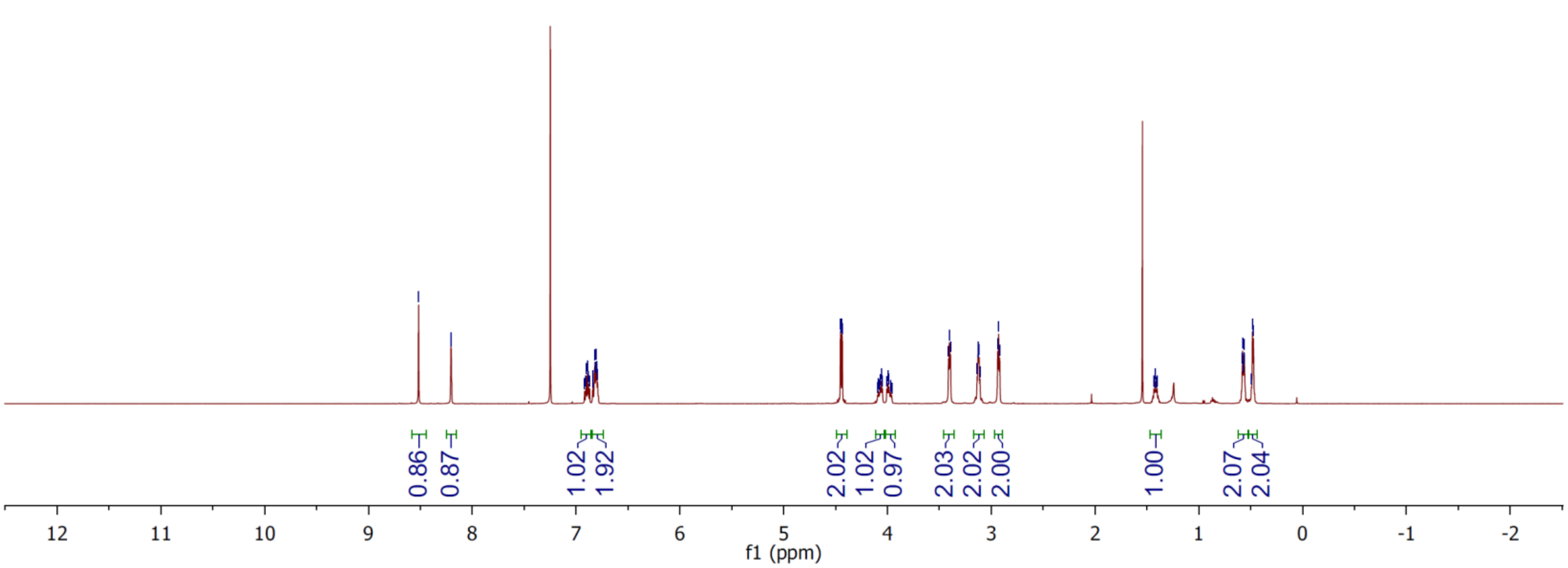

**13**

$^{13}C$ NMR

125 MHz, $CDCl_3$

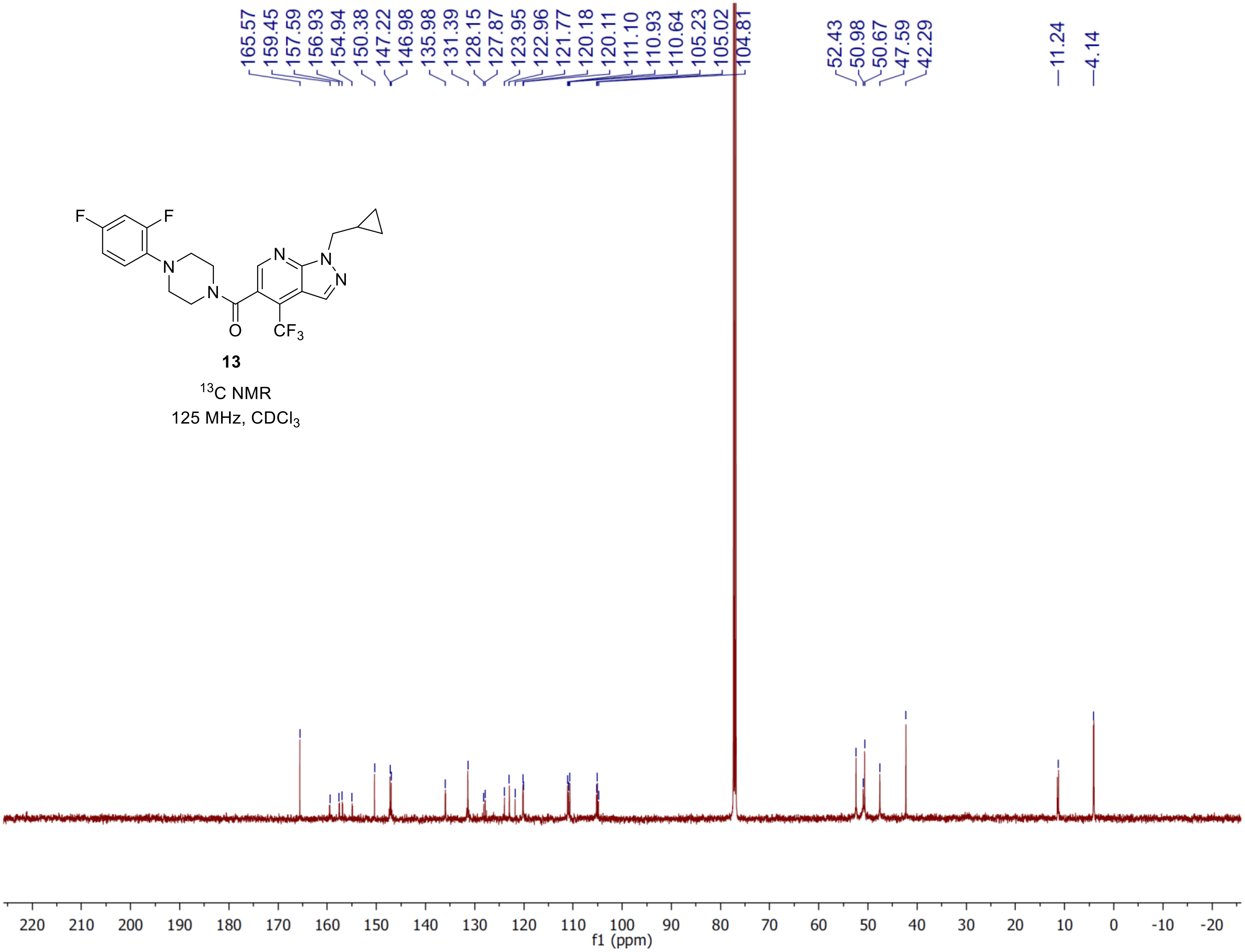

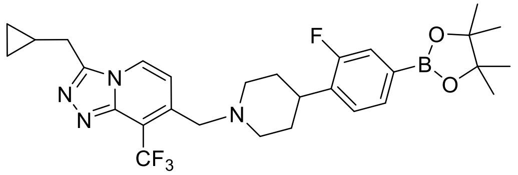

**37**

$^1$H NMR

500 MHz, $CDCl_3$

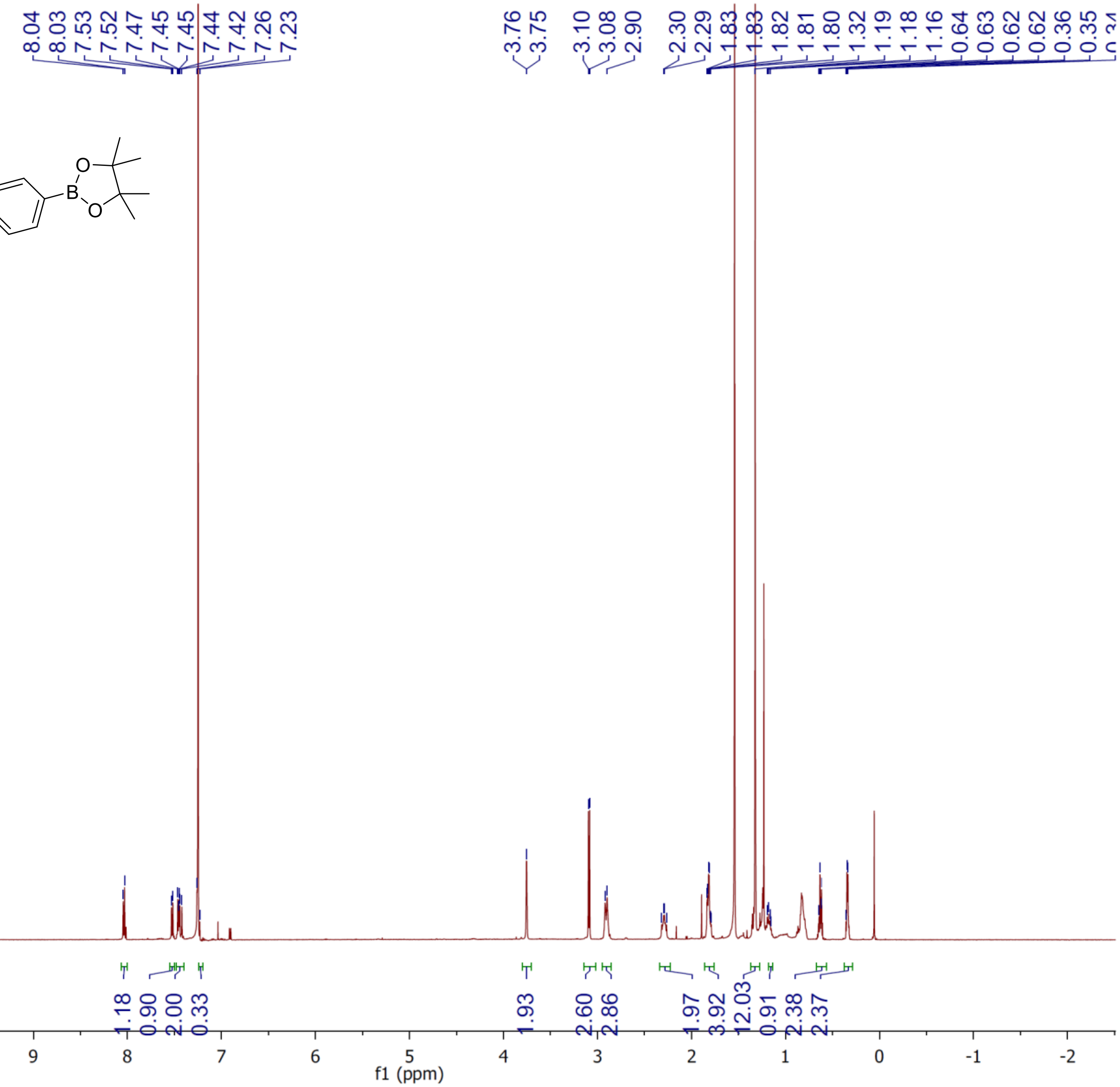

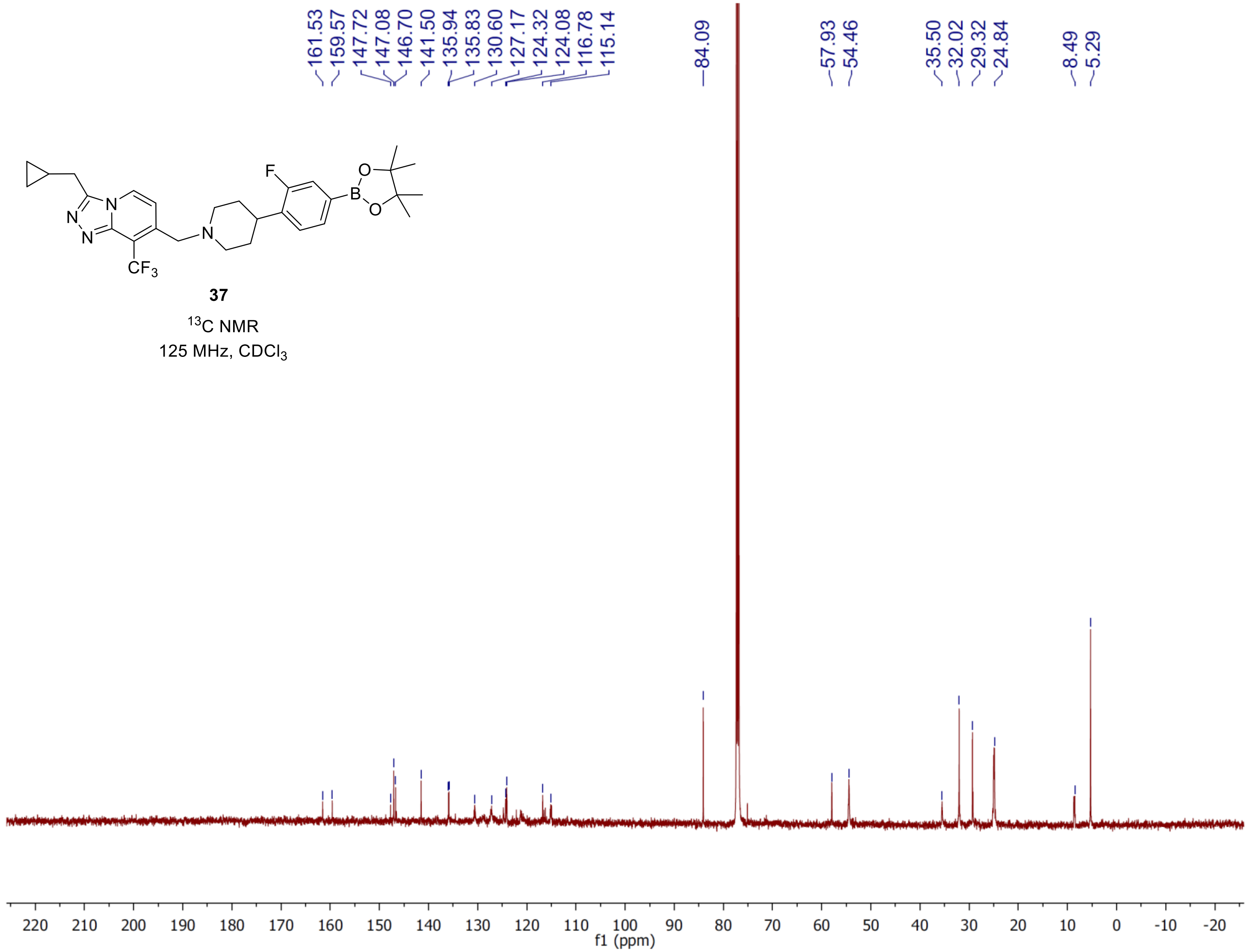

161.53
159.57
147.72
147.08
146.70
141.50
135.94
135.83
130.60
127.17
124.32
124.08
116.78
115.14
84.09
57.93
54.46
35.50
32.02
29.32
24.84
8.49
5.29
F
O
B
O
N
N
N
N
CF3
37
13C NMR
125 MHz, CDCl3
220 210 200 190 180 170 160 150 140 130 120 110 100 90 80 70 60 50 40 30 20 10 0 -10 -20
f1 (ppm)

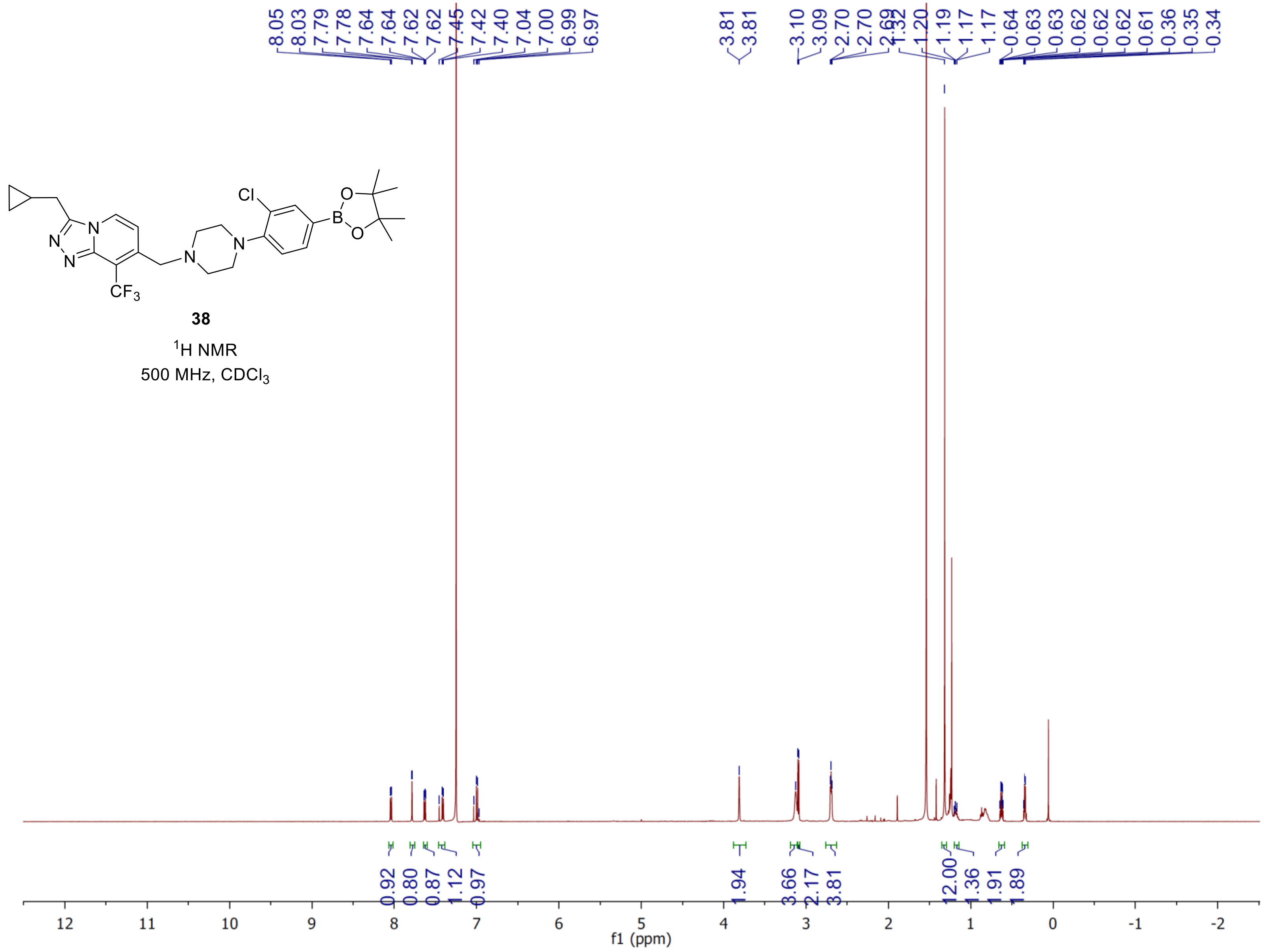


**38**

$^{1}$H NMR

500 MHz, $CDCl_3$

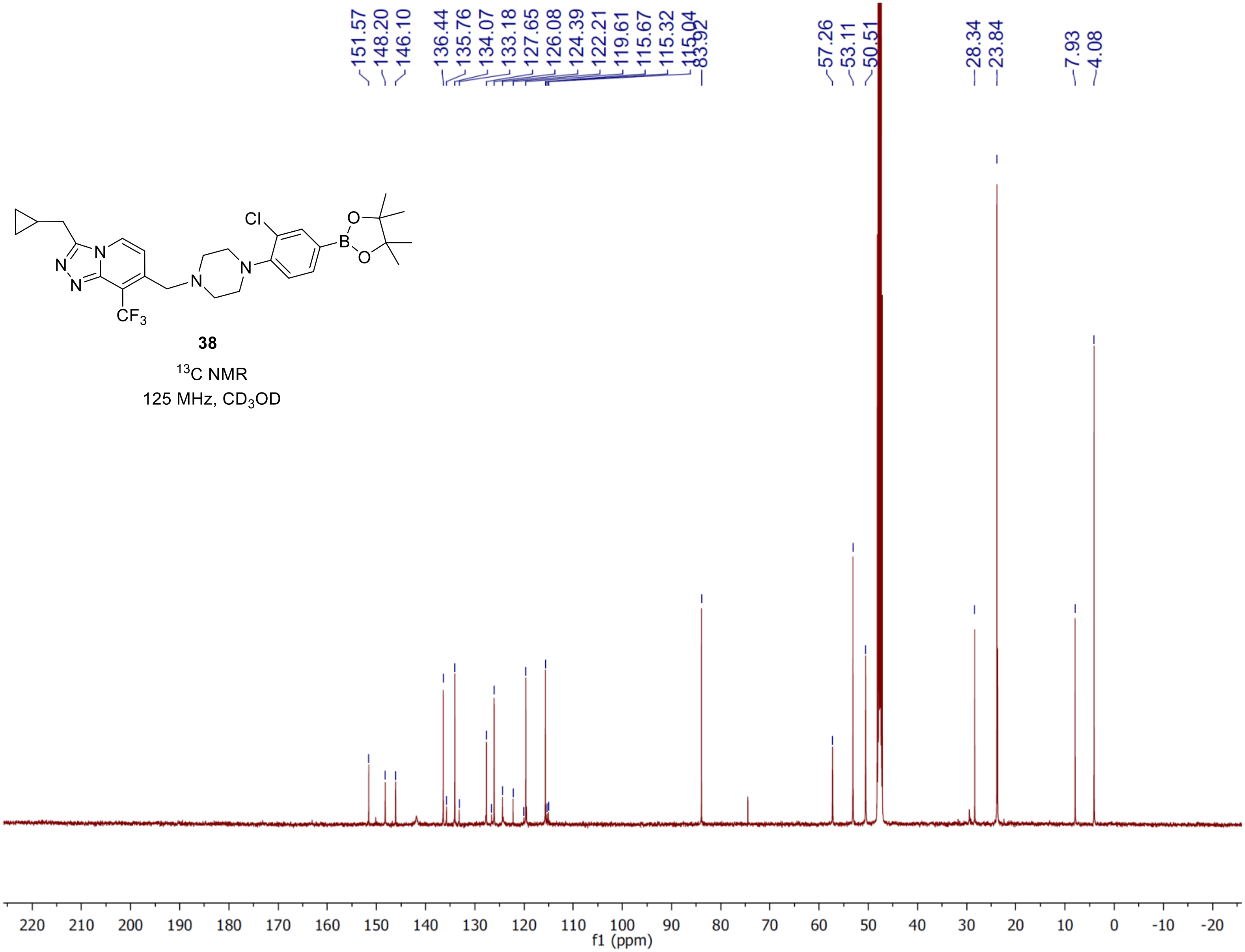
Cl
B
O
O
N
N
N
N
N
CF3
38
13C NMR
125 MHz, CD3OD
151.57
148.20
146.10
136.44
135.76
134.07
133.18
127.65
126.08
124.39
122.21
119.61
115.67
115.32
115.04
83.92
57.26
53.11
50.51
28.34
23.84
7.93
4.08
220 210 200 190 180 170 160 150 140 130 120 110 100 90 80 70 60 50 40 30 20 10 0 -10 -20
f1 (ppm)
